# DDI2 protease activity controls embryonic development and inflammation via TCF11/NRF1

**DOI:** 10.1101/2020.12.16.423023

**Authors:** Monika Siva, Stefanie Haberecht-Müller, Michaela Prochazkova, Jan Prochazka, Frantisek Sedlak, Kallayanee Chawengsaksophak, Petr Kasparek, Radislav Sedlacek, Jan Konvalinka, Elke Krüger, Klara Grantz Saskova

## Abstract

DDI2 is an aspartic protease that cleaves polyubiquitinated substrates. Upon proteotoxic stress, DDI2 activates the ER-bound transcription factor TCF11/NRF1 (NFE2L1), a master regulator of proteostasis maintenance in mammalian cells, and ensures the expression of rescue factors including proteasome subunits. Here we describe the consequences of DDI2 ablation both *in vivo* and in cells. Knock-out of DDI2 in mice resulted in embryonic lethality at E12.5 with severe developmental failure. Molecular characterization of the embryos and surrogate *DDI2* knock-out cell lines showed insufficient proteasome expression with proteotoxic stress, accumulation of high molecular weight ubiquitin conjugates, and induction of the unfolded protein and integrated stress responses. We also show that *DDI2* KO-induced proteotoxic stress causes the cell-autonomous innate immune system to induce a type I interferon signature. These results indicate an important role for DDI2 in the proteostasis network of cells and tissues and in the maintenance of a balanced immune response.

**Highlights:**

- DDI2-deficiency in mice causes severe developmental failure and embryonic lethality at mid-late gestation
- DDI2-deficiency causes severe proteotoxic stress and proteasome impairment
- DDI2-deficiency induces the UPR and ISR signaling pathways
- DDI2-deficient cells survive via STAT3-dependent interferon signaling

## INTRODUCTION

The ubiquitin proteasome system (UPS) with the proteasome as the major proteolytic enzyme complex ensures the degradation of most intracellular proteins. Regulatory proteins as well as proteins that are misfolded and/or oxidatively damaged are tagged with ubiquitin chains, making them targets for destruction by the proteasome. The proteasome itself is composed of the 20S proteolytic core complex containing the active site β-subunits and axially attached regulatory complexes that can bind, deconjugate, unfold and translocate ubiquitin-modified substrates into the proteolytic 20S complex (Collins and Goldberg, 2017). By controlling the intracellular pool of regulators, proteasomes actively participate in the regulation of myriad signaling pathways, including mTOR, the unfolded protein response (UPR), the integrated stress response (ISR), and both innate and adaptive immune responses (Ling et al., 2012; Xu et al., 2015; Yu and Hayward, 2010; Zhao et al., 2016; Zhao and Goldberg, 2016). By maintaining protein homeostasis (proteostasis), the UPS also helps to maintain cell integrity, viability and function. Proteotoxic stress from various sources can be counteracted by the shutdown of global protein translation, or by upregulation of protein quality control and degradation machineries including stress-specific sets of ubiquitin-conjugation and deconjugation factors as well as alternative proteasome isoforms. Thus, the ISR and UPR pathways act as central hubs integrating cellular responses to proteotoxic stress in order to determine cell fate (Hetz, 2012; Hetz et al., 2020; Hipp et al., 2014; Pakos-Zebrucka et al., 2016). Interestingly, the ISR and UPR have been reported to play regulatory roles beyond protein quality control with important implications for inflammation and metabolism. Central to this mechanism is the phosphorylation of eIF2α by the endoplasmic reticulum (ER) stress sensor PERK and/or cytoplasmic stress sensors (such as PKR and GCN2), which results in repression of protein translation to decrease the protein load. In addition, regulated IRE1-dependent decay (RIDD) contributes to this mechanism through the degradation of diverse mRNAs (Maurel et al., 2014; So et al., 2012).

The NGLY1-p97/VCP-TCF11/NRF1-DDI2 axis represents another important rescue mechanism to combat proteotoxic stress with implications for the treatment of cancer using proteasome inhibitors (Fassmannová et al., 2020; Northrop et al., 2020; Radhakrishnan et al., 2010). The transcription factor TCF11/NRF1 encoded by the *NFE2L1* (NFE2-related factor 1) gene has been identified as the key regulator for proteasome formation and induces a time and concentration-dependent adaptive response (Meiners et al., 2003; Radhakrishnan et al., 2010; Steffen et al., 2010). Under non-stressed conditions the ER-tethered TCF11/NRF1 is permanently targeted for ER-associated degradation (ERAD) involving the E3 ubiquitin ligase HRD1 and the AAA-ATPase p97/VCP. In response to proteotoxic stress, TCF11/NRF1 is extracted by p97/VCP, deglycosylated by NGLY1, and subsequently cleaved by the aspartyl-protease DDI2, enabling its nuclear translocation (Koizumi et al., 2016; Siva et al., 2016; Tomlin et al., 2017). This is crucial for the induction of rescue factors including new proteasome subunits and other UPS-related factors (Radhakrishnan et al., 2010). Calpain-1 has been shown to be involved in the degradation of membranebound TCF11/NRF1 under non-inducing conditions. DDI2 in turn delays TCF11/NRF1 degradation, probably by promoting the activation of TCF11/NRF1 rather than its degradation pathway and protecting TCF11/NRF1 from ERAD (Northrop et al., 2020; Nowak et al., 2018). In addition, NRF1, the mouse ortholog of TCF11/NRF1, has been shown to be involved in cholesterol and fat metabolism (Bartelt et al., 2018; Widenmaier et al., 2017).

Pioneering recent work shows that activation of the TCF11/NRF1 transcriptional pathways by proteotoxic stress during proteasome impairment or oxidative stress is accompanied by induction of typical UPR and ISR downstream events such as inactivation of translation by phosphorylation of eIF2α, induced transcription of ER chaperones, and processing of XBP-1 (Sotzny et al., 2016; Studencka-Turski et al., 2019). Upregulation of type I interferon (IFN) induction has also been observed both in hematopoietic and non-hematopoietic cells in response to proteasome impairment by inhibitors as well as in patients with rare proteasome loss of function mutations. Thus, cellular responses to proteasome impairment include activation of TCF11/NRF1, activation of the UPR/ISR, and initiation of an IFN response (Brehm et al., 2015; Poli et al., 2018; Sotzny et al., 2016; Studencka-Turski et al., 2019).

Recently, a homozygous deep intronic mutation in the *PSMC3* gene, which encodes the proteasome ATPase subunit RPT5, was shown to induce proteotoxic stress in the cells of a patient, although the proteasome content and activity in the patient’s fibroblasts was unaffected. The TCF11/NRF1 transcriptional pathway was unable to promote proteasome recovery resulting in a fulminant proteotoxic crisis (Kroll-Hermi et al., 2020).

DDI2 belongs to the family of Ddi1-like proteins. These are proteasomal shuttles which contain a retroviral protease-like domain (RVP) strikingly reminiscent of HIV protease. The structure of the DDI2 RVP confirmed the conserved fold of the active site, with flexible flaps covering a large active-site cavity, and suggested that DDI2 RVP cleaves bulky substrates (Siva et al., 2016). The N-terminal ubiquitin-like (UBL) domain and the ubiquitin interaction motif (UIM) at the C-terminus of the structure were shown to weakly but specifically bind ubiquitin (Siva et al., 2016). Furthermore, the helical domain of DDI2 (HDD) could also be involved in substrate recognition (Siva et al., 2016; Trempe et al., 2016). This hypothesis is supported by the observation that Ddi1p and DDI2 are required for removal of RPT2 from stalled replisomes and that HDD from yeast Ddi1 is involved in replication stress (Serbyn et al., 2020; Svoboda et al., 2019). Very recently, yeast Ddi1 and mammalian DDI2 proteins have been shown to be polyubiquit-independent endoproteases (Dirac-Svejstrup et al., 2020; Yip et al., 2020) with TCF11/NRF1 and NRF3 as specific substrates (Chowdhury et al., 2017; Koizumi et al., 2016).

Although some aspects of mammalian DDI2 functions have been studied on the cellular level, little is known about the physiological role and functional complexity of DDI2 *in vivo*. Applying conditions of DDI2 deficiency to cells and mouse models, here we provide evidence that DDI2 is required for physiological growth and development as well as for maintenance of proteostasis. Using mutations which target the proteolytic domain, we show that proteolytic activity is the dominant function of DDI2 during embryonic development. We also confirm that its absence causes severe proteotoxic stress with accumulation of high molecular weight ubiquitin conjugates and impairment of TCF11/NRF1 activation. The impaired capacity of DDI2-deficient cells to compensate for *de novo* proteasome formation results in activation of UPR and ISR and ultimately promotes the induction of type I IFN stimulated genes.

## RESULTS

### DDI2 deficiency in mice results in mid-gestation embryonic lethality

With the goal of understanding the biological role of DDI2 in a complex system, we generated two mouse strains in which DDI2 function was switched off by either knocking out *Ddi2* or replacing it with a *Ddi2* protease-defective allele. The full knock-out strain *C57BL/6NCrl-Ddi2^tm1b(EUCOMM)Hmgu^/P*h, in text referred as *Ddi2^KO^* (genotypes *Ddi2^+/+^, Ddi2^+/-^* and *Ddi2^-/-^*), lacks the critical exon 2 (with an additional polyA insertion instead) and also contains a *LacZ* reporter gene. The second strain was designed to disrupt two critical functions of the DDI2 protease domain – catalytic activity and dimerization. This *C57Bl/6NCrl-Ddi2^em1^/Ph* strain, in text further referred as *Ddi2^PD^* (genotypes *Ddi2^ex6+/+^, Ddi2^ex6+/-^* and *Ddi2^ex6-/-^*), was generated by TALEN-mediated excision of exon 6 of the *Ddi2* gene in a *C57Bl/6NCrl* background, resulting in ablation of the DDI2 protease domain (Δ254-296) (Figure 1A). Genomic DNA of *Ddi2^PD^* F1-generation animals used to establish colonies was screened for off-target modifications at the 12 most likely sites on chromosome 4 with no off-target modifications detected. Systemic phenotyping screens of adult mice of both *Ddi2*-altered strains performed on animals at the age of 9-16 weeks showed that heterozygous *Ddi2^+/-^* and *Ddi2^ex6+/-^* mice are viable, fertile and without a clear distinct phenotype as adults. In both strains, however, homozygous mutants exhibited embryonic lethality in the mid-gestation period.

**Figure 1:**
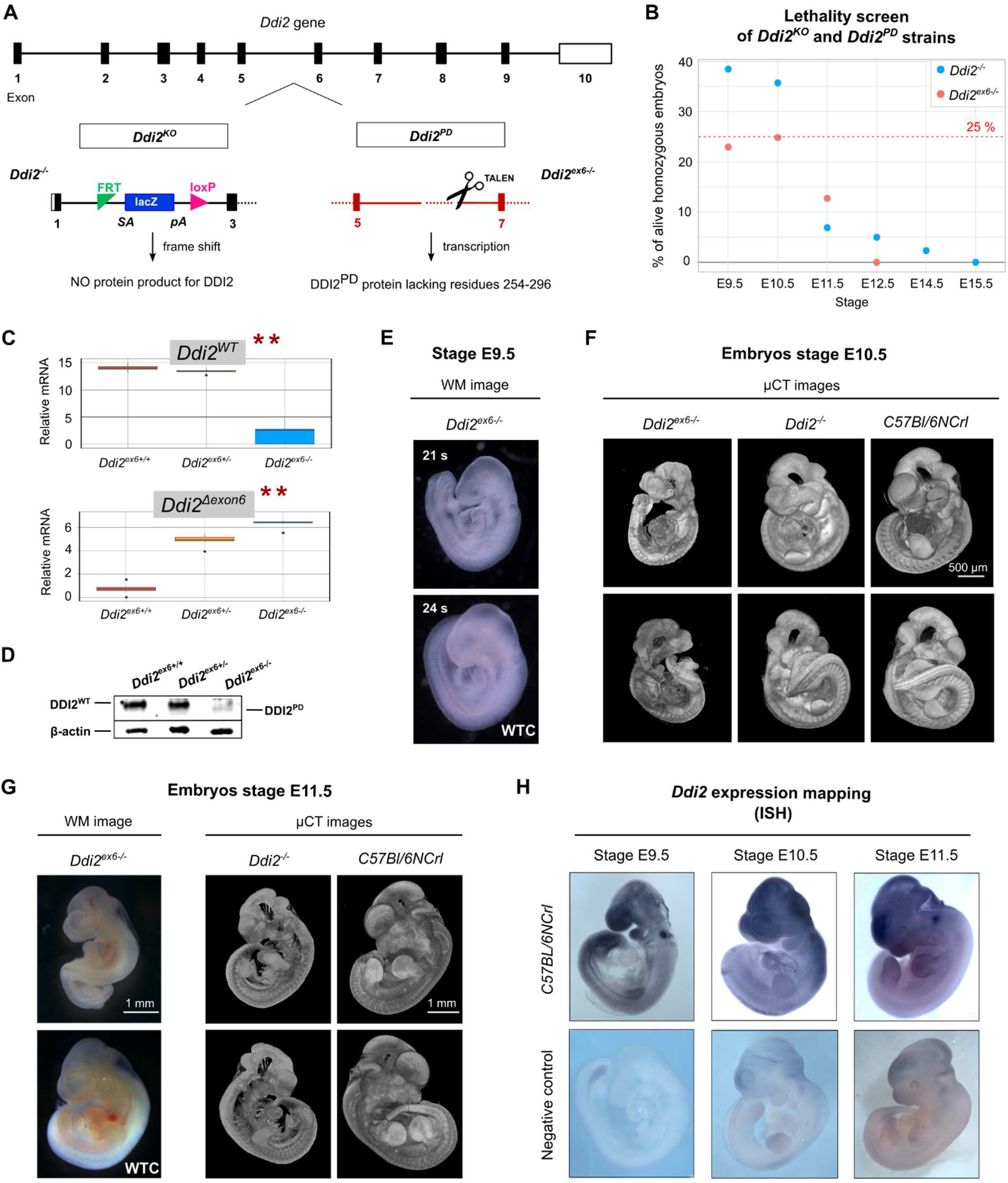
DDI2 deficiency in mice results in mid-gestation embryonic lethality. **(A)** Scheme of *Ddi2* gene alternations used in this study. Two mouse model strains were generated: left – *Ddi2^KO^* carries a *lacZ* reporter gene-tagged allele lacking the critical exon 2 and represents the full knock-out model, right – *Ddi2^PD^* carries a deletion of exon 6 generated by TALEN-mediated excision that results in inactivation of the protease domain. **(B)** Lethality screening of *Ddi2^KO^* and *Ddi2^PD^* mouse strains. As can be observed in the plot showing the percentage of living homozygous embryos of both strains, *Ddi2^ex6-/-^* embryos exhibit narrower window of lethality when compared to *Ddi2^-/-^* embryos. Data from lethality screening was analyzed using the program R, version 3.6.2 (2019-12-12). Detailed distributions of genotypes per stage and per group of living and dead embryos is shown in both barplots and tables in Supplemental Figure S1A and B. **(C)** Boxplots displaying relative expression of *Ddi2^WT^* (top) and truncated *Ddi2^Δexon6^* (bottom) forms of mRNA expressed in *Ddi2^PD^* E10.5 stage embryos (*Ddi2^ex6+/+^* - red, *Ddi2^ex6+/-^* - yellow, *Ddi2^ex6-/-^* - blue). Relative expression was normalized to *Tbp* and *H2afz* housekeeping genes with applied ANOVA statistical analysis (**). Outliers were omitted based on Grubbs’ test. Error bars denote SD. Data were analyzed using the program R, version 3.6.2 (2019-12-12). **(D)** Western blot analysis of E10.5 embryos detected only low amounts of the *DDI2^PD^* (Δ254-296) protein in *Ddi2^ex6-/-^* embryo lysates compared to the expression of DDI2^WT^ in *Ddi2^ex6+/-^* and *Ddi2^ex6-/-^* embryos. β-actin was used as a loading control. **(E)** No difference is yet observed between *Ddi2^ex6-/-^* (top) and *Ddi2^ex6+/+^* (bottom) littermate embryos at developmental stage E9.5. Number of somites is stated in each whole mount image. **(F)** Phenotype analysis of stage E10.5 embryos. Comparison of μCT scans of *Ddi2^ex6-/-^* (right), *Ddi2^-/-^* (middle) and control *C57BL/6NCrl* (left) embryos using surface rendering representation. The 2D images of 3D μCT scans are shown from both sides. The scale bar shows 500 μm. **(G)** Phenotype analysis of both *Ddi2^-/-^* and *Ddi2^ex6-/-^* embryos at developmental stage E11.5. Whole mount image of *Ddi2^ex6-/-^* embryo (top left) compared to its wild-type littermate (bottom left). Surface rendering of *Ddi2^-/-^* and *C57BL/6NCrl* control embryo μCT scans (middle). The 2D images of 3D μCT scans are shown from both sides. WTC - wild-type control. **(H)** *Ddi2* expression analysis using RNA in-situ hybridization of *C57BL/6NCrl* embryos. Stages E9.5, E10.5 and E11.5 are shown. These are the three developmental stages at the beginning of developmental failure onset in the *Ddi2* knock-out model strains.

*Ddi2^-/-^* embryos died between stages E11.5 and E14.5, with only one highly retarded survivor detected at stage E14.5 out of 48 total embryos harvested at this stage (Figure 1B, Figure S1A, B). A significant alteration of the expected Mendelian ratio appeared at stage E11.5, in which half of the embryos were dead and the rest exhibited obvious growth retardation (Figure 1G). Intriguingly, the *Ddi2^ex6-/-^* embryos phenomimic the *Ddi2^KO^* embryos in the onset of the lethal period at E11.5 and with a similar distribution of deceased and growth retarded embryos. However, in contrast to the *Ddi2^KO^* embryos, we observed the first signs of severe growth retardation in the *Ddi2^ex6-/-^* embryos at E10.5 (Figure 1F) and were not able detect living embryos at E12.5, suggesting an earlier appearance of developmental defects and a narrower window of lethality in the *Ddi2^ex6-/-^* embryos (Figure 1B). We next analyzed the expression pattern of both the *Ddi2^WT^* and *Ddi2^Δexon6^* versions of *Ddi2* mRNA in *Ddi2^PD^* at E10.5. While qRT-PCR experiments confirmed expression of *Ddi2^Δexon6^* mRNA in embryonic samples of both the *Ddi2^ex6+/-^* and the *Ddi2^ex6-/-^* genotypes (Figure 1B), the DDI2^PD^ protein was scarcely detectable in embryonic tissue lysates in western blot experiments using an anti-DDI2 antibody (Figure 1C). We then analyzed folding and stability of the recombinant *DDI2^PD^* protein using nuclear magnetic resonance (NMR) and differential scanning fluorimetry (DSF). The 1D NMR spectrum of the *DDI2^PD^* protein showed protein signals of acquired secondary structures, probably resulting from the remaining N-terminal UBL and HDD domains of the crippled *DDI2^PD^* protein. The melting temperature of *DDI2^PD^* could not be measured by DSF, although that of DDI2^WT^ could (Supplemental information, Figure S1C and S1D, respectively). Moreover, levels of *DDI2^PD^* protein expression in a model HEK293-TetOff-A cell line were lower at identical time points after transfection when compared to the DDI2^WT^ protein (Supplemental information, Figure S1E). Taken together these results indicate that *in vivo*, the *DDI2^PD^* protein likely undergoes rapid degradation.

The general nature of the severe growth retardation of both *Ddi2^KO^* and *Ddi2^PD^* strains suggests a systemic effect of *Ddi2* ablation on critical developmental components. In order to map the expression of *Ddi2* during embryonic development we utilized *LacZ* reporter knock-in in *Ddi2^+/-^* mice combined with *Ddi2* hybridization *in situ*. Expression of *Ddi2* at E9.5 was detected in the most dynamically developing regions of the embryo at this stage such as the orofacial processes (maxilla, mandible), the otic placode, the dorsal aorta and the tail bud (Figure 1H). Levels of *Ddi2* mRNA were 2-fold higher at this stage compared to E10.5 and E11.5 (Figure S1F). Expression of *Ddi2* later expanded into most other tissues and was also visible in the developing heart (Figure S1H) and the placenta (Figure S1G). Sagittal sections of the head at stage E14.5 revealed localization in a specific layer of the cortex (forebrain and trigeminal ganglion) in the ectoderm and in mesodermal tissue, such as cranium (Figure S1H). The broad expression of *Ddi2* suggests that *Ddi2* is an essential gene for systemic physiological functions during embryonic development.

We also analyzed the pattern of *LacZ* expression in adult mice. This showed that *Ddi2* is expressed in organs developed from all three germ layers (Figure S2 and S3). The pattern of expression of *Ddi2* in ectodermal and mesodermal tissue was also mapped in the reproductive system of both sexes, the lungs, and the kidneys (Figure S2). *Ddi2* was also detected in the grey matter of the brain, in peripheral neurons, in the epidermis and in a number of glands of the main ectodermal tissue (Figure S3). Note that *Ddi2* is also expressed in the bone marrow and the endothelial layer of vessels, which was a key criterion for selection of an endothelial cell line as a proper human cell line model for further experiments *in vitro*.

### Impairment of DDI2 function in mice and endothelial cells alters expression of UPS genes

To investigate on the molecular level the effect of impairing DDI2 function, we decided to further analyze the *Ddi2^PD^* strain embryos (which exhibited a more extreme phenotype). DDI2 has been identified as the major protease responsible for the cleavage reaction which activates the transcription factor TCF11/NRF1 (Dirac-Svejstrup et al., 2020; Koizumi et al., 2016; Northrop et al., 2020) thus modulating expression of a subset of UPS factors and almost all proteasome subunits (Radhakrishnan et al., 2010; Steffen et al., 2010). We therefore performed qRT-PCR analysis of *Nfe2l1* (NRF1) and its downstream target genes in *Ddi2^PD^* embryos harvested at stages prior to the onset of lethality. *Nfe2l1* and *Ngly1* each showed a two-fold increase in expression at stage E10.5 compared to *Ddi2^ex6+/+^*, and this increase in expression was statistically significant throughout all three embryonic stages. The proteasomal subunit genes transactivated by NRF1 (*Psma4, Psma6, Psmb6*) showed no striking differences in expression between the three embryonic stages. On the other hand, expression of the shuttling proteins *Rad23a* and *Rad23b* increased upon depletion of DDI2 function in embryos at stage E10.5 compared to *Ddi2^ex6+/+^*, and the increase in expression was also statistically significant throughout all three embryonic stages (Figure 2A).

**Figure 2:**
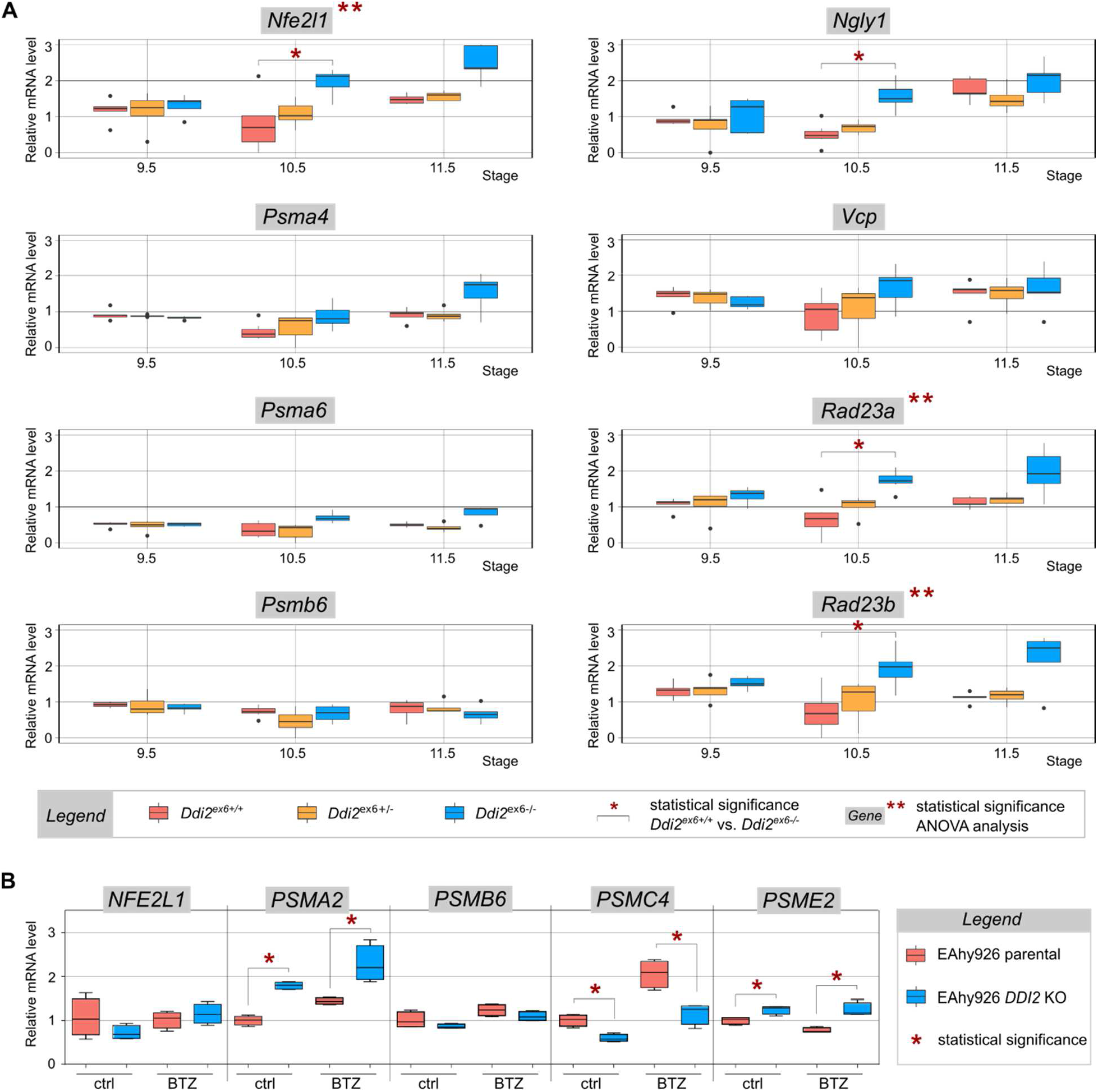
Impairment of DDI2 function in mice and endothelial cells alters expression of the UPS genes. **(A)** qRT-PCR analysis of NRF1-regulated genes in the UPS pathway in *Ddi2^em1^/Ph* embryos in stages prior to the onset of lethality. Legend: *Ddi2^ex6+/+^* - red, *Ddi2^ex6+/-^* - yellow, *Ddi2^ex6-/-^* - blue; E9.5 (n=5), E10.5 (n=7), E11.5 (n=5). Relative expression of genes was normalized to the *Tbp* and *H2afz* housekeeping genes; outliers were omitted based on Grubbs’ test. Statistical significance was calculated either for each gene throughout all three developmental stages with application of ANOVA statistical analysis (**) or using a linear mixed-effects model (LMM) for comparison of gene expression between wild-type and homozygous embryos at each developmental stage (*). Both analyses were subjected to Bonferroni correction. Data and boxplots were processed using the program R, version 3.6.2 (2019-12-12). **(B)** qRT-PCR analysis of *NFE2L1* (TCF11/NRF1) and proteasomal subunits *PSMA2* (α2), *PSMB6* (β1), *PSMC4* (RPT3) and *PSME2* (PA28β) mRNA levels in EAhy926 parental (red) and *DDI2* KO (blue) cells. Cells treated with 50 nM BTZ for 8 hours to inhibit proteasomal activity are compared to non-treated controls. Messenger RNA levels were normalized to *RPLP0*. No outliers were found using Grubbs’ test. Statistical significance was calculated individually within the BTZ treatment experiment and the non-treatment control for differences in gene expression between the parental and *DDI2* KO cell line. Unpaired two-sided t-test (n=4) was calculated and significance was determined after application of the Holm-Šidák correction (*α = 5%) using GraphPad Prism 6 software.

Given the fact that *Ddi2* is expressed mainly in highly dynamic tissues during embryonic development in mice we next aimed to explore how human *DDI2* knock-out (*DDI2* KO) endothelial cells cope with disrupted proteostasis. We treated previously described *DDI2* KO EAhy926 cells (Nowak et al., 2018) and EAhy926 parental cells with bortezomib (BTZ), an inhibitor of the chymotrypsin-like activity of the proteasome (Cavo et al., 2015). To address the importance of DDI2 in the regulation of proteasome gene expression (Figure 2B) we performed qRT-PCR analysis of messenger RNA (mRNA) levels of representative subunits *PSMA2*/α2, *PSMB6/β1, PSMC4*/RPT3, *PSME2*/PA28β of the core and regulatory proteasome particle. Relative mRNA levels of *NFE2L1* in *DDI2* KO cells were modestly increased upon BTZ treatment compared to BTZ treated parental cells. As expected and consistent with previous data, expression of proteasomal subunits (except PSME2/PA28β) increased in EAhy926 parental cells upon proteasome inhibition, although to different extents (Radhakrishnan et al., 2010; Steffen et al., 2010). Expression of *PSMB6* and *PSMC4* was impaired in *DDI2* KO cells compared to parental cells upon proteasome inhibition (Koizumi et al., 2016; Northrop et al., 2020). Upregulation of *PSMA2* was not affected in the *DDI2* KO background, suggesting that transcription factors other than TCF11/NRF1 can transactivate the *PSMA2* gene in this context. Unexpectedly, expression of *PSME2* was significantly upregulated in *DDI2* KO cells compared to BTZ treated parental cells, indicating that *DDI2* KO cells upregulate alternative proteasome regulators (see also Figure 3).

**Figure 3:**
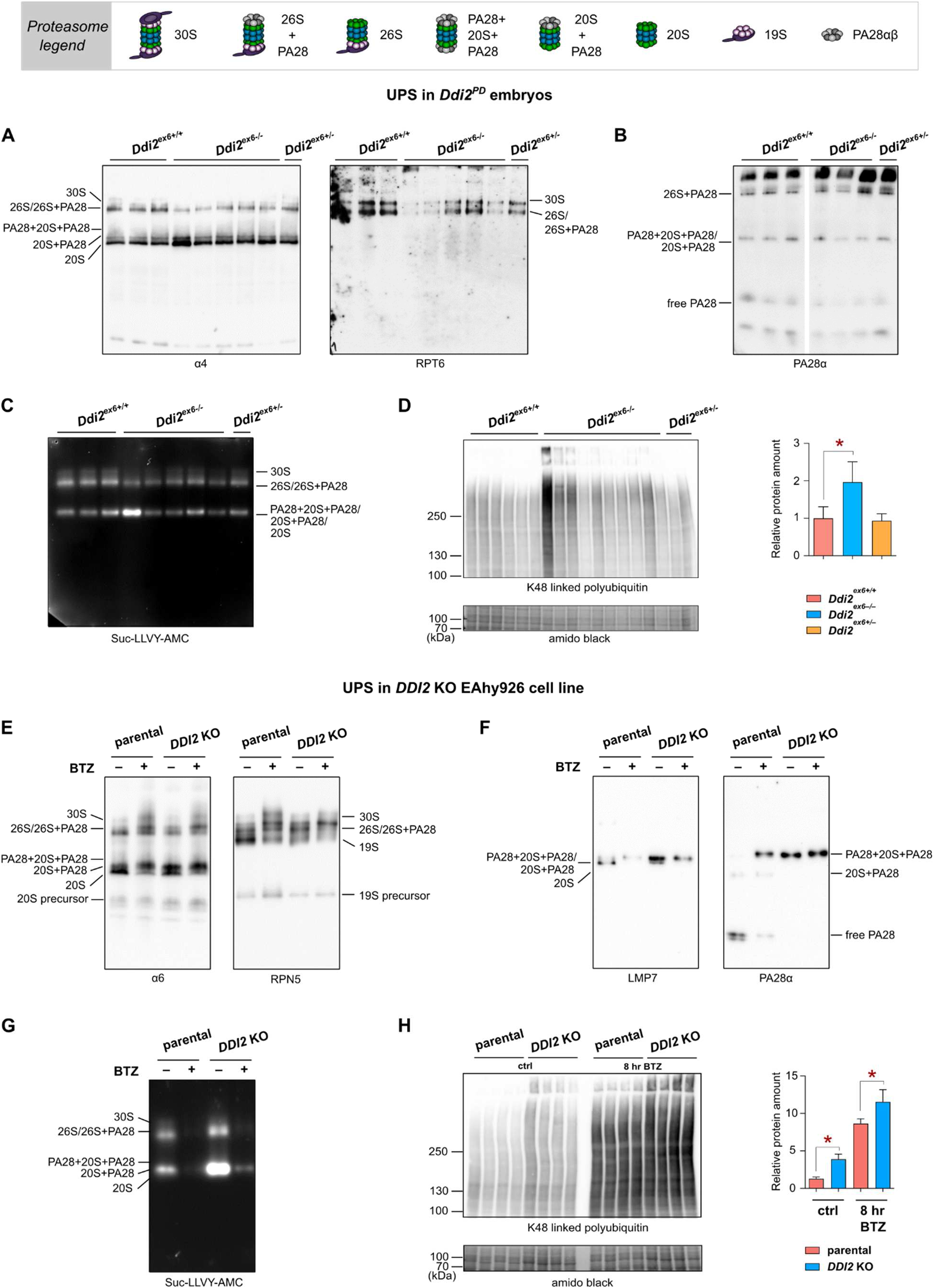
Deficiency of DDI2 function leads to altered proteasome composition and activity accompanied by accumulation of polyubiquitinated proteins of high molecular weight. Proteasomal complexes analyzed in the immunoblots are schematically illustrated in the legend. **(A), (B)** Immunoblots of native tissue lysates of *Ddi2^ex6+/+^* (n=8), *Ddi2^ex6-/-^* (n=15, resp. n=12 for PA28α) and *Ddi2^ex6+/-^* (n=6) embryos were probed for proteasomal subunits α4, RPT6 and PA28α. Proteins (15 μg of total protein/lane) were separated on non-denaturing gels, and amido black staining of the blotted membrane was used as loading control (see Figure S5 in Supplemental Information). **(C)** Chymotrypsin-like activity of native tissue lysates of *Ddi2^ex6+/+^* (n=8), *Ddi2^ex6-/-^* (n=15) and *Ddi2^ex6+/-^* (n=6) embryos was measured in gels based on the hydrolysis of the fluorogenic substrate Suc-LLVY-AMC. **(D)** Western blot analysis of the insoluble fraction of *Ddi2^ex6+/+^* (n=6, red), *Ddi2^ex6-/-^* (n=10, blue) and *Ddi2^ex6+/-^* (n=2, yellow) embryo tissue lysates show accumulation of polyubiquitinated proteins of higher molecular weight upon impairment of *Ddi2* function. The membrane was probed with an anti-K48 linked polyubiquitin antibody, and the expression of ubiquitin-conjugates with a molecular weight above 250 kDa was densitometrically quantified using ImageJ software and normalized to the amido black loading control. The outlier in lane 7 was excluded from the calculation. Statistical significance was determined between *Ddi2^ex6+/+^* and *Ddi2^ex6-/-^* and between *Ddi2^ex6+/+^* and *Ddi2^ex6+/-^* using GraphPad Prism 6 software (mean ± SD, *p value < 0.05, unpaired two-sided t-test). **(E), (F)** Immunoblots of native cell lysates of EAhy926 parental and *DDI2* KO cells treated with 50 nM BTZ for 8 hours compared to non-treatment controls. The immunoblots were probed for the proteasomal subunits α6, RPN5, LMP7 and PA28α (n=4). Proteins (20 μg of total protein/lane) were separated on nondenaturing Bis-Tris gels, and amido black staining was used as loading control (see Figure S5 in Supplemental Information). **(G)** Chymotrypsin-like activity of cell lysates of EAhy926 parental and *DDI2* KO cells treated with 50 nM BTZ for 8 hours compared to non-treatment controls (n=4) was measured in a gel-based assay with the fluorogenic substrate Suc-LLVY-AMC. **(H)** Western blot analysis of whole cell extracts of EAhy926 parental (red) and *DDI2* KO (blue) cells treated with 50 nM BTZ for 8 hours compared to non-treated controls. Staining and quantification was performed as described in (D). Statistical significance of the difference in the relative amount of protein between the parental and DDI2 KO cells was calculated separately for the BTZ treatment experiment and control samples using GraphPad Prism 6 software (*p value < 0.05, n=4, unpaired two-sided t-test).

### Deficiency of DDI2 function leads to altered proteasome composition and activity accompanied by accumulation of polyubiquitinated proteins of high molecular weight

To verify our conclusions from the mRNA expression experiments described above with respect to proteasome expression, composition, and activity, we next studied the effect of DDI2 deficiency in *Ddi2^em1^/Ph* embryos and EAhy926 *DDI2* KO cells by native PAGE. Decreased 26S and 30S proteasome levels were detected in *Ddi2^em1^/Ph* embryos by staining cellular proteasome complexes for the alpha subunit α4 and the 19S regulator subunit RPT6 (Figure 3A). PA28/11S can act as an alternative activator for proteasomes. In this context, however, its association with proteasomes was almost unchanged (Figure 3B). Concomitant with decreased levels of 26S/30S proteasomes, we observed a reduction in chymotrypsin-like (CT-L) activity of 26S and 30S proteasomes in homozygous *Ddi2^em1^/Ph* embryos (Figure 3C). This was accompanied by significant accumulation of K48-linked ubiquitin conjugates in homozygous *Ddi2^em1^/Ph* embryos compared to wild-type littermates (Figure 3D).

The abundance of different proteasome complexes and ubiquitin conjugates in EAhy926 parental and *DDI2* KO cells was also analyzed in BTZ treated cells compared to untreated controls. Immunoblots of native PAGE gels stained for the α6 or RPN5 subunits revealed elevated levels of 26S/30S proteasomes as well their precursor complexes in parental cells after BTZ treatment due to upregulation of proteasomes by TCF11/NRF1 (Figure 3E lanes 1 and 2). *DDI2* KO cells failed to upregulate 26S/30S proteasomes in response to proteasome inhibition and displayed slightly decreased levels of proteasomes under standard growth conditions (Figure 3E, RPN5, lanes 3 and 4). The decreased proportion of free 19S particles (see RPN5 staining) suggested that 19S regulatory particles associate with 20S core proteasomes to form single or double capped proteasome complexes and enhance proteolytic capacity under these conditions (Finley, 2009).

Of note, both BTZ treatment and DDI2 deficiency led to an altered proteasome composition with the incorporation of β5i/LMP7 into proteasomes in KO cells and preferential association of the PA28/11S activator to proteasome complexes in response to BTZ in parental cells and under all conditions tested in DDI2-deficient cells (Figure 3F). Thus, *DDI2* KO cells compensate for impaired proteasome formation by induction of the cytokine inducible immuno- and hybrid proteasomes (Kruger and Kloetzel, 2012). Both parental and *DDI2* KO cells express β5i/LMP7 under standard conditions, but this is expressed at a notably higher level in the KO. β5i/LMP7 was not expressed in *Ddi2^em1^/Ph* embryos as confirmed by SDS-PAGE (data not shown). Levels of *PSME2* mRNA were not significantly elevated upon BTZ treatment in parental cells, so that the increase of 20S-PA28 complexes can be explained by augmented assembly (Welk et al., 2016).

As expected-CT-L activity was almost completely eliminated in BTZ treated parental cells. In contrast-the *DDI2* KO cells exhibited higher CT-like activity, especially of the 20S proteasome, under untreated conditions compared to parental cells, and maintained a modest activity after BTZ treatment (Figure 3G). This is due to increased recruitment of PA28/11S- which is known to activate the peptide hydrolysis activity of 20S proteasome complexes (Figure 3F) (Whitby et al., 2000). In conclusion, deficiency of DDI2 *in vivo* as well as *in cellulo* leads to altered proteasome complexes with enhanced incorporation of the PA28/11S regulator instead of 19S, which results in higher peptide hydrolysis activity.

Nevertheless-*DDI2* KO cells and embryos are unable to compensate for impaired proteasome activity. Analysis of ubiquitin conjugates by western blot (Figure 3D, H) showed a significant enrichment of K48 linked polyubiquitylated proteins in homozygous *Ddi2^em1^/Ph* embryos as well as EAhy926 *DDI2* KO cells in all experimental setups. Although the EAhy926 parental cells also accumulate polyubiquitylated proteins of high molecular weight upon BTZ treatment-this is even more pronounced in the *DDI2* KO (Figure 3H).

Notably, and consistent with recent studies of the effects of DDI2 deficiency in cells (Dirac-Svejstrup et al., 2020), ubiquitin conjugates in both Ddi2^ex6-/-^ embryos and *DDI2* KO cells were of considerably higher molecular weight and could be detected especially in stacking gels (Figure 3D, H), whereas such conjugates were less abundant in the wild-type embryos and untreated parental cells.

### The NRF2 pathway is activated upon loss of DDI2

It was previously shown that the functions of *Nfe2l1* and *Nfe2l2* in the oxidative stress response overlap during mouse early embryonic development (Leung et al., 2003). Because *Nfe2l1* function is impaired in our model *Ddi2^em1^/Ph* embryos due to *Ddi2* functional deficiency, we next investigated whether the NRF2 pathway is activated in developmental stages prior to the onset of lethality. qRT-PCR analysis revealed a statistically significant increase in expression of *Nfe2l2* mRNA in *Ddi2^ex6-/-^* embryos compared to *Ddi2^ex6+/+^* at E10.5 and E11.5, and also of several genes (*Gclm, Gss, Hmox1*, and *iNos2*) involved in oxidative stress response (Figure 4A) (Vomund et al., 2017). Of these, *Hmox1* exhibits the largest increase in expression upon loss of *Ddi2* function at stages E10.5 and E11.5. Interestingly, the expression level of the second DDI2 substrate *Nfe2l3* mRNA did not change in embryonic stages prior to the onset of lethality in *Ddi2^em1^/Ph* embryos (Chevillard and Blank, 2011; Chowdhury et al., 2017). NRF2 and heme oxygenase (HO-1) mRNA expression correlated with moderate increase of both proteins (Figure S7 in Supplemental Information).

**Figure 4:**
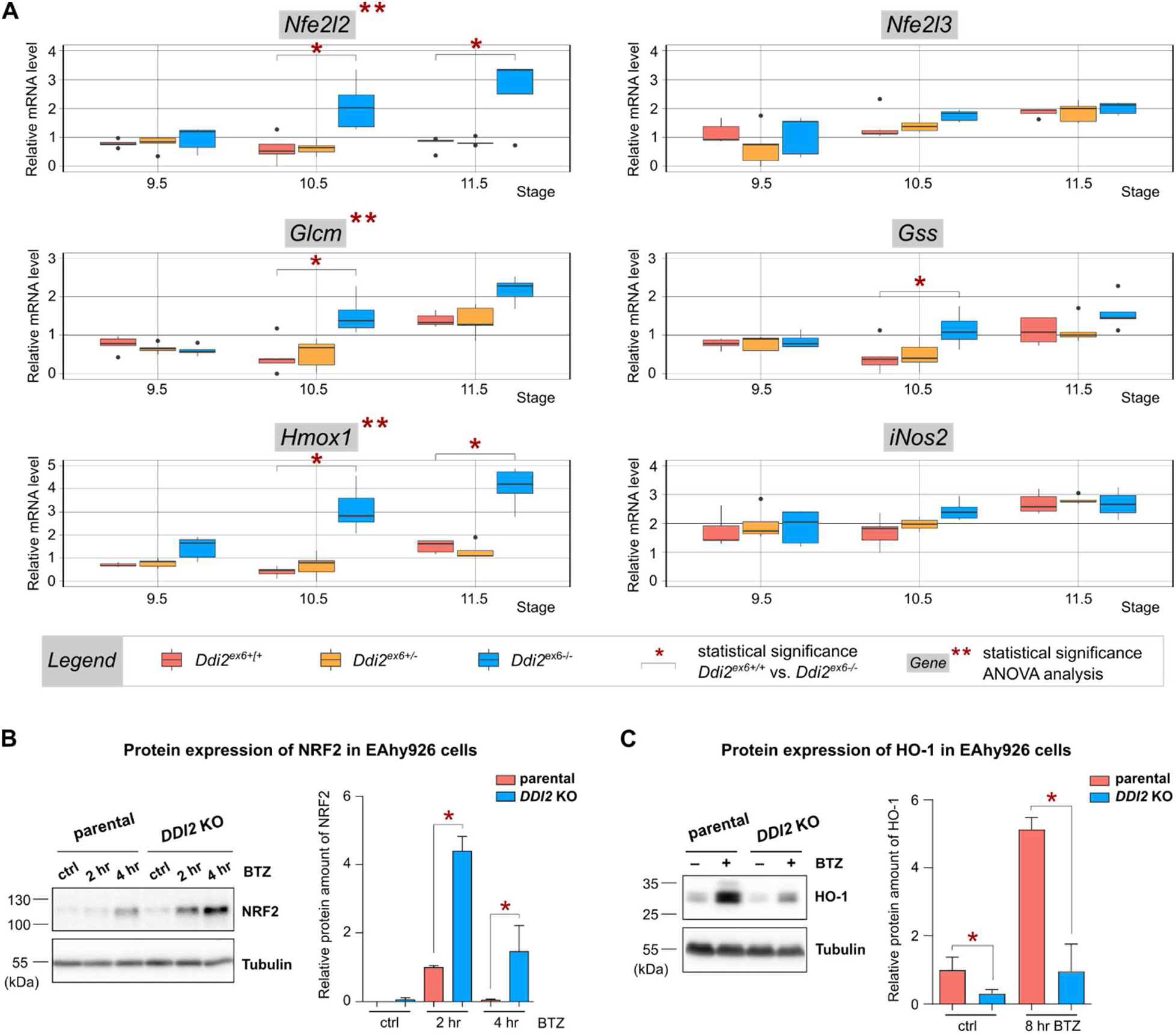
The NRF2 pathway is activated upon loss of DDI2. **(A)** qRT-PCR analysis of NRF1 and NRF2-regulated genes involved in the oxidative stress response in *Ddi2^em1^/Ph* embryos in stages prior to the onset of lethality. Legend: *Ddi2^ex6+/+^* - red, *Ddi2^ex6+/-^* - yellow, *Ddi2^ex6-/-^* - blue; E9.5 (n=5), E10.5 (n=7), E11.5 (n=5). Relative expression of genes was normalized to *Tbp* and *H2afz* housekeeping genes; outliers were omitted based on Grubbs’ test. Statistical significance was calculated either for each gene throughout all three developmental stages with application of ANOVA statistical analysis (**) or using a linear mixed-effects model (LMM) for comparison of gene expression between wild-type and homozygous embryos at each developmental stage (*). Both analyses were subjected to Bonferroni correction. Data and boxplots were processed using the program R, version 3.6.2 (2019-12-12). **(B)** Western blot analysis of NRF2 and HO-1 expression in whole cell extracts of EAhy926 parental (red) and *DDI2* KO (blue) treated with 50 nM BTZ for 2 and 4 hours (NRF2 detection) or 8 hours (HO-1 detection) with non-treatment controls. Protein expression was quantified using ImageJ software and normalized to a tubulin control signal. Statistical significance was determined using GraphPad Prism 6 software (n=4, mean ± SD, *p value < 0.05, unpaired two-sided t-test).

Expression of NRF2 upon BTZ treatment was strongly induced in the DDI2-deficient cell model. Expression of NRF2 upon BTZ treatment was also more pronounced and occurred at an earlier time point in the KO compared to parental cells. HO-1 expression in turn was more strongly induced in parental cells compared to DDI2-deficient cells indicating that HO-1 expression in endothelial cells is strongly dependent on TCF11/NRF1 and less on NRF2.

### Functional loss of DDI2 leads to induction of downstream UPR markers

It was previously shown that activation of TCF11/NRF1 transcriptional pathways by proteotoxic stress during proteasome impairment is accompanied by induction of UPR and ISR (Poli et al., 2018; Sotzny et al., 2016; Studencka-Turski et al., 2019). Therefore, we examined whether the functional loss of DDI2 in mouse alters downstream UPR signaling. We performed qRT-PCR analysis of *Ddi2^em1^/Ph* embryos harvested at stages prior to the onset of lethality and analyzed the mRNA levels of characteristic UPR genes (Figure 5A). Expression of *Atf4* and *Chop* mRNA was significantly increased in *Ddi2^ex6-/-^* embryos in stages E10.5 and E11.5 compared to *Ddi2^ex6+/+^*, and the increase was statistically significant throughout the three embryonic stages. *Herpud1*, a gene upregulated upon ER stress, exhibits increased expression in DDI2-deficient embryos compared to *Ddi2^ex6+/+^* (and also throughout the three embryonic stages), while ubiquillins which are known to attenuate the induction of UPR genes during ER stress (such as CHOP, HSPA5 and PDIA2) showed increased expression in *Ddi2^ex6-/-^* embryos compared to *Ddi2^ex6+/+^* at the edge of statistical significance.

**Figure 5:**
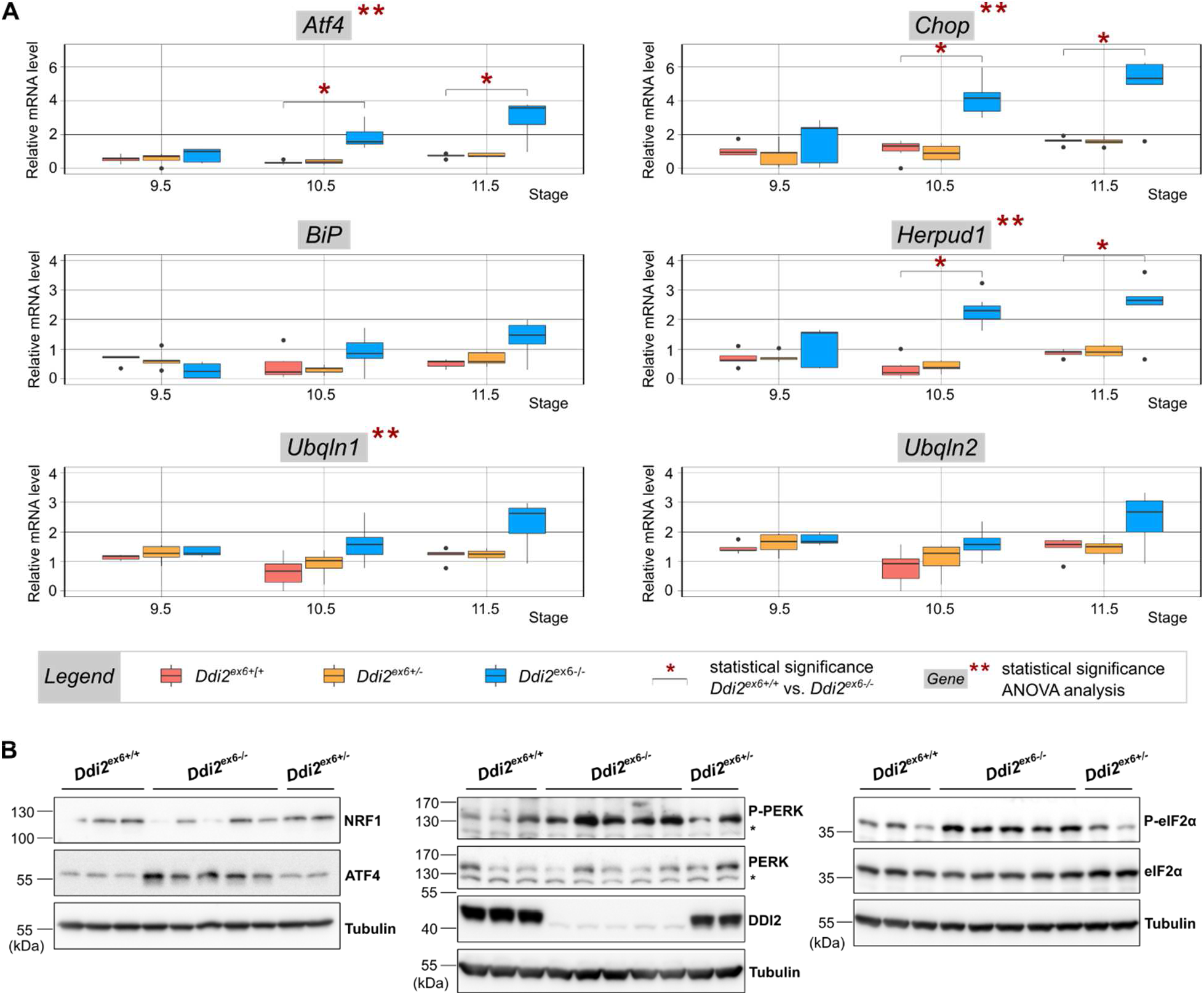
Functional loss of DDI2 in mouse embryos leads to increased phosphorylation of PERK and induction of downstream UPR markers. **(A)** qRT-PCR analysis of genes involved in UPR in *Ddi2^em1^/Ph* embryos among stages prior to the onset of lethality. Legend: *Ddi2^ex6+/+^* - red, *Ddi2^ex6+/-^* - yellow, *Ddi2^ex6-/-^* - blue; E9.5 (n=5), E10.5 (n=7), E11.5 (n=5). Relative expression of genes was normalized to *Tbp* and *H2afz* housekeeping genes; outliers were omitted based on Grubbs’ test. Statistical significance was calculated either for each gene throughout all three developmental stages with application of ANOVA statistical analysis (**) or using a linear mixed-effects model (LMM) for comparison gene expression between wild-type and homozygous embryos at each developmental stage (*). Both analyses were subjected to Bonferroni correction. Data and boxplots were processed using the program R, version 3.6.2 (2019-12-12). **(B)** Immunoblots of tissue lysates of *Ddi2^ex6+/+^* (n=6), *Ddi2^ex6-/-^* (n=10) and *Ddi2^ex6+/-^* (n=3) embryos were stained for NRF1, ATF4, P-PERK, PERK, DDI2, P-eIF2α and eIF2α. Tubulin was used as a loading control. Asterisks denote unspecific bands.

Western blot analysis of embryo lysates (harvested at E10.5) also revealed activation of the PERK pathway of the UPR. We detected increased auto-phosphorylation of PERK (P-PERK) in *Ddi2^ex6-/-^* embryos compared to *Ddi2^ex6+/+^* and subsequent increased phosphorylation of eIF2α (P-eIF2α). This resulted in increased expression of ATF4, a gene which governs multiple signaling pathways including autophagy, oxidative stress, inflammation, and translation and is upregulated in many pathological conditions (Figure 5B). ATF4 in turn transactivates CHOP (Figure 5A), indicating that cell death is induced in these stages of embryonic development (Hetz et al., 2020).

The induction of the UPR we observed in DDI2 deficient embryos was next investigated in *DDI2* KO cells compared to parental cells as well as in response to BTZ in time course experiments. We observed induction of *ATF4* mRNA in DDI2-deficient cells even under control conditions, which was more pronounced after treating for 8 hours with BTZ and was less pronounced than that observed after treating parental cells for 16 hours with BTZ (Figure 6A). This suggests that ATF4 could be aberrantly regulated. Consistent with this stronger induction of *ATF4* mRNA, ATF4 protein levels were increased in the nucleus (Figure 6B) and in total lysates of DDI2 deficient cells under all conditions (Figure 6C). In addition, we detected induced protein expression of CHOP as a downstream target of ATF4 after a 16 hour BTZ treatment in DDI2-deficient cells (Figure 6C). As expected and consistent with previously published work, TCF11/NRF1 processing and nuclear translocation was almost completely impaired in *DDI2* KO cells. However, a small amount of TCF11/NRF1 could still translocate into the nucleus upon BTZ treatment, indicating an additional mechanism for TCF11/NRF1 activation (Northrop et al., 2020).

**Figure 6:**
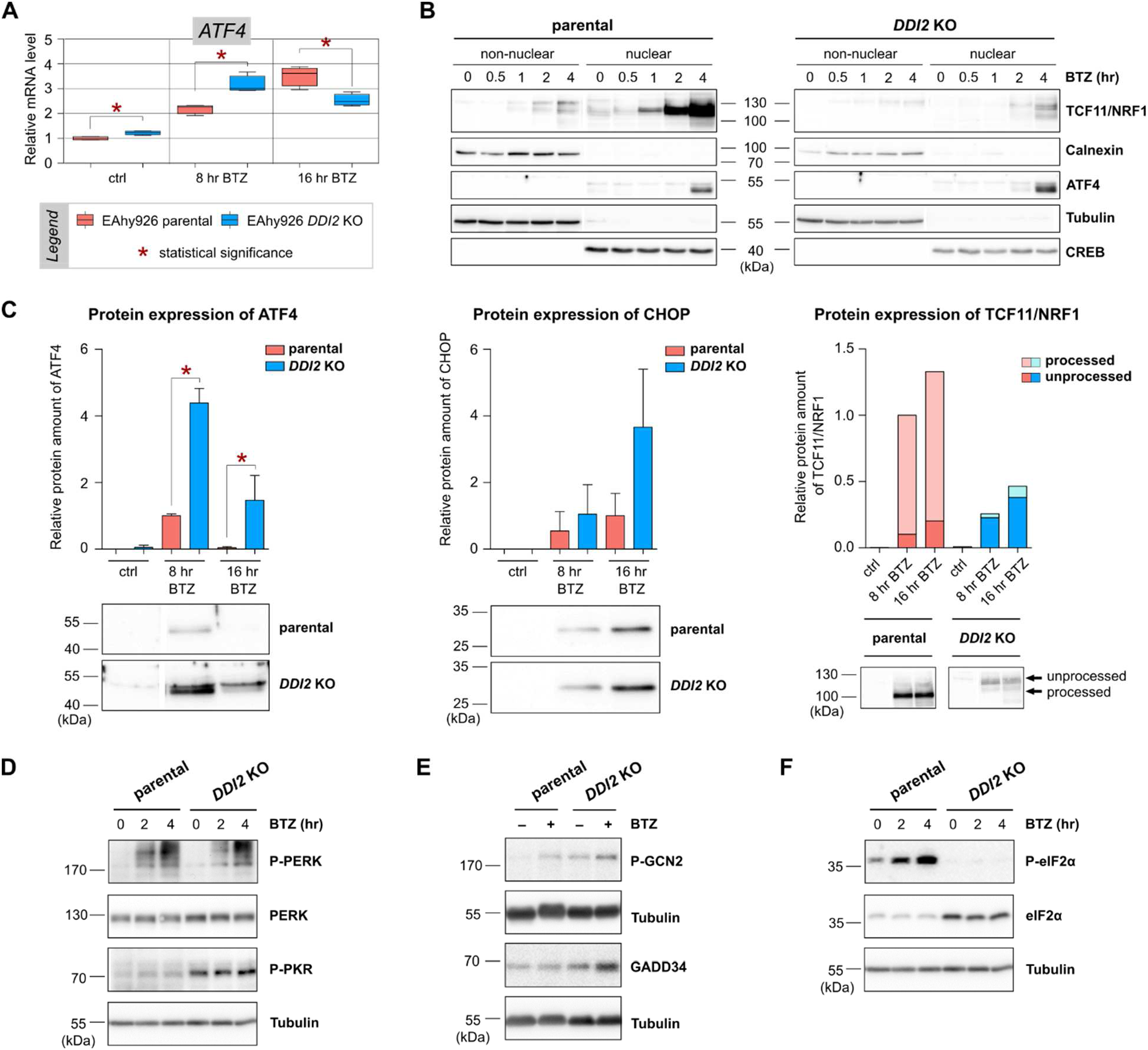
*DDI2* KO cells exhibit strongly diminished activation of TCF11/NRF1 accompanied by induction of downstream UPR markers. **(A)** qRT-PCR analysis of *ATF4* mRNA level in EAhy926 parental (red) and *DDI2* KO (blue) cells. Cells were treated with 50 nM BTZ for 8 and 16 hours. Non-treated cells were used as a negative control; mRNA amounts were normalized to *RPLP0* and Grubbs’ outlier test was applied. Statistical significance of the difference between parental and *DDI2* KO cells was calculated individually for each BTZ treatment time point and non-treatment control sample using the unpaired two-sided t-test (n=4) and significance was determined after application of the Holm-Šidák correction (*α = 5%) using GraphPad Prism 6 software. **(B)** Cellular fractionation of EAhy926 parental and *DDI2* KO cells. The non-nuclear and nuclear fractions are shown. Cells were treated with 50 nM BTZ in a 4 hour time course (n=5). Immunoblots were stained for TCF11/NRF1 and ATF4. Calnexin, tubulin and CREB served as marker proteins for the individual fractions. For separate fractionation of chromatin-associated proteins see Figure S6B in the Supplemental Information. **(C)** Western blot analysis of whole cell extracts of EAhy926 parental (red) and *DDI2* KO (blue) treated with 50 nM BTZ for 8 and 16 hours and non-treatment controls. The membranes were stained for ATF4, CHOP and TCF11/NRF1. For the complete time course experiment see Figure S6C in the Supplemental Information. Expression of the analyzed proteins was normalized to an amido black loading control and densitometrically quantified using ImageJ software. As the EAhy926 parental and *DDI2* KO samples of each time course experiment were analyzed on separate immunoblots, another western blot was run for normalization of protein expression levels (see Figure S6D in the Supplemental Information). Amounts of the ATF4 and TCF11/NRF1 proteins were normalized to 8 hour BTZ treated parental cells, whereas CHOP was normalized to a 16 hour BTZ treatment. Asterisks denote statistical significance, which was determined using GraphPad Prism 6 software (n=3, mean ± SD, *p value < 0.05, unpaired two-sided t-test). For complete time course experimental results and the comparative normalization immunoblot see Figure S6D in the Supplemental Information. **(D)** Western blot analysis of protein expression in whole cell extracts of EAhy926 parental and *DDI2* KO cells treated with 50 nM BTZ in a time course of 4 hours (n=3). Membranes were probed for P-PERK, PERK and P-PKR, and tubulin served as a loading control. **(E)** Western blot analysis of protein expression in whole cell extracts of EAhy926 parental and *DDI2* KO cells treated with 50 nM BTZ for 8 hours (n=4). Non-treated cells served as a control. Membranes were probed for P-GCN2 and GADD34. Tubulin served as a loading control. **(F)** Western blot analysis of protein expression in whole cell extracts of EAhy926 parental and *DDI2* KO cells treated with 50 nM BTZ in a time course of 4 hours (n=3). Membranes were probed for P-eIF2α and eIF2α, and tubulin served as a loading control.

One of the major downstream events of the UPR for ER stress and the ISR for cytosolic stress is phosphorylation of eIF2α by the kinases PERK, PKR, and GCN2 (Pakos-Zebrucka et al., 2016). In accordance with the concept of cellular responses to proteotoxic stress and activation of eIF2α kinases, we observed increased levels of the phosphorylated forms of PERK, PKR, and GCN2 in both DDI2-deficient cells and BTZ-treated parental cells due to the activation of autophosphorylation (Figure 6D, 6E). Autophosphorylation of the eIF2α kinases PKR and GCN2 was even more pronounced in DDI2-deficient cells than in parental cells. Consequently, parental cells also exhibited increased levels of phosphorylation of eIF2α itself in response to BTZ treatment. Surprisingly, we could not detect increased levels of phosphorylation of eIF2α in DDI2-deficient cells even though the respective kinases were activated. Levels of the unphosphorylated form of eIF2α, however, were significantly increased (Figure 6F). This might be the result of induced expression of the GADD34 phosphatase subunit in *DDI2* KO cells (Figure 6E), which could dephosphorylate eIF2α. In summary, these data suggest that DDI2-deficiency caused severe proteotoxic stress and led to chronic activation of the UPR and ISR pathways with dysfunctional eIF2α phosphorylation.

### Deletion of DDI2 causes type I interferon signaling

The data presented so far suggest that DDI2 deficiency results in severe proteotoxic stress with embryonic cell death starting at stage E11.5. This is also consistent with our observation of high levels of CHOP induction at stage E10.5 (see Figure 5A). *DDI2* KO cells, however, exhibited only moderate levels of CHOP induction and did not show signs of growth impairment or increased cell death (see Figure S6 in the Supplemental Information). In the case of permanent proteasome impairment by proteasome inhibitors or by loss of function mutations of proteasome subunits, induction of type I IFN was observed ((Ebstein et al., 2019; Studencka-Turski et al., 2019). We therefore investigated activation of IFN by endogenous mechanisms in DDI2-deficient embryos and cells.

Signals such as damage associated molecular patterns (DAMPS) are recognized by pattern recognition receptors (PRRs) of the cell autonomous innate immune response to promote autophosphorylation of TBK1/IKKε/DDX3, subsequent phosphorylation of the transcription factors IRF3 and IRF7, and activation of an IFN signature by their target genes. IFNa and ß generated in this way may in turn stimulate autocrine and paracrine activation of IFN signaling (Figure 7A). We therefore stained lysates of DDI2 deficient embryos for both the unphosphorylated and phosphorylated forms of TBK1 and STAT1 in immunoblots. We observed upregulation of phospho-TBK1, but no phosphorylation of STAT1 (Figure 7B). This indicates that the cell autonomous innate immune receptors detected danger signals and activated TBK1 without downstream induction of IFN production and STAT1 phosphorylation. Consequently, we did not detect significant IFN-stimulated gene (ISG) expression in DDI2-deficient embryos (Figure S7 in the Supplemental Information).

**Figure 7:**
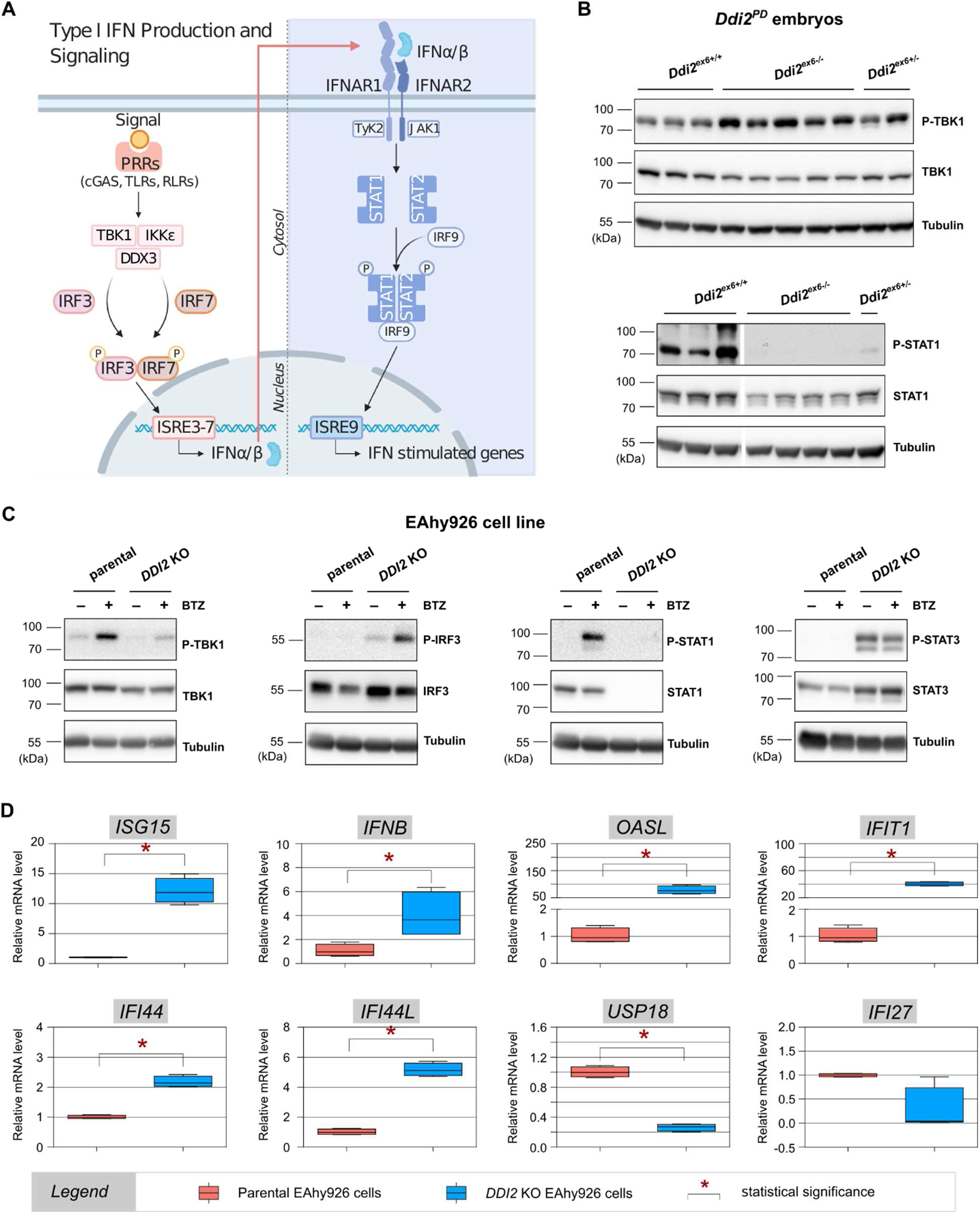
Depletion of DDI2 affects type I interferon signaling. **(A)** Simplified scheme of one of the major type I interferon (IFN) production pathways and downstream signaling. The activation of various pattern recognition receptors (PRRs) initiates activity of IκB kinase-ε (IKKε) and TANK-binding kinase-1 (TBK1). This leads to DEAD box protein 3 (DDX3) mediated phosphorylation of transcription factors IFN regulatory factor 3 (IRF3) and IRF7 which serve as transcriptional activators of type I IFN. Upon binding of IFNα/β to interferon alpha/beta receptor 1 and 2 (IFNAR1/2) the two Janus kinases (JAK) bound to the receptor chains, JAK1 and tyrosine kinase 2 (TYK2), are activated. Signal transducer and activator of transcription 1 and 2 (STAT1/STAT2) are subsequently phosphorylated and released. These transcription factors form heterodimers and interact with IRF9. This complex migrates to the nucleus and induces transcription of IFN stimulated genes (ISGs). The scheme was Created with BioRender.com. **(B)** Differences in protein expression of P-TBK1, TBK1, P-STAT1 and STAT1 were analyzed on immunoblots using tissue lysates of *Ddi2^ex6+/+^* (n=6), *Ddi2^ex6-/-^* (n=10) and *Ddi2^ex6+/-^* (n=3) embryos. Tubulin was used as a loading control. **(C)** Western blot analysis of protein expression in whole cell extracts of EAhy926 parental and *DDI2* KO cells treated with 50 nM BTZ for 8 hours (n=4). Non-treated cells served as a control. Membranes were probed for P-TBK1, TBK1, P-IRF3, IRF3, P-STAT1, STAT1, P-STAT3 and STAT3, and tubulin was used as a loading control. **(D)** qRT-PCR analysis of *ISG15, IFNB, OASL, IFIT1, IFI44, IFI44L, USP18* and *IFI27* mRNA levels in untreated EAhy926 parental (red) and *DDI2* KO (blue) cells. Messenger RNA amounts were normalized to *RPLP0* and Grubbs’ outlier test was applied. Statistical significance was calculated using the unpaired two-sided t-test and significance was determined after application of Holm-Šidák correction (with *α = 5%) using GraphPad Prism 6 software.

In contrast, in DDI2-deficient cells we observed strong phosphorylation of IRF3, but did not detect phosphoTBK1. Interestingly, expression of STAT1 was completely abolished in DDI2 KO cells, and phosphoSTAT1 was therefore not detected (Figure 7C). Increased phosphorylation of IRF3 in DDI2 KO cells was accompanied by chronic induction of specific ISGs along with *IFNB* (encoding IFN-β) and, with the exception of *IFI27* and *USP18* (Figure 7D), a signature typical for interferonopathies (Rice et al., 2017). Moreover, DDI2-deficient cells activated phosphorylation of STAT3 instead of STAT1 (Figure 7C). In contrast, parental cells induced the classical pathway by phosphorylation of TBK1 as well as phosphorylation of STAT1, indicating activation of the cell autonomous innate response and low-grade induction of a type I IFN signature in response to proteasome impairment (Figure 7C and supplementary Figure S7D).

## DISCUSSION

Proteostasis in multicellular organisms is differentially affected during all stages of life by processes such as development and differentiation, metabolic changes, immune responses, and aging (Garcia-Prat et al., 2017; Hipp et al., 2019; Llamas et al., 2020). Therefore, sophisticated mechanisms of tissue and cell-specific proteostatic pathways to adapt to changing conditions are fundamental to prevent protein aggregation, inclusion body formation, and ultimately cell death.

In this study, we demonstrated that the loss of *Ddi2* function in mice led to mid-late gestation embryonic lethality. DDI2 is an aspartic protease that has been recently shown to proteolytically activate the highly polyubiquitylated transcription factor TCF11/NRF1 (NFE2L1) (Dirac-Svejstrup et al., 2020; Koizumi et al., 2016; Siva et al., 2016; Yip et al., 2020). The NGLY1-p97/VCP-TCF11/NRF1-DDI2 axis represents a major stress adaptation pathway activated upon proteasome impairment to evade apoptosis and promote survival of the cell (Radhakrishnan et al., 2010; Steffen et al., 2010; Vangala et al., 2016). Whereas gene knock-outs of *Ngly1* (Fujihira et al., 2017), *Nfe2l1* (Chan et al., 1998), and proteasomal subunits (Sakao et al., 2000) have been described in detail and in mice result in embryonic lethality, nothing was known about the physiological consequences of *Ddi2* dysfunction in the context of the whole organism. Therefore, the importance of DDI2 compared to other components of the pathway has not been clear. Both the full knock-out and the protease defective mouse models described here exhibit homozygous embryonic lethality in the mid-late gestation period with severe developmental failure. Detailed expression analysis using various methods revealed developmental defects in tissues with high levels of DDI2 expression, mainly in the craniofacial region, limb buds and heart (Figure 1 and S1C-E). Interestingly, NRF1-deficient mice had no obvious defects in development prior to death, but suffered from anemia as a result of abnormal fetal liver erythropoiesis (Chan et al., 1998). We were unable to detect differences in hematopoiesis at E9.5 and E10.5 in *Ddi2^ex6-/-^* embryos (data not shown).

Analysis of potential transcription factor binding sites in the *DDI2* promoter region revealed hits for the FOX (forkhead box) and E2F families of transcription factors, indicating that expression of DDI2 is dependent on the cell cycle as well as developmental signaling pathways. This is also supported by our gene expression analysis (GeneHancer database http://genome.ucsc.edu/, unpublished results) (Kent et al., 2002). Specific expression of *Ddi2* at around E10.5 is similar to the previously described expression profile of the DDI2 substrate *Ne2lf1* (NRF1) (Gray et al., 2004). A second substrate identified for DDI2 is NRF3 (Chowdhury et al., 2017). Expression of NRF3 at E10.5 was also detected in the forelimb buds, but was low in other part of the developing embryos (Lewandowski et al., 2015). Interestingly, expression of *Nfe2l3* does not appear to be affected by DDI2 deficiency in the mid to late gestation period (Figure 4A). Expression of *Ddi2* mRNA starting at E13.5 and continuing in mice up to 9 months of age had been previously studied only in mouse brain and its level of expression did not fluctuate. This differs from the dramatic increase of *Ddi1* mRNA at E16.5, which was attributed to a DDI1-specific role in development of the brain (Ramirez et al., 2018). We hypothesized that DDI1 could compensate for a deficiency in DDI2, as is the case in DNA damage response (Kottemann et al., 2018; Svoboda et al., 2019). However, we did not detect an increase in expression of *Ddi1* mRNA in the *Ddi2^ex6-/-^* embryo by qRT-PCR during the three crucial developmental stages prior to death (Figure S4B). Moreover, the expression pattern of the two homologs is distinct (Ramirez et al., 2018; Yousaf et al., 2020). Thus, the two DDI homologues cannot compensate for one another.

Interestingly, the expression patterns of several key members of this stress response axis (NRF1/NFE2L1, ATF4) are similar to that of *Ddi2* (Figure 1F, G, H - pharyngeal arches, limb buds) throughout embryonic stages E9.5 – E11.5, from which the last stage turned out to be the decision point for our DDI2 deficient mouse models (Figure 1C, D). Note that while the upstream regulators of these stress response pathways (Ngly1, proteasome) belong to the essential genes that exhibit embryonic lethality (Chan et al., 1998; Fujihira et al., 2017; Yokoi and Hanaoka, 2017), knock-out mouse models of the downstream regulators such as ATF4, CHOP, STAT1 or STAT3 typically die during postnatal development (Ariyama et al., 2007; Masuoka and Townes, 2002; Tanaka and Chiba, 1998). In this context it is important to note that proper stem cell function as well as signaling pathways of development and differentiation strongly rely on a functional UPS. It is also well established that signaling pathways such as Wnt, Notch and Hedgehog require the timely degradation of modulators to activate down-stream transcription factors (Baloghova et al., 2019; Boukhalfa et al., 2019; Gerhardt et al., 2016; Hamilton and Zito, 2013; Muller-Newen et al., 2017). In conclusion, the developmental defects of DDI2-deficient mice we observed can be most likely attributed to proteasome dysfunction (Figure 2 and 3), while other compensating mechanisms such as NRF2 or UPR are induced (Figure 5) but do not appear to compensate for DDI2-deficiency. Of note, our UPR data for DDI2-deficient embryos and the strong induction of CHOP at time points at which embryonic cell death occurs (Figure 3) support this notion and point to induction of cell death due to unresolved proteotoxic stress.

Upregulation of alternative proteasome isoforms and regulators such as immunoproteasomes and PA28/11S can help to maintain the proteostatic potential of cells and tissues during inflammation (Ebstein et al., 2013; Seifert et al., 2010). In the case of the DDI2-deficient cells and embryos analyzed in this study, impaired proteasome function also induced such compensatory mechanisms (Figures 2 and 3). However, these (typically IFN-regulated) proteins were unable to resolve the DDI2-deficiency-induced proteotoxic crisis. Moreover, PA28 has been shown to be a trancriptional target of NRF2 (Jung et al., 2014) indicating that upregulation relies on both IFN signaling and NRF2.

Although knocking out key members of the NGLY1-p97/VCP-TCF11/NRF1-DDI2-proteasome pathway resulted in embryonic lethality, successful ablation in cells was previously demonstrated for DDI2 (Koizumi et al., 2016; Northrop et al., 2020), NGLY1 (Yang et al., 2018), and NRF1 (Chan et al., 1998; Radhakrishnan et al., 2010). The difference between the embryonic lethality of DDI2^exon6-/-^ on the one hand and the almost normally dividing DDI2 KO cells on the other was intriguing, especially since induction of the PERK-arm of the UPR as well as ATF4 induction has been observed for both DDI2-deficient embryos and cells (Figures 5 and 6). Strikingly, we could not detect phosphorylated eIF2α in DDI2-deficient cells, although the eIF2α kinases PERK, GCN2 and PKR were activated (Figure 6 F). In contrast, the inducible subunit of the eIF2α phosphatase GADD34 was upregulated. Mechanistically this suggests that dephosphorylation of eIF2α by GADD34 may overcome translational arrest to produce IFN-β (Dalet et al., 2017). The cycle of eIF2α phosphorylation and dephosphorylation is of particular importance in promoting translational recovery and survival after stress (Bertolotti, 2018; Novoa et al., 2003; Schneider et al., 2020). Proteasome impairment by pharmacological inhibition or by loss of function mutations in genes which encode subunits of the proteasome result in induction of type I IFNs (Brehm and Kruger, 2015; Ebstein et al., 2019; Poli et al., 2018; Studencka-Turski et al., 2019). Moreover, NGLY1 impairment in murine and human cells causes perturbation of mitochondrial homeostasis and IFN signaling in a manner similar to that observed when the proteasome is impaired (Yang et al., 2018). Our data for DDI2-deficient cells show that fulminant proteotoxic stress leads to induction of a type I IFN signature (Figure 7) and IFN signaling. Subsets of differentially upregulated ISGs have been previously observed, and can also be seen in Figure 6D. Upregulation of ISG15, IFIT1, IFI44, IFI44L, OASL, and IFNB is a signature typically observed in interferonopathies such Aicardy-Goutieres-Syndrom as well as proteasome associated autoinflammatory syndromes (PRAAS) (Kim et al., 2018; Rice et al., 2017). Downregulation of Usp18 is consistent with signal amplification, because Usp18 has been identified not only as an isopeptidase for the ubiquitin-like modifier ISG15, but also as a negative regulator of IFN-signaling (Honke et al., 2016). Furthermore, the observed decrease in IFI27, which encodes mitochondrial IFN alpha-inducible protein 27, a protein involved in IFN-induced apoptosis pathways (Gytz et al., 2017; Liu et al., 2014; Rosebeck and Leaman, 2008), indicates that apoptosis signaling is suppressed as is also seen for UPR-driven apoptosis by CHOP (see Figure 5).

The canonical pathway for IFN activation via cell autonomous innate immune mechanisms is dependent on pattern recognition receptors as well as the TBK1/ IKKε/ DDX3 kinases (Figure 6A). We observed activation of TBK1 in DDI2-deficient embryos and parental cells treated with BTZ. This indicates that these cells induce the canonical pathway via TBK1 (Figure 7). Acute proteotoxic stress eventually causes cell death. The TBK1-independent activation of ISGs in DDI2 deficient cells was therefore surprising. However, it is known from TBK KO mice that IRF3 can be activated in a TBK-independent manner in response to RNA viral infection. In this pathway TBK and IKKε act redundantly (Balka et al., 2020; Miyahira et al., 2009; Perry et al., 2004; Tsuzuki et al., 2016). Therefore, it is likely here that IRF3 is phosphorylated by IKKε to induce IFNB and other ISGs. It is also known that IRF3 can directly induce expression of ISGs via ISRE elements in promoter regions (Ashley et al., 2019; Csumita et al., 2020; DeFilippis et al., 2006). Upon IFNβ production, JAK/STAT is triggered by autocrine or paracrine signaling-typically by phosphorylation of STAT1 (see Figure 7A). The expected phosphorylation of STAT1 only occurred when acute stress was induced by BTZ treatment of parental cells. Strikingly, we could not detect STAT1 phosphorylation in our DDI2-deficient models. Instead STAT3 was observed in its phosphorylated form (Figure 7B,C). This non-canonical signaling is most likely induced in response to chronic stress, when cells require additional survival mechanisms.

JAK-STAT signaling represents such a survival pathway involved in stemness, the epithelial-mesenchymal transition, and cancerogenesis (Deschenes-Simard et al., 2019; Ganguly et al., 2018; Jin et al., 2016; Lin et al., 2020; Lu et al., 2019). STAT3 in particular has been shown to fine-tune type I IFN responses (Charras et al., 2020; Tsai et al., 2019). This appears to be important to ensure proper development, because type I IFN signaling impairs stem cell function and the potential for differentiation (Eggenberger et al., 2019; Yu et al., 2015). Besides its roles in controlling IFN responses, development, and proliferation, the clinical importance of STAT3 is highlighted by its deleterious oncogenic potential and hyperactivation in various malignancies (Todoric and Karin, 2019). Herein, crosstalk between JAK/STAT3 signaling and proliferation pathways such as MAPK/ERK and PI3K/AKT/mTOR occurs (Rawlings et al., 2004), which represent targets for tumor therapy (Bai et al., 2019). In this context it is important to note that the proteostatic potential of myeloma cells is targeted by proteasome inhibitors which are known in turn to activate the NGLY1-DDI2-TCF11/NRF1 axis. This suggests that DDI2 represents a new drug target for multiple myeloma and myeloproliferative neoplasms (Fassmannová et al., 2020; Gu et al., 2020; Steffen et al., 2010). Consistent with this idea, EGFR/JAK1/STAT3 signaling contributes to BTZ resistance in multiple myelomas via induction of immunoproteasomes (Zhang et al., 2016).

In conclusion, we identified DDI2 as an important regulator of proliferative, developmental, and immune signaling with implications for cancerogenesis. These findings pave the way to further explore the mechanisms of DDI2 function and ultimately elucidate the relationship between proteasome impairment and innate immune signaling. In this context the NGLY1-DDI2-TCF11/NRF1 axis and DDI2 itself represent important drug targets. On one hand, inhibition of this pathway with new small molecules may contribute to novel chemotherapeutic strategies in cancer treatment. On the other hand, activators of this pathway are urgently required for conditions with proteasome impairment such as neurodegeneration and other proteinopathies.

## ACKNOWLEDGEMENTS

This work was supported by the Ministry of Education, Youth and Sports of the Czech Republic within the National Sustainability Program II [Project BIOCEV-FAR LQ1604] and by the project “BIOCEV” (CZ.1.05/1.1.00/02.0109). The work was further supported by the Charles University Grant Agency (grant #1294317) and the European Regional Development Fund; OP RDE; Project “Chemical biology for drugging undruggable targets (ChemBioDrug)” (No. CZ.02.1.01/0.0/16_019/0000729). This work was also supported by the German Research Foundation SFBTR 186/ A13 to EK.

The authors thank Edward Curtis for language editing. Special thanks go to Karolína Šrámková, Jana Starková, Jaroslav Kurfürst and Zuzana Kružíková for excellent technical assistance. The authors also thank Dr. Václav Veverka and Michal Svoboda for help with evaluation of NMR and DSF experiments and help with DDI2 mutagenesis, and to Dr. Vendula Novosadová (IMG Prague) for help with statistical analysis.

## AUTHORS CONTRIBUTIONS

Conceptualization, K.G.S., J.K., R.S. and E.K.; Study design, M.S., J.P., P.K., K.G.S, and E.K; Methodology, M.S., S.H.M., M.P., J.P., P.K., F.S., K.CH., K.G.S. and E.K.; Investigation, M.S., S.H.M., M.P., F.S., P.K. and K.CH.; Writing – Original Draft, M.S. and S.H.M.; Writing – K.G.S. and E.K.; Review & Editing, E.K. and K.G.S.; Funding Acquisition, R.S., J.K., E.K. and K.G.S.; Resources, R.S., J.K., E.K. and K.G.S.; Supervision, K.G.S. and E.K.

## STAR METHODS

### RESOURCE AVAILABILITY

#### Lead contact

Further information and requests for resources and reagents should be directed to and will be fulfilled by the Lead Contact, Klara Grantz Saskova.

#### Material Availability

Plasmids will be provided upon request from Klara Grantz Saskova.

Cell lines will be provided upon request and MTA signing from Elke Krüger and Klara Grantz Saskova.

The mouse line *Ddi2^KO^(C57BL/6NCrl-Ddi2^tm1b(EUCOMM)Hmgu^/Ph*) used in this study was generated at the CCP-IMG CAS (Prague, Czech Republic). It has been deposited in the EMMA repository and is available for order via IMPC using ID EM:12340 (https://www.infrafrontier.eu).

The *Ddi2^PD^* mouse line (*C57BL/6NCrl-Ddi2^em1^/Ph*) used in this study was also generated at the CCP-IMG CAS and will be deposited in the EMMA repository upon review process.

Both mouse lines are available upon MTA signing from the laboratory of Jan Konvalinka, IOCB Prague.

#### Data and code availability

Phenotyping data of the *Ddi2^tm1b^* strain (*C57BL/6NCrl-Ddi2^tm1b(EUCOMM)Hmgu^/Ph*) strain has been deposited at https://www.mousephenotype.org/data/genes/MGI:1917244.

Original data used in the figures of the paper will be available at Mendelay Data.

### EXPERIMENTAL MODEL AND SUBJECT DETAILS

#### Cell Culture and Inhibitors

EAhy926 cells (RRID:CVCL_3901) were purchased from ATCC (CRL-2922™). Generation of *DDI2* knock-out EAhy926 cells was previously described (Nowak K. et al., 2018). Cells were cultured in Iscove’s Modified Dulbecco’s Medium (IMDM; PAN-Biotech) supplemented with 10% v/v fetal bovine serum (FBS; PANBiotech), 100 U/ml penicillin and 100 μg/ml streptomycin (PAN-Biotech). Cells were grown at 37°C, 5% CO2 and 95% humidity and routinely passaged 2 times a week.

Proteasomal activity was inhibited using 50 nM water-soluble bortezomib (BTZ; Velcade, Hexal) for the indicated period of time.

HEK293-TetOff-A2 cells were created in the laboratory of Jan Konvalinka (Tykvart et al., 2015). Cells were cultured in IMDM complete medium (Thermo Fisher Scientific), 10% FBS and 40 mM L-glutamine (Sigma-Aldrich) at 37°C, 5% CO2 and 95% humidity and routinely passaged 2 times a week.

#### Animal models

*C57BL/6NCrl* mouse strain animals used for either colony management or experiments were purchased from animal facility of the Institute of Molecular Genetics of the Czech Academy of Science (IMG CAS, Prague, Czech Republic).

The *Ddi2^KO^* strain (*C57BL/6NCrl-Ddi2^tm1b(EUCOMM)Hmgu^/Ph*, RRID: IMSR_EM:12340) originates from the ES cell clone HEPD0660_5_E02 of The European Conditional Mouse Mutagenesis Program (EUCOMM). It was generated for this study by the CCP-IMG CAS (Prague, Czech Republic) by crossing *Gt(ROSA)26Sor^tm1(ACTB-cre,-EGFP)Ics^*) and *Ddi2^tm1a^ (C57BL/6NCrl-Ddi2^tm1a(EUCOMM)Hmgu^/Ph*) mice. The double-heterozygous offspring obtained in this cross were then crossed to *C57BL/6NCrl* wild-type mice and further maintained in this background.

The *Ddi2^PD^* strain (*C57BL/6NCrl-Ddi2^em1^/Ph*) was generated in this study by TALEN-mediated genome editing and maintained in *C57BL/6Ncrl* background.

All mice were housed in IVC cages in the SPF animal facility at the Institute of Molecular Genetics of the Czech Academy of Science (Prague, Czech Republic). All work with mice was approved by the Animal Care Committee of the IMG CAS according to institutional and national guidelines of the Czech Central Commission for Animal Welfare and in accordance with European directive 2010/63/EU.

## METHOD DETAILS

### Generation and characterization of mouse lines

#### Generation of the *C57BL/6NCrl-Ddi2^PD^* mouse line

Exon 6 of the *Ddi2* gene was excised using TAL Effector Nucleotide Targeter 2.0 (Cermak T. et al., 2011, Doyle E. L. et al., 2012). Briefly, TALENs were assembled using the Golden Gate Cloning system (Cermak T., et al., 2011), cloned into the ELD-KKR backbone plasmid (Flemr M. et al., 2013) and transcribed into mRNAs using the mMESSAGE mMACHINE T7 kit (Thermo Fisher Scientific). TALEN mRNAs were polyadenylated with the Poly(A) Tailing kit (Thermo Fisher Scientific), purified using the RNeasy mini kit (QIAGEN) and microinjected into male nucleoli of zygotes isolated from *C57BL/6N* mice as previously described in Kasparek P. et al., 2014. For the off-target control, the F1 generation mice used for colony establishment were screened for twelve possible TALEN off-target sites on chromosome 4 predicted by the TAL Effector Nucleotide Targeter 2.0 (Doyle E. L., et al., 2012) by PCR amplification and sequencing. Sequences of TALENs are provided in Supplementary Table S1. Sequences of primers used in the off-target screen are provided in Table S2.

#### Genotyping of animals

Mice and embryos were genotyped from a gDNA template isolated from the tail-tip, ear or yolk sac by overnight incubation at 55°C with DirectPCR Lysis Reagent (Viagen) and Proteinase K (New England Biolabs). PCR reactions were performed using the Mouse Direct PCR Kit (Bimake). Primers Ddi2F and Ddi2R were used for genotyping of *Ddi2^PD^* strain samples. In cases of inefficient amplification during the first round of PCR, mainly when DNA was purified from small bits of yolk sac tissue, two rounds of nested PCR were added to the genotyping procedure. In Nested PCR 1, primers in closer proximity to exon6 were used (Ddi2 nested F, Ddi2 long R). Nested PCR 2 was designed with a reverse primer (Ddi2 IN R) inside exon6. Primers LacZ F, Ddi2^tm1b^ WT F and Ddi2^tm1b^ RV were used for genotyping of the *Ddi2^tm1b^* strain. Primer sequences are listed in Table S3 in the Supplemental Information.

#### Phenotyping of adult mice

Phenotyping of adult heterozygous mice of both strains was performed in collaboration with the CCP-IMG CAS according to the International Mouse Phenotyping Consortium (IMPC) pipeline workflow and standard operating procedures (available at https://www.mousephenotype.org/).

A *Ddi2^tm1b^* mouse cohort (7 *Ddi2^+/-^* males and 8 *Ddi2^+/-^* females) was analyzed with respect to body composition and weight, performance in behavioral, cardiovascular and lung function tests, hematology and biochemistry, glucose metabolism (IpGTT), gross pathology, and histology after termination. The results were compared to those of a *C57BL/6NCrl* reference cohort housed at the IMG CAS.

*Ddi2^ex6+/-^* mice (8 of each sex) were studied for glucose metabolism (IpGTT), hematology, biochemistry, gross pathology and histology after termination. The results for *Ddi2^ex6+/-^* mice were compared to values from a cohort of *Ddi2^ex6+/+^* mice of the same size and a *C57BL/6NCrl* cohort housed at IMG CAS.

#### MicroCT

Embryos of the *Ddi2*-deficient strains and the control strain *C57BL/6NCrl* were harvested at stages E10.5 and E11.5, fixed in 4% PFA for 3 days and immersed in the contrasting agent 1% phosphotungstic acid for 1 week. Contrasted specimens were embedded in 2.5% low gelling temperature agarose dissolved in water. Scans were performed using a SkyScan 1272 high-resolution microCT (Bruker) with the resolution set to 1.2 μm.

### Biophysical characterization of DDI2^WT^ and Ddi2^PD^ proteins

#### Cloning, recombinant protein expression and purification

Murine DDI2^WT^ (residues 1-399) and *DDI2^PD^* (Δ256-294) protein coding sequences were amplified by PCR from cDNA acquired from embryonal tissue. RNA isolation and reverse transcription were performed as described in the STAR METHODS section *“Ddi2* expression analysis in mouse embryos using qRT-PCR.” Constructs encoding the DDI2^WT^ and *DDI2^PD^* proteins were cloned into p905 (gift from Pavlína Řezáčová, IOCB CAS, Prague) and pTreTight (Clontech) expression vectors.

p905 bacterial expression vectors encoding DDI2^WT^ and *DDI2^PD^* were expressed in BL21(DE3)RIL host cells (Novagen). These proteins, which are in frame with an N-terminal histidine tag, were purified by two rounds of nickel affinity chromatography using Ni-NTA agarose (QIAGEN), with overnight cleavage by N-terminally His-tagged TEV protease (expressed in our laboratory) in-between. The later flow-through fraction containing the desired protein was further purified by size-exclusion chromatography in 50 mM sodium phosphate buffer pH 7.4, 0.5 % (v/v) glycerol by FPLC (ÄKTA explorer, Amersham Pharmacia Biotech) using a Superdex™ 200pg 16/60 FPLC column (GE Healthcare). Final protein fractions were analyzed by SDS-PAGE electrophoresis.

#### 1D NMR

The ^1^H HSQC spectra of DDI2^WT^ and *DDI2^PD^* were acquired using 350 μl of 50 μM protein samples in 50 mM phosphate buffer at pH 7.4 with 0.5% glycerol at 25°C on a 600 MHz Bruker Avance spectrometer (Bruker BioSpin GmbH) equipped with a triple resonance (^15^N, ^13^C, ^1^H) cryoprobe. Spectra were processed using Topspin 3.2 (Bruker).

#### DSF

DSF experiments were performed using 20 μM final concentration of *DDI2^PD^* and DDI2^WT^ proteins in 50 mM phosphate buffer pH 7.4, 0.5% glycerol with 5000x diluted SYPRO^®^ Orange protein gel stain (Sigma-Aldrich) in total reaction volume 25 μl. The thermofluor assay and calculation of melting temperatures were performed on a LightCycler^®^ 480 II using LightCycler^®^ 480 Software (Roche).

### Experimental procedures with mouse tissue

#### B-galactosidase activity detection

For mapping of *Ddi2* expression using the B-galactosidase activity detection method, *Ddi2^+/-^* adult mice of both sexes (age: 16 weeks) and embryos at the embryonal stages E9.5, E10.5, E12.5, and E14.5 were harvested. Embryos and adult tissue used for whole mount staining were fixed in 4% PFA and thoroughly washed in 1 M phosphate buffer pH 7.5, 0.5 M EGTA, 0.01% sodium deoxycholate, 2 mM MgCl_2_, and 0.02% Nonidet P-40. They were then stained overnight at 37°C in the dark in X-gal staining solution containing 0.1 M phosphate buffer pH 7.5, 0.02% Nonidet P-40, 0.01% sodium deoxycholate, 5 mM potassium ferricyanide, 5 mM potassium ferrocyanide, 2 mM MgCl_2_, and 1mg/ml X-Gal (Thermo Fisher Scientific), rinsed in PBS and post-fixed in 4% PFA prior to imaging.

Embryos used for sectioning were embedded in 30% sucrose/PBS overnight and frozen in OCT prior to cryo-sectioning (10 μm sagittal sections) according to standard operating protocols of the Czech Centre for Phenogenomics. Following washing and staining, procedures were performed as for whole mount staining. After post-fixation in 4% PFA, slides were washed in PBS, counter-stained with Nuclear Fast Red (Sigma-Aldrich) and mounted in Aquatex (Merck Millipore). Imaging of sections was performed using a Zeiss AxioImager Z2 and a Zeiss AxioScan Z1. Zeiss AxioZoom with Apotome module macroscope was used for whole mount imaging.

#### RNA *In situ* hybridization

Specific types of tissues from adult mice and whole embryos of the *C57BL/6NCrl* strain at the E9.5, E10.5 and E11.5 stages were harvested and processed as either paraffin sections or as whole-mount samples. Adult tissues were fixed in 4% PFA, embedded in paraffin, and sectioned at 7μm. Whole-mount embryos were fixed in 4% PFA and frozen in methanol at −20°C, which was followed by tissue hydration, proteinase K treatment, acetylation, and pre-hybridization as previously described (Wilkinson and Nieto, 1993). Solutions were prepared with RNAse free water. Hybridization on sections was performed overnight at 70°C with DIG-labelled probes in hybridization buffer (1.25X saline-sodium citrate, pH 7.0, 50% formamide, 0.1% Tween, 20, 100X Denhardt’s solution, heparin (50 μg/ml), tRNA (50 μg/ml), and salmon sperm DNA (50 μg/ml)). Hybridization on whole-mount embryos was performed overnight at 69°C with DIG-labelled probes in hybridization buffer (1.3X saline-sodium citrate, pH 7.0, 50% formamide, 0.2% Tween-20-5mM EDTA, 0.5% CHAPS, heparin (100 μg/ml), and tRNA (50 μg/ml)). Digoxygenin-labeled RNA probes (DIG RNA labeling kit, Roche) were generated by *in vitro* transcription from a PCR-amplified fragment of murine *Ddi2*. The sequence of the *Ddi2* antisense probe is shown in Table S5 in the Supplements. An anti-DIG antibody conjugated to alkaline phosphatase and BM purple alkaline phosphatase substrate precipitating solution (Merck Roche) were used for staining. All samples were postfixed with 4% PFA. Imaging of whole-mount samples was performed using a Zeiss AxioZoom with an Apotome module macroscope and sections were imaged using a Zeiss Axio Imager 2.

#### *Ddi2* expression analysis in mouse embryos using qRT-PCR

RNA was isolated from frozen RNA later^®^ soaked tissue from the embryo from either *C57BL/6NCrl* or *Ddi2^PD^* strains using RNeasy plus Micro and Mini Kits (QIAGEN). Samples with RIN values above 7 in the Bioanalyzer RNA 6000 Nano assay (Agilent) were used for the synthesis of complementary DNA and RT controls using a reverse transcription TATAA GrandScript cDNA Synthesis Kit (TATAA Biocenter). qRT-PCR was performed using a LightCycler^®^ 480 (Roche) and TATAA SYBR^®^ GrandMaster^®^ Mix (TATAA Biocenter) by the Gene Core facility at the IBT CAS. Raw data were preprocessed using the GenEx− program (MultiD Analyses AB). Primers are listed in Table S4 in the Supplemental Information.

#### Tissue lysis of *Ddi2^PD^* embryos for immunoblotting

Frozen embryos were homogenized in TSDG buffer (10 mM Tris/HCl pH 7.5, 25 mM KCl, 10 mM NaCl, 1 mM MgCl_2_, 0.1 mM EDTA, 0.5 mM DTT, 0.2 mM ATP, and 10 % (v/v) glycerol) for 30 s at a frequency of 15 s^-1^ using a TissueLyser II (QIAGEN). As the tissue homogenates were to be used for both native and SDS PAGE, 3/5 of the sample was transferred to a new micro test tube and different inhibitors (10 mM NEM, 10 mM NaF, 1 mM Na_3_VO_4_, 2 mM Na_4_P_2_O_7_, 10 μM MG132, and 1x cOmplete™ Mini protease inhibitor cocktail (EDTA free, Merck Roche)) were added. Both homogenates were then lysed by four freezing and thawing cycles in liquid nitrogen. Lysates were centrifuged at 17,000 x g for 15 min at 4°C. The pellets were stored at −80°C until further analysis. Protein concentrations were determined using the Bradford reagent (0.01% (w/v) Coomassie brilliant blue G250, 4.8% (v/v) ethanol, and 8.5% phosphoric acid).

For detection of ubiquitinated proteins and transcription factors in the insoluble fraction, pellets of tissue lysate were resuspended in PBS with 5000 U/mL of *S. marcescens* endonuclease (Protean). After incubating for 10 min at room temperature while shaking, samples were denatured by adding SDS-PAGE sample buffer (final concentration: 62.5 mM Tris-HCl pH 6.8, 2% (w/v) SDS, 10% (v/v) glycerol, 1% β-mercaptoethanol, and 0,01% (w/v) bromophenol blue). The amount of lysis buffer added was calculated using the measured embryo weight, and 33 μL of buffer was added per 15 mg of embryo.

### Experimental procedures with human cell lines

#### Cell extract preparation

For the preparation of whole cell extracts for immunoblotting, EAhy926 cells were grown in 35 mm or 60 mm dishes to 80-90% confluency, rinsed with ice-cold PBS, and lysed on the plate in RIPA lysis buffer (150 mM NaCl, 50 mM Tris-HCL (pH 8), 5 mM Na-EDTA (pH 8), 0.5 % (v/v) NP-40, 0.5 % (v/v) Triton X-100, 0.5 % sodium dodecyl sulfate, 1 mM Na_3_VO_4_, 10 mM NaF, 2 mM Na_4_P_2_O_7_, 10 μM MG132, 10 mM NEM and 1x cOmplete™ Mini protease inhibitor cocktail (EDTA free, Merck Roche)) for 5 min on ice. The cells were then scraped off the plate, lysates were collected in micro test tubes and kept on ice for 15 min followed by freezing at −80°C. Debris was removed by 15 min of centrifugation at 17,000 x g and 4°C.

For the preparation of cell extracts for native gel analysis, EAhy926 cells were grown in 35 mm dishes to 80-90% confluency. Cell pellets were lysed in TSDG buffer (10 mM Tris/HCl pH 7.5, 25 mM KCl, 10 mM NaCl, 1 mM MgCl_2_, 0.1 mM EDTA, 0.5 mM DTT, 0.2 mM ATP, and 10 % (v/v) glycerol) by four freezing and thawing cycles in liquid nitrogen. Lysates were centrifuged at 17,000 x g for 15 min at 4°C and the supernatant was used for further analysis.

Protein concentrations were determined using the Bradford reagent (0.01% (w/v) Coomassie brilliant blue G250, 4.8% (v/v) ethanol, and 8.5% phosphoric acid).

#### Cellular fractionation

EAhy926 cells were grown in 100 mm dishes to 80-90% confluency. All centrifugation steps were carried out at 4°C. Cell pellets were resuspended in 200 μL buffer A (20 mM Tris-HCL (pH 7.5), 10 mM KCl, 1.5 mM MgCl_2_, 10% glycerol, 1 mM Na_3_VO_4_, 10 mM NaF, 10 μM MG132 and 1x cOmplete™ Mini protease inhibitor cocktail (EDTA free, Merck Roche)) and incubated on ice for 15 min before addition of 5 μL of NP-40 (12.5%). Samples were gently mixed immediately and incubated on ice for 5 min. After 10 min of centrifugation at 2,000 x g the supernatant was transferred to a new micro test tube and centrifuged again at 13,000 x g for 10 min. The cleared supernatant (non-nuclear fraction) was transferred to a new micro test tube. The pellet from the first centrifugation step was washed three times with 200 μL buffer A and centrifuged each time at 6,000 x g for 10 min. It was then resuspended in 50 μL buffer C (20 mM Tris-HCL (pH 7.5), 420 mM NaCl, 1.5 mM MgCl_2_, 0.2 mM EDTA, 10% glycerol, 1 mM Na_3_VO_4_, 10 mM NaF, 10 μM MG132, and 1x protease inhibitor cocktail) and incubated on ice for 1 h, during which time the suspension was mixed every 10 min for 15 sec. After 10 min of centrifugation at 13,000 x g the supernatant (nuclear fraction) was transferred to a new micro test tube. The insoluble pellet was mixed with 25 μL buffer E (20 mM Tris-HCL (pH 7.5), 150 mM NaCl, 1.5 mM MgCl_2_, 5 mM CaCl_2_, 10% glycerol, 1 mM Na_3_VO_4_, 10 mM NaF, 10 μM MG132, and 1x protease inhibitor cocktail) and 1 μL Benzonase^®^ nuclease (250 U/μL, Sigma-Aldrich) and incubated on ice for 40 min, during which time the suspension was mixed every 10 min. 25 μL buffer E/1 (buffer E + 2% SDS) was then added. The suspension was incubated at room temperature for 5 min and centrifuged at 13,000 x g for 10 min. The supernatant (chromatin-associated fraction) was transferred to a new micro test tube.

Protein concentrations were determined using the Bradford reagent (0.01% (w/v) Coomassie brilliant blue G250, 4.8% (v/v) ethanol, and 8.5% phosphoric acid). 30 μg total protein of each fraction was used for Western blot analysis.

#### qRT-PCR of human cell cultures

To isolate RNA, EAhy926 cells were grown in 35 mm dishes to 80-90% confluency, rinsed with ice-cold PBS, and lysed on the dish in peqGOLD TriFastTM reagent (VWR Peqlab) for 5 min at room temperature. Lysates were transferred to micro test tubes and, if necessary, stored at −80°C before isolation of RNA according to the manufacturer’s protocol. Complementary DNA was synthesized using 1.5 μg of RNA with random Oligo-dT-Primers (Promega) and M-MLV Reverse Transcriptase (Promega) according to the manufacturer’s instructions. Real-time PCR was performed using Premix Ex Taq− probe (Takara) and TaqMan assays (Applied Biosystems) with a CFX96TM Real-Time System (Bio-Rad).

#### Recombinant expression of mouse DDI2 proteins in human cell lines

Plasmids pTreTight (7.5 μg of plasmid DNA) were transfected into HEK293-TetOff-A2 cells at confluence of approx. 70% with opti-MEM medium (Thermo Fisher Scientific) and 10% (v/v) polyethylenimine (Sigma-Aldrich) and harvested 4, 8, 16, 24, 32, 40 and 48 hours after transfection. Each treatment and time point was performed in triplicate.

#### Transfection with plasmid DNA

EAhy926 *DDI2* KO cells were grown in 35 mm dishes to 70% confluency and transfected with 50 ng of plasmid DNA and 950 ng of control plasmid (1 μg DNA in total) or control plasmid alone using jetPRIME^®^ (Polyplus) according to the manufacturer’s protocol. 21 h after transfection, cells were treated with 500 nM BTZ for 3 hours. Control plasmid pcDNA3.1 was purchased from Invitrogen. DDI2-V5-His (wild type) was previously cloned in the laboratory of Dr. Elke Krüger. Inactive mutant constructs DDI2 D252A (Siva et al., 2016) and DDI2 D252N (generated by site-directed mutagenesis in the laboratory of Dr. Grantz Saskova) were cloned into a pcDNA3.1/V5-His-TOPO vector backbone.

### Protein analysis

#### Western blot analysis

For Western blot analysis, 20-40 μg of embryo tissue lysates with inhibitors, 2-10 μL of insoluble pellet lysate or 10-40 μg of cell extracts were used for sodium dodecyl sulfate polyacrylamide gel electrophoresis (SDS PAGE). To analyze polyubiquitinated proteins, samples were separated on 4-14% Tris-Glycine gradient gels, whereas 10% or 12% Tris-Glycine gels were used for all other experiments. Proteins were transferred onto PVDF membranes (Merck Millipore) by wet electroblotting for 1 h at 100 V using the Mini-PROTEAN Tetra system (Bio-Rad). Equal protein loading was assessed by amido black staining (0.02% (m/v) amido black 10B, 9% (v/v) methanol, and 2% (v/v) acetic acid). Membranes were blocked with 1x ROTI Block (Carl Roth) and incubated overnight with the appropriate primary antibody. Excess antibody was removed by 3 x 5 min washing with 1x TBS-T (0.1% Tween-20). Membranes were then incubated for 30 min with a horseradish peroxidase-linked secondary antibody (anti-rabbit or antimouse IgG, Cell Signaling). The Clarity Western ECL Substrate (Bio-Rad) was used for chemiluminescent detection and membranes were visualized using Fusion FX (Vilber Lourmat) or RP NEW/UV film (CEA). If reprobing with another primary antibody was necessary, membranes were stripped using Restore™ Western Blot Stripping Buffer (Thermo Fisher Scientific) before blocking.

HEK293offA2 cells transfected with pTreTight plasmids were lysed in SDS sample buffer without dye (60 mM Tris pH 6.8, 60 nM SDS, and 0.3 mM β-mercaptoethanol) with cOmplete™ Mini, EDTA-free Protease Inhibitor Cocktail (Merck Roche) and 1 μl of (15x diluted) Benzonase^®^ (Novagen^®^, Merck Millipore) by sonication on ice. Proteins were separated by SDS-PAGE electrophoresis and transferred to nitrocellulose membranes (Bio-Rad) using a wet transfer apparatus (Bio-Rad). Western blot analysis of protein expression was performed using protein-specific primary antibodies (anti-DDI2 (Bethyl) and anti-β-actin (Sigma Aldrich)) and fluorescent secondary antibodies using an Odyssey^®^ CLx Infrared Imaging System (LI-COR Biosciences).

#### Native gel analysis

Fresh embryo lysates without inhibitors or fresh native cell extracts were used for native PAGE analysis. Equal amounts of protein (15 to 20 μg) were mixed with 5x native PAGE sample buffer (final concentration: 50 mM Bis-Tris/HCl pH 6.8, 10% (v/v) glycerol-50 mM NaCl, and 0.01% (w/v) bromophenol blue). Samples were loaded on 3-12% non-denaturing Bis-Tris gels (Invitrogen) and subjected to electrophoresis for 20 h at 45 V (4°C) using the Hoefer miniVE system (SE300-10A-1.0, Hoefer Pharmacia Biotech). The chymotrypsin-like activity of the proteasome was detected by incubating the gel in overlay buffer (20 mM Tris, 5 mM MgCl_2_) containing 100 μM Suc-LLVY-AMC (Bachem) for 30 min at 37°C. Gels were analyzed using the Fusion FX (Vilber Lourmat) with an F-535 Y2 emission filter (530–550 nm). Proteasome complexes were subsequently blotted for 1 h at 200 V on ice onto PVDF membranes (Merck Millipore) and further handled as described above.

### QUANTIFICATION AND STATISTICAL ANALYSIS

Expression of *Ddi2* in *C57BL/6NCrl* embryos was computed relative to expression of the *H2afz* housekeeping gene. In the qRT-PCR screen of 33 genes in *Ddi2^PD^* strain embryos, relative expression of genes was computed using normalization to *Tbp* and *H2afz* housekeeping genes with separately applied ANOVA statistical analysis (p value > 0.05) and a linear mixed-effects model (LMM). Outliers were determined according to Grubbs’s test. Bonferroni correction was applied to evaluate significance in both statistical approaches.

In the qRT-PCR screen of gene expression in EAhy926 cells, expression was calculated relative to expression of *RPLP0* using the unpaired two-sided t test (p value > 0.05). The Grubbs’s outlier test was applied, but no outliers were found. The Holm-Šidák correction (α=5%) was applied to evaluate statistical significance.

Quantification of western blot analysis was performed using ImageJ software. The unpaired two-sided t test was used to evaluate statistical significance (p value > 0.05).

Further statistical details for individual experiments can be found in the figure legends.

### KEY RESOURCES TABLE

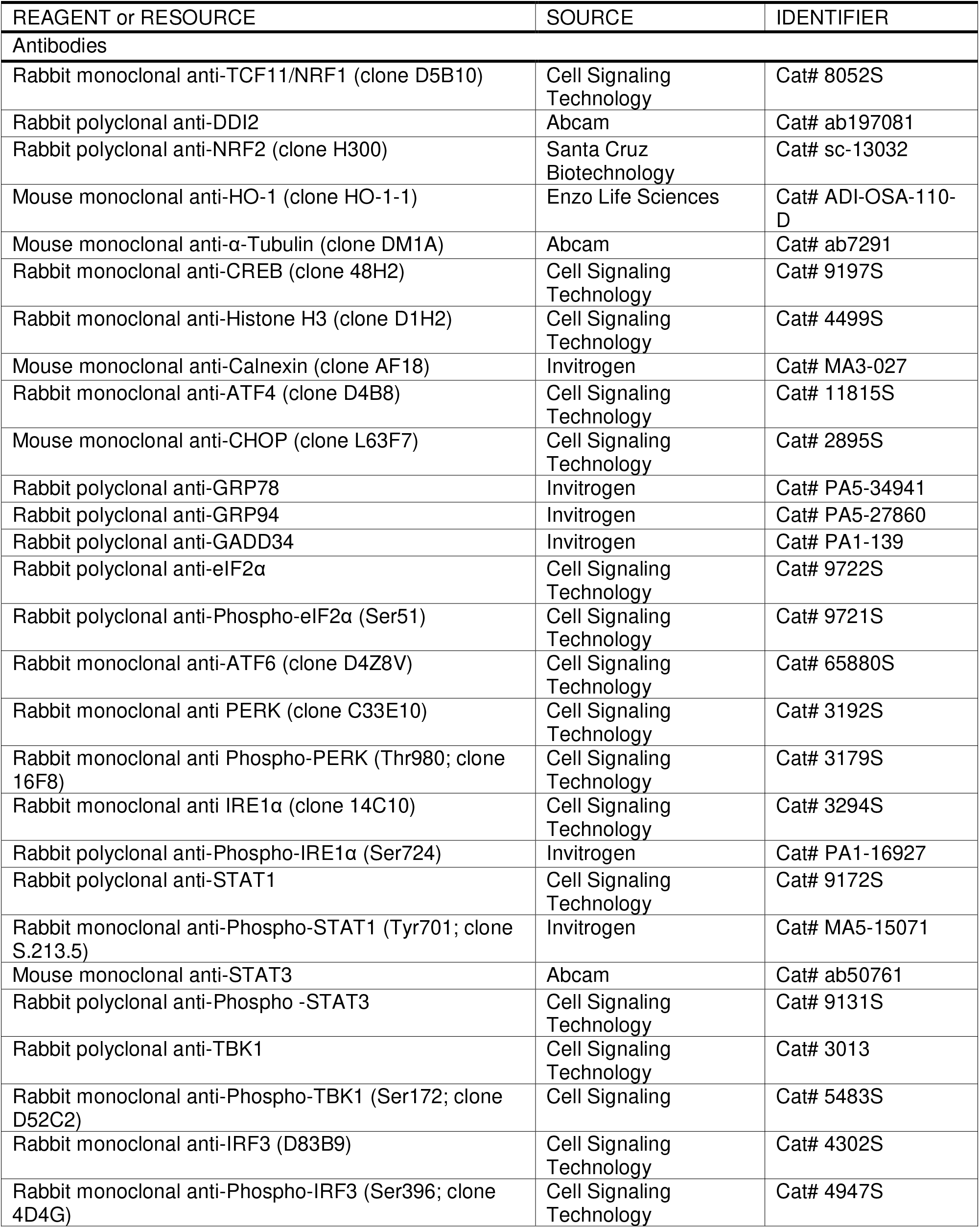
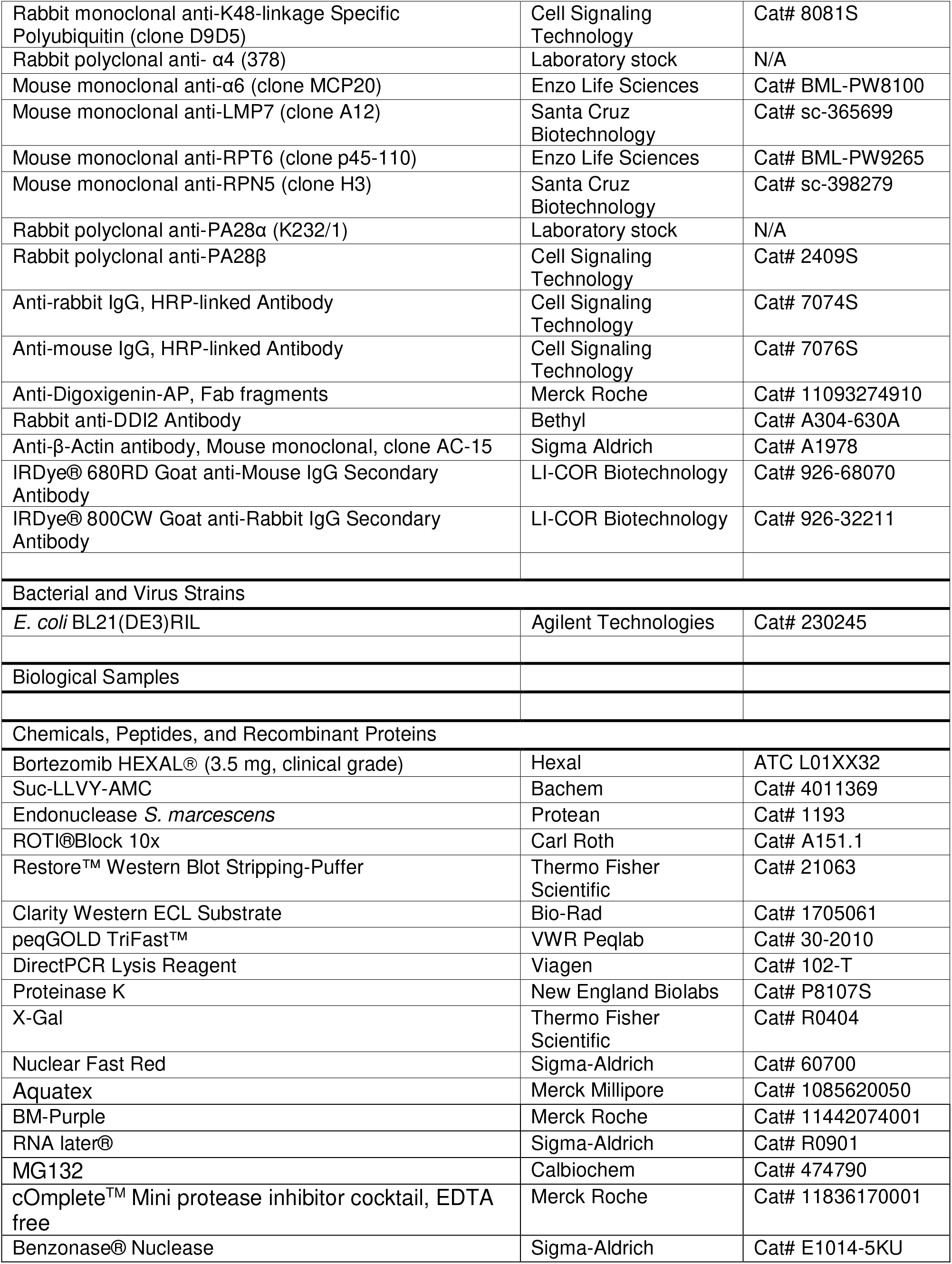
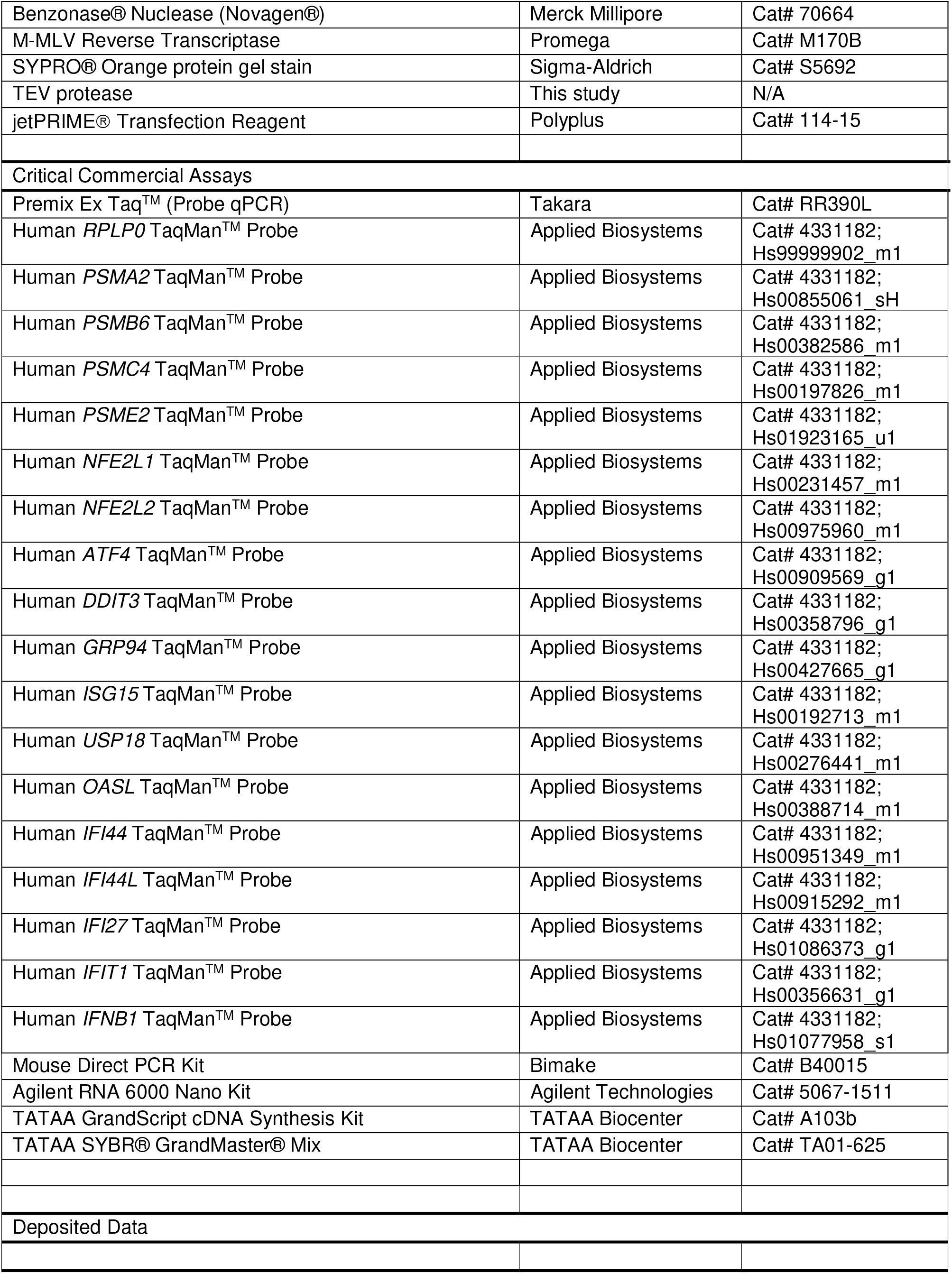
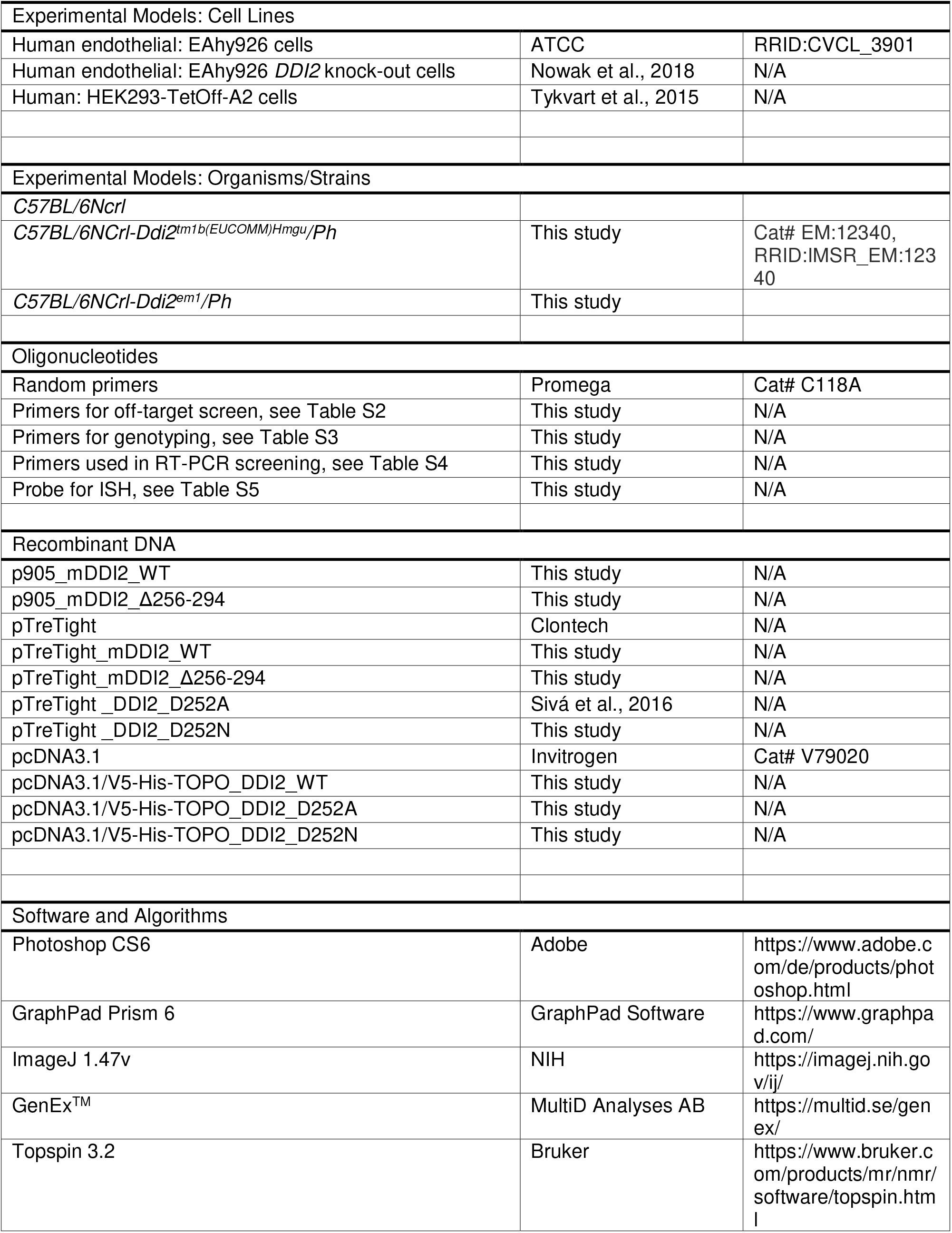
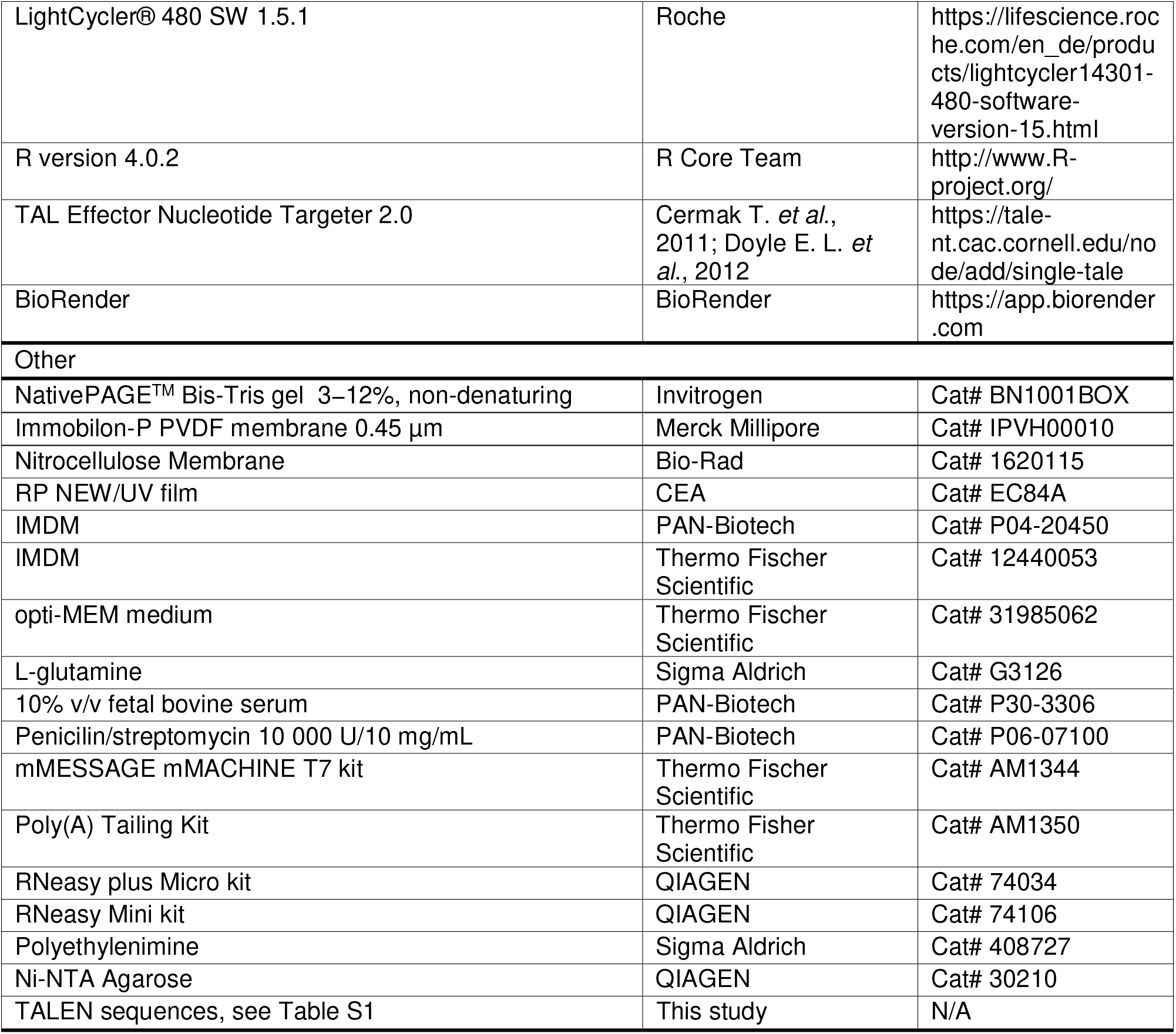

## REFERENCES

Ariyama, Y., Shimizu, H., Satoh, T., Tsuchiya, T., Okada, S., Oyadomari, S., Mori, M., and Mori, M. (2007). Chop-deficient mice showed increased adiposity but no glucose intolerance. Obesity 15, 1647–1656.

Ashley, C.L., Abendroth, A., McSharry, B.P., and Slobedman, B. (2019). Interferon-Independent Upregulation of Interferon-Stimulated Genes during Human Cytomegalovirus Infection is Dependent on IRF3 Expression. Viruses-Basel 11.

Bai, L.C., Zhou, H.B., Xu, R.Q., Zhao, Y.J., Chinnaswamy, K., McEachern, D., Chen, J.Y., Yang, C.Y., Liu, Z.M., Wang, M., et al. (2019). A Potent and Selective Small-Molecule Degrader of STAT3 Achieves Complete Tumor Regression In Vivo. Cancer Cell 36, 498–+.

Balka, K.R., Louis, C., Saunders, T.L., Smith, A.M., Calleja, D.J., D’Silva, D.B., Moghaddas, F., Tailler, M., Lawlor, K.E., Zhan, Y.F., et al. (2020). TBK1 and IKK epsilon Act Redundantly to Mediate STING-Induced NF-kappa B Responses in Myeloid Cells. Cell Reports 31.

Baloghova, N., Lidak, T., and Cermak, L. (2019). Ubiquitin Ligases Involved in the Regulation of Wnt, TGF-beta, and Notch Signaling Pathways and Their Roles in Mouse Development and Homeostasis. Genes (Basel) 10.

Bartelt, A., Widenmaier, S.B., Schlein, C., Johann, K., Goncalves, R.L.S., Eguchi, K., Fischer, A.W., Parlakgul, G., Snyder, N.A., Nguyen, T.B., et al. (2018). Brown adipose tissue thermogenic adaptation requires Nrf1-mediated proteasomal activity. Nat Med 24, 292–+.

Bertolotti, A. (2018). The split protein phosphatase system. Biochem J 475, 3707–3723.

Boukhalfa, A., Miceli, C., Avalos, Y., Morel, E., and Dupont, N. (2019). Interplay between primary cilia, ubiquitin-proteasome system and autophagy. Biochimie 166, 286–292.

Brehm, A., and Kruger, E. (2015). Dysfunction in protein clearance by the proteasome: impact on autoinflammatory diseases. Semin Immunopathol 37, 323–333.

Brehm, A., Liu, Y., Sheikh, A., Marrero, B., Omoyinmi, E., Zhou, Q., Montealegre, G., Biancotto, A., Reinhardt, A., Almeida de Jesus, A., et al. (2015). Additive loss-of-function proteasome subunit mutations in CANDLE/PRAAS patients promote type I IFN production. J Clin Invest 125, 4196–4211.

Cavo, M., Pantani, L., Pezzi, A., Petrucci, M.T., Patriarca, F., Di Raimondo, F., Marzocchi, G., Galli, M., Montefusco, V., Zamagni, E., et al. (2015). Bortezomib-thalidomide-dexamethasone (VTD) is superior to bortezomib-cyclophosphamide-dexamethasone (VCD) as induction therapy prior to autologous stem cell transplantation in multiple myeloma. Leukemia 29, 2429–2431.

Chan, J.Y., Kwong, M., Lu, R., Chang, J., Wang, B., Yen, T.S., and Kan, Y.W. (1998). Targeted disruption of the ubiquitous CNC-bZIP transcription factor, Nrf-1, results in anemia and embryonic lethality in mice. EMBO J 17, 1779–1787.

Charras, A., Arvaniti, P., Le Dantec, C., Arleevskaya, M.I., Zachou, K., Dalekos, G.N., Bordron, A., and Renaudineau, Y. (2020). JAK Inhibitors Suppress Innate Epigenetic Reprogramming: a Promise for Patients with Sjogren’s Syndrome. Clin Rev Allerg Immu 58, 182–193.

Chevillard, G., and Blank, V. (2011). NFE2L3 (NRF3): the Cinderella of the Cap’n’Collar transcription factors. Cell Mol Life Sci 68, 3337–3348.

Chowdhury, A.M.M.A., Katoh, H., Hatanaka, A., Iwanari, H., Nakamura, N., Hamakubo, T., Natsume, T., Waku, T., and Kobayashi, A. (2017). Multiple regulatory mechanisms of the biological function of NRF3 (NFE2L3) control cancer cell proliferation. Sci Rep-Uk 7.

Collins, G.A., and Goldberg, A.L. (2017). The Logic of the 26S Proteasome. Cell 169, 792–806.

Csumita, M., Csermely, A., Horvath, A., Nagy, G., Monori, F., Goczi, L., Orbea, H.A., Reith, W., and Szeles, L. (2020). Specific enhancer selection by IRF3, IRF5 and IRF9 is determined by ISRE half-sites, 5 ‘ and 3 ‘ flanking bases, collaborating transcription factors and the chromatin environment in a combinatorial fashion. Nucleic Acids Res 48, 589–604.

Dalet, A., Arguello, R.J., Combes, A., Spinelli, L., Jaeger, S., Fallet, M., Manh, T.P.V., Mendes, A., Perego, J., Reverendo, M., et al. (2017). Protein synthesis inhibition and GADD34 control IFN-beta heterogeneous expression in response to dsRNA. Embo Journal 36, 761–782.

DeFilippis, V.R., Robinson, B., Keck, T.M., Hansen, S.G., Nelson, J.A., and Fruh, K.J. (2006). Interferon regulatory factor 3 is necessary for induction of antiviral genes during human cytomegalovirus infection. Journal of Virology 80, 1032–1037.

Deschenes-Simard, X., Parisotto, M., Rowell, M.C., Le Calve, B., Igelmann, S., Moineau-Vallee, K., Saint-Germain, E., Kalegari, P., Bourdeau, V., Kottakis, F., et al. (2019). Circumventing senescence is associated with stem cell properties and metformin sensitivity. Aging Cell 18.

Dirac-Svejstrup, A.B., Walker, J., Faull, P., Encheva, V., Akimov, V., Puglia, M., Perkins, D., Kumper, S., Hunjan, S.S., Blagoev, B., et al. (2020). DDI2 Is a Ubiquitin-Directed Endoprotease Responsible for Cleavage of Transcription Factor NRF1. Molecular Cell 79, 332–+.

Ebstein, F., Poli Harlowe, M.C., Studencka-Turski, M., and Kruger, E. (2019). Contribution of the Unfolded Protein Response (UPR) to the Pathogenesis of Proteasome-Associated Autoinflammatory Syndromes (PRAAS). Front Immunol 10, 2756.

Ebstein, F., Voigt, A., Lange, N., Warnatsch, A., Schroter, F., Prozorovski, T., Kuckelkorn, U., Aktas, O., Seifert, U., Kloetzel, P.M., et al. (2013). Immunoproteasomes are important for proteostasis in immune responses. Cell 152, 935–937.

Eggenberger, J., Blanco-Melo, D., Panis, M., Brennand, K.J., and tenOever, B.R. (2019). Type I interferon response impairs differentiation potential of pluripotent stem cells. Proc Natl Acad Sci U S A 116, 1384–1393.

Fassmannová, D., Sedlák, F., Sedláček, J., Špička, I., and Grantz Šašková, K. (2020). Nelfinavir Inhibits the TCF11/Nrf1-Mediated Proteasome Recovery Pathway in Multiple Myeloma. Cancers 12.

Finley, D. (2009). Recognition and Processing of Ubiquitin-Protein Conjugates by the Proteasome. Annu Rev Biochem 78, 477–513.

Fujihira, H., Masahara-Negishi, Y., Tamura, M., Huang, C., Harada, Y., Wakana, S., Takakura, D., Kawasaki, N., Taniguchi, N., Kondoh, G., et al. (2017). Lethality of mice bearing a knockout of the Ngly1-gene is partially rescued by the additional deletion of the Engase gene. PLoS Genet 13, e1006696.

Ganguly, D., Sims, M., Cai, C., Fan, M.Y., and Pfeffer, L.M. (2018). Chromatin Remodeling Factor BRG1 Regulates Stemness and Chemosensitivity of Glioma Initiating Cells. Stem Cells 36, 1804–1815.

Garcia-Prat, L., Sousa-Victor, P., and Munoz-Canoves, P. (2017). Proteostatic and Metabolic Control of Stemness. Cell Stem Cell 20, 593–608.

Gerhardt, C., Wiegering, A., Leu, T., and Ruther, U. (2016). Control of Hedgehog Signalling by the Cilia-Regulated Proteasome. J Dev Biol 4.

Gray, P.A., Fu, H., Luo, P., Zhao, Q., Yu, J., Ferrari, A., Tenzen, T., Yuk, D.I., Tsung, E.F., Cai, Z.H., et al. (2004). Mouse brain organization revealed through direct genome-scale TF expression analysis. Science 306, 2255–2257.

Gu, Y., Wang, X., Wang, Y., Wang, Y., Li, J., and Yu, F.X. (2020). Nelfinavir inhibits human DDI2 and potentiates cytotoxicity of proteasome inhibitors. Cell Signal 75, 109775.

Gytz, H., Hansen, M.F., Skovbjerg, S., Kristensen, A.C., Horlyck, S., Jensen, M.B., Fredborg, M., Markert, L.D., McMillan, N.A., Christensen, E.I., et al. (2017). Apoptotic properties of the type 1 interferon induced family of human mitochondrial membrane ISG12 proteins. Biol Cell 109, 94–112.

Hamilton, A.M., and Zito, K. (2013). Breaking It Down: The Ubiquitin Proteasome System in Neuronal Morphogenesis. Neural Plast 2013.

Hetz, C. (2012). The unfolded protein response: controlling cell fate decisions under ER stress and beyond. Nat Rev Mol Cell Biol 13, 89–102.

Hetz, C., Zhang, K.Z., and Kaufman, R.J. (2020). Mechanisms, regulation and functions of the unfolded protein response. Nat Rev Mol Cell Bio 21, 421–438.

Hipp, M.S., Kasturi, P., and Hartl, F.U. (2019). The proteostasis network and its decline in ageing. Nat Rev Mol Cell Bio 20, 421–435.

Hipp, M.S., Park, S.H., and Hartl, F.U. (2014). Proteostasis impairment in protein-misfolding and - aggregation diseases. Trends Cell Biol 24, 506–514.

Honke, N., Shaabani, N., Zhang, D.E., Hardt, C., and Lang, K.S. (2016). Multiple functions of USP18. Cell Death Dis 7.

Jin, J.L., Liu, J., Chen, C., Liu, Z.P., Jiang, C., Chu, H.S., Pan, W.J., Wang, X.B., Zhang, L.Q., Li, B., et al. (2016). The deubiquitinase USP21 maintains the stemness of mouse embryonic stem cells via stabilization of Nanog. Nat Commun 7.

Jung, T., Hohn, A., and Grune, T. (2014). The proteasome and the degradation of oxidized proteins: Part III-Redox regulation of the proteasomal system. Redox Biol 2, 388–394.

Kent, W.J., Sugnet, C.W., Furey, T.S., Roskin, K.M., Pringle, T.H., Zahler, A.M., and Haussler, D. (2002). The human genome browser at UCSC. Genome Res 12, 996–1006.

Kim, H., de Jesus, A.A., Brooks, S.R., Liu, Y., Huang, Y., VanTries, R., Sanchez, G.A.M., Rotman, Y., Gadina, M., and Goldbach-Mansky, R. (2018). Development of a Validated Interferon Score Using NanoString Technology. J Interf Cytok Res 38, 171–185.

Koizumi, S., Irie, T., Hirayama, S., Sakurai, Y., Yashiroda, H., Naguro, I., Ichijo, H., Hamazaki, J., and Murata, S. (2016). The aspartyl protease DDI2 activates Nrf1 to compensate for proteasome dysfunction. Elife 5.

Kottemann, M.C., Conti, B.A., Lach, F.P., and Smogorzewska, A. (2018). Removal of RTF2 from Stalled Replisomes Promotes Maintenance of Genome Integrity. Molecular Cell 69, 24–+.

Kroll-Hermi, A., Ebstein, F., Stoetzel, C., Geoffroy, V., Schaefer, E., Scheidecker, S., Bar, S., Takamiya, M., Kawakami, K., Zieba, B.A., et al. (2020). Proteasome subunit PSMC3 variants cause neurosensory syndrome combining deafness and cataract due to proteotoxic stress. Embo Mol Med.

Kruger, E., and Kloetzel, P.M. (2012). Immunoproteasomes at the interface of innate and adaptive immune responses: two faces of one enzyme. Curr Opin Immunol 24, 77–83.

Leung, L., Kwong, M., Hou, S., Lee, C., and Chan, J.Y. (2003). Deficiency of the Nrf1 and Nrf2 transcription factors results in early embryonic lethality and severe oxidative stress. J Biol Chem 278, 48021–48029.

Lewandowski, J.P., Du, F., Zhang, S.L., Powell, M.B., Falkenstein, K.N., Ji, H.K., and Vokes, S.A. (2015). Spatiotemporal regulation of GLI target genes in the mammalian limb bud. Dev Biol 406, 92–103.

Lin, L., Wang, Y., Sun, B.J., Liu, L.Y., Ying, W.J., Wang, W.J., Zhou, Q.H., Hou, J., Yao, H.L., Hu, L.Y., et al. (2020). The clinical-immunological and genetic features of 12 Chinese patients with STAT3 mutations. Allergy Asthma Cl Im 16.

Ling, S.C., Lau, E.K., Al-Shabeeb, A., Nikolic, A., Catalano, A., Iland, H., Horvath, N., Ho, P.J., Harrison, S., Fleming, S., et al. (2012). Response of myeloma to the proteasome inhibitor bortezomib is correlated with the unfolded protein response regulator XBP-1. Haematologica 97, 64–72.

Liu, N., Zuo, C., Wang, X., Chen, T., Yang, D., Wang, J., and Zhu, H. (2014). miR-942 decreases TRAIL-induced apoptosis through ISG12a downregulation and is regulated by AKT. Oncotarget 5, 4959–4971.

Llamas, E., Alirzayeva, H., Loureiro, R., and Vilchez, D. (2020). The intrinsic proteostasis network of stem cells. Curr Opin Cell Biol 67, 46–55.

Lu, T.G., Bankhead, A., Ljungman, M., and Neamati, N. (2019). Multi-omics profiling reveals key signaling pathways in ovarian cancer controlled by STAT3. Theranostics 9, 5478–5496.

Masuoka, H.C., and Townes, T.M. (2002). Targeted disruption of the activating transcription factor 4 gene results in severe fetal anemia in mice. Blood 99, 736–745.

Maurel, M., Chevet, E., Tavernier, J., and Gerlo, S. (2014). Getting RIDD of RNA: IRE1 in cell fate regulation. Trends Biochem Sci 39, 245–254.

Meiners, S., Heyken, D., Weller, A., Ludwig, A., Stangl, K., Kloetzel, P.M., and Kruger, E. (2003). Inhibition of proteasome activity induces concerted expression of proteasome genes and de novo formation of Mammalian proteasomes. J Biol Chem 278, 21517–21525.

Miyahira, A.K., Shahangian, A., Hwang, S.M., Sun, R., and Cheng, G.H. (2009). TANK-Binding Kinase-1 Plays an Important Role during In Vitro and In Vivo Type I IFN Responses to DNA Virus Infections. J Immunol 182, 2248–2257.

Muller-Newen, G., Stope, M.B., Kraus, T., and Ziegler, P. (2017). Development of platelets during steady state and inflammation. J Leukocyte Biol 101, 1109–1117.

Northrop, A., Vangala, J.R., Feygin, A., and Radhakrishnan, S.K. (2020). Disabling the Protease DDI2 Attenuates the Transcriptional Activity of NRF1 and Potentiates Proteasome Inhibitor Cytotoxicity. International Journal of Molecular Sciences 21.

Novoa, I., Zhang, Y.H., Zeng, H.Q., Jungreis, R., Harding, H.P., and Ron, D. (2003). Stress-induced gene expression requires programmed recovery from translational repression (vol 22, pg 1180, 2003). Embo Journal 22, 2307–2307.

Nowak, K., Taubert, R.M., Haberecht, S., Venz, S., and Kruger, E. (2018). Inhibition of calpain-1 stabilizes TCF11/Nrf1 but does not affect its activation in response to proteasome inhibition. Biosci Rep 38.

Pakos-Zebrucka, K., Koryga, I., Mnich, K., Ljujic, M., Samali, A., and Gorman, A.M. (2016). The integrated stress response. Embo Rep 17, 1374–1395.

Perry, A.K., Chow, E.K., Goodnough, J.B., Yeh, W.C., and Cheng, G. (2004). Differential requirement for TANK-binding kinase-1 in type I interferon responses to toll-like receptor activation and viral infection. Journal of Experimental Medicine 199, 1651–1658.

Poli, M.C., Ebstein, F., Nicholas, S.K., de Guzman, M.M., Forbes, L.R., Chinn, I.K., Mace, E.M., Vogel, T.P., Carisey, A.F., Benavides, F., et al. (2018). Heterozygous Truncating Variants in POMP Escape Nonsense-Mediated Decay and Cause a Unique Immune Dysregulatory Syndrome. Am J Hum Genet 102, 11261142.

Radhakrishnan, S.K., Lee, C.S., Young, P., Beskow, A., Chan, J.Y., and Deshaies, R.J. (2010). Transcription Factor Nrf1 Mediates the Proteasome Recovery Pathway after Proteasome Inhibition in Mammalian Cells. Molecular Cell 38, 17–28.

Ramirez, J., Lectez, B., Osinalde, N., Siva, M., Elu, N., Aloria, K., Prochazkova, M., Perez, C., Martinez-Hernandez, J., Barrio, R., et al. (2018). Quantitative proteomics reveals neuronal ubiquitination of Rngo/Ddi1 and several proteasomal subunits by Ube3a, accounting for the complexity of Angelman syndrome. Hum Mol Genet 27, 1955–1971.

Rawlings, J.S., Rosler, K.M., and Harrison, D.A. (2004). The JAK/STAT signaling pathway. J Cell Sci 117, 1281–1283.

Rice, G.I., Melki, I., Fremond, M.L., Briggs, T.A., Rodero, M.P., Kitabayashi, N., Oojageer, A., Bader-Meunier, B., Belot, A., Bodemer, C., et al. (2017). Assessment of Type I Interferon Signaling in Pediatric Inflammatory Disease. J Clin Immunol 37, 123–132.

Rosebeck, S., and Leaman, D.W. (2008). Mitochondrial localization and pro-apoptotic effects of the interferon-inducible protein ISG12a. Apoptosis 13, 562–572.

Sakao, Y., Kawai, T., Takeuchi, O., Copeland, N.G., Gilbert, D.J., Jenkins, N.A., Takeda, K., and Akira, S. (2000). Mouse proteasomal ATPases Psmc3 and Psmc4: genomic organization and gene targeting. Genomics 67, 1–7.

Schneider, K., Nelson, G.M., Watson, J.L., Morf, J., Dalglish, M., Luh, L.M., Weber, A., and Bertolotti, A. (2020). Protein Stability Buffers the Cost of Translation Attenuation following eIF2alpha Phosphorylation. Cell Rep 32, 108154.

Seifert, U., Bialy, L.P., Ebstein, F., Bech-Otschir, D., Voigt, A., Schroter, F., Prozorovski, T., Lange, N., Steffen, J., Rieger, M., et al. (2010). Immunoproteasomes preserve protein homeostasis upon interferon-induced oxidative stress. Cell 142, 613–624.

Serbyn, N., Noireterre, A., Bagdiul, I., Plank, M., Michel, A.H., Loewith, R., Kornmann, B., and Stutz, F. (2020). The Aspartic Protease Ddi1 Contributes to DNA-Protein Crosslink Repair in Yeast. Molecular Cell 77, 1066–+.

Siva, M., Svoboda, M., Veverka, V., Trempe, J.F., Hofmann, K., Kozisek, M., Hexnerova, R., Sedlak, F., Belza, J., Brynda, J., et al. (2016). Human DNA-Damage-Inducible 2 Protein Is Structurally and Functionally Distinct from Its Yeast Ortholog. Sci Rep 6, 30443.

So, J.S., Hur, K.Y., Tarrio, M., Ruda, V., Frank-Kamenetsky, M., Fitzgerald, K., Koteliansky, V., Lichtman, A.H., Iwawaki, T., Glimcher, L.H., et al. (2012). Silencing of lipid metabolism genes through IRE1alpha-mediated mRNA decay lowers plasma lipids in mice. Cell Metab 16, 487–499.

Sotzny, F., Schormann, E., Kuhlewindt, I., Koch, A., Brehm, A., Goldbach-Mansky, R., Gilling, K.E., and Kruger, E. (2016). TCF11/Nrf1-Mediated Induction of Proteasome Expression Prevents Cytotoxicity by Rotenone. Antioxid Redox Signal 25, 870–885.

Steffen, J., Seeger, M., Koch, A., and Kruger, E. (2010). Proteasomal degradation is transcriptionally controlled by TCF11 via an ERAD-dependent feedback loop. Mol Cell 40, 147–158.

Studencka-Turski, M., Cetin, G., Junker, H., Ebstein, F., and Kruger, E. (2019). Molecular Insight Into the IRE1alpha-Mediated Type I Interferon Response Induced by Proteasome Impairment in Myeloid Cells of the Brain. Front Immunol 10, 2900.

Svoboda, M., Konvalinka, J., Trempe, J.F., and Saskovaa, K.G. (2019). The yeast proteases Ddi1 and Wss1 are both involved in the DNA replication stress response. DNA Repair 80, 45–51.

Tanaka, K., and Chiba, T. (1998). The proteasome: a protein-destroying machine. Genes Cells 3, 499–510.

Todoric, J., and Karin, M. (2019). The Fire within: Cell-Autonomous Mechanisms in Inflammation-Driven Cancer. Cancer Cell 35, 714–720.

Tomlin, F.M., Gerling-Driessen, U.I.M., Liu, Y.C., Flynn, R.A., Vangala, J.R., Lentz, C.S., Clauder-Muenster, S., Jakob, P., Mueller, W.F., Ordonez-Rueda, D., et al. (2017). Inhibition of NGLY1 Inactivates the Transcription Factor Nrf1 and Potentiates Proteasome Inhibitor Cytotoxicity. ACS Cent Sci 3, 1143–1155.

Trempe, J.F., Saskova, K.G., Siva, M., Ratcliffe, C.D., Veverka, V., Hoegl, A., Menade, M., Feng, X., Shenker, S., Svoboda, M., et al. (2016). Structural studies of the yeast DNA damage-inducible protein Ddi1 reveal domain architecture of this eukaryotic protein family. Sci Rep 6, 33671.

Tsai, M.H., Pai, L.M., and Lee, C.K. (2019). Fine-Tuning of Type I Interferon Response by STAT3. Frontiers in Immunology 10.

Tsuzuki, S., Tachibana, M., Hemmi, M., Yamaguchi, T., Shoji, M., Sakurai, F., Kobiyama, K., Kawabata, K., Ishii, K.J., Akira, S., et al. (2016). TANK-binding kinase 1-dependent or -independent signaling elicits the cell-type-specific innate immune responses induced by the adenovirus vector. Int Immunol 28, 105–115.

Vangala, J.R., Sotzny, F., Kruger, E., Deshaies, R.J., and Radhakrishnan, S.K. (2016). Nrf1 can be processed and activated in a proteasome-independent manner. Curr Biol 26, R834–R835.

Vomund, S., Schafer, A., Parnham, M.J., Brune, B., and von Knethen, A. (2017). Nrf2, the Master Regulator of Anti-Oxidative Responses. International Journal of Molecular Sciences 18.

Welk, V., Coux, O., Kleene, V., Abeza, C., Trumbach, D., Eickelberg, O., and Meiners, S. (2016). Inhibition of Proteasome Activity Induces Formation of Alternative Proteasome Complexes. Journal of Biological Chemistry 291, 13147–13159.

Whitby, F.G., Masters, E.I., Kramer, L., Knowlton, J.R., Yao, Y., Wang, C.C., and Hill, C.P. (2000). Structural basis for the activation of 20S proteasomes by 11S regulators. Nature 408, 115–120.

Widenmaier, S.B., Snyder, N.A., Nguyen, T.B., Arduini, A., Lee, G.Y., Arruda, A.P., Saksi, J., Bartelt, A., and Hotamisligil, G.S. (2017). NRF1 Is an ER Membrane Sensor that Is Central to Cholesterol Homeostasis. Cell 171, 1094–+.

Wilkinson, D.G., and Nieto, M.A. (1993). Detection of Messenger-Rna by in-Situ Hybridization to Tissue-Sections and Whole Mounts. Method Enzymol 225, 361–373.

Xu, H., You, M., Shi, H., and Hou, Y. (2015). Ubiquitin-mediated NFkappaB degradation pathway. Cell Mol Immunol 12, 653–655.

Yang, K., Huang, R., Fujihira, H., Suzuki, T., and Yan, N. (2018). N-glycanase NGLY1 regulates mitochondrial homeostasis and inflammation through NRF1. J Exp Med 215, 2600–2616.

Yip, M.C.J., Bodnar, N.O., and Rapoport, T.A. (2020). Ddi1 is a ubiquitin-dependent protease. Proc Natl Acad Sci U S A.

Yokoi, M., and Hanaoka, F. (2017). Two mammalian homologs of yeast Rad23, HR23A and HR23B, as multifunctional proteins. Gene 597, 1–9.

Yousaf, A., Wu, Y., Khan, R., Shah, W., Khan, I., Shi, Q., and Jiang, X. (2020). Normal spermatogenesis and fertility in Ddi1 (DNA damage inducible 1) mutant mice. Reprod Biol.

Yu, Q., Katlinskaya, Y.V., Carbone, C.J., Zhao, B., Katlinski, K.V., Zheng, H., Guha, M., Li, N., Chen, Q., Yang, T., et al. (2015). DNA-damage-induced type I interferon promotes senescence and inhibits stem cell function. Cell Rep 11, 785–797.

Yu, Y., and Hayward, G.S. (2010). The ubiquitin E3 ligase RAUL negatively regulates type i interferon through ubiquitination of the transcription factors IRF7 and IRF3. Immunity 33, 863–877.

Zhang, X.D., Baladandayuthapani, V., Lin, H., Mulligan, G., Li, B., Esseltine, D.L.W., Qi, L., Xu, J.L., Hunziker, W., Barlogie, B., et al. (2016). Tight Junction Protein 1 Modulates Proteasome Capacity and Proteasome Inhibitor Sensitivity in Multiple Myeloma via EGFR/JAK1/STAT3 Signaling. Cancer Cell 29, 639–652.

Zhao, J., Garcia, G.A., and Goldberg, A.L. (2016). Control of proteasomal proteolysis by mTOR. Nature 529, E1–2.

Zhao, J., and Goldberg, A.L. (2016). Coordinate regulation of autophagy and the ubiquitin proteasome system by MTOR. Autophagy 12, 1967–1970.

