## Supplemental Information for "DDI2 protease activity controls embryonic development and inflammation via TCF11/NRF1"

Supplementary Figures 1-7

Supplementary Table 1-5 TALENs, oligonucleotides and probes used in this study

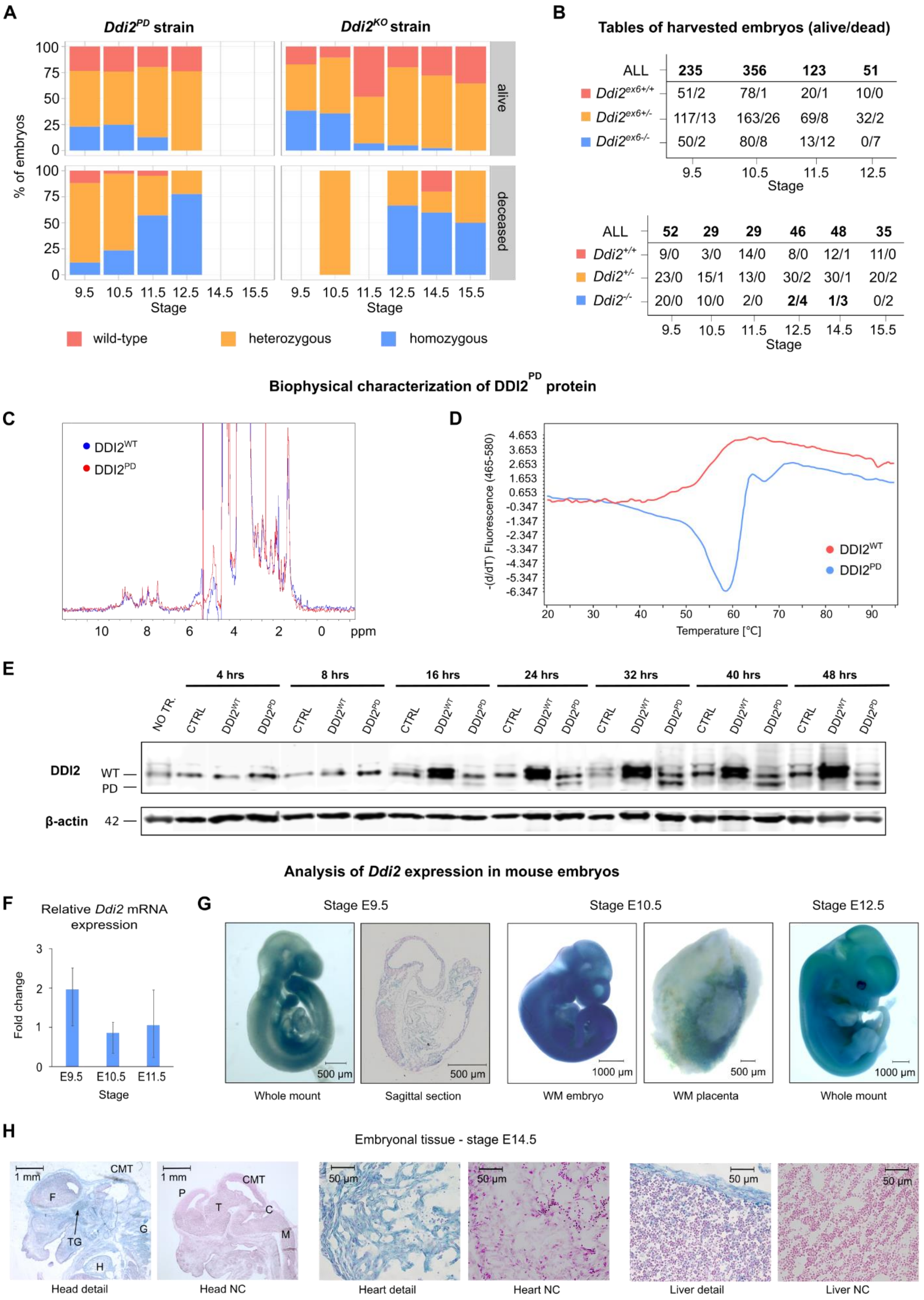
A
Ddi2PD strain
Ddi2KO strain
% of embryos
alive
deceased
100
75
50
25
0
9.5 10.5 11.5 12.5 14.5 15.5
Stage
wild-type
heterozygous
homozygous
B
Tables of harvested embryos (alive/dead)
ALL 235 356 123 51
Ddi2ex6+/+ 51/2 78/1 20/1 10/0
Ddi2ex6+/- 117/13 163/26 69/8 32/2
Ddi2ex6-/- 50/2 80/8 13/12 0/7
9.5 10.5 11.5 12.5
Stage
ALL 52 29 29 46 48 35
Ddi2+/+ 9/0 3/0 14/0 8/0 12/1 11/0
Ddi2+/- 23/0 15/1 13/0 30/2 30/1 20/2
Ddi2-/- 20/0 10/0 2/0 2/4 1/3 0/2
9.5 10.5 11.5 12.5 14.5 15.5
Stage
Biophysical characterization of DDI2PD protein
C
DDI2WT
DDI2PD
10 8 6 4 2 0 ppm
D
-(d/dT) Fluorescence (465-580)
4.653
3.653
2.653
1.653
0.653
-0.347
-1.347
-2.347
-3.347
-4.347
-5.347
-6.347
DDI2WT
DDI2PD
20 30 40 50 60 70 80 90
Temperature [°C]
E
4 hrs
8 hrs
16 hrs
24 hrs
32 hrs
40 hrs
48 hrs
NO TR.
CTRL
DDI2WT
DDI2PD
DDI2
WT
PD
β-actin
42
Analysis of Ddi2 expression in mouse embryos
F
Relative Ddi2 mRNA expression
Fold change
3
2
1
0
E9.5 E10.5 E11.5
Stage
G
Stage E9.5
Stage E10.5
Stage E12.5
500 µm
500 µm
1000 µm
500 µm
1000 µm
Whole mount
Sagittal section
WM embryo
WM placenta
Whole mount
H
Embryonal tissue - stage E14.5
1 mm
CMT
F
TG
G
H
P
T
C
M
50 µm
Head detail
Head NC
Heart detail
Heart NC
Liver detail
Liver NC

**Figure S1: Mapping of *Ddi2* expression in *Ddi2$^{PD}$ and Ddi2$^{KO}$* mouse strains and characterization of protein folding and stability of DDI2$^{PD}$ expressed in *Ddi2$^{ex6+/-}$* and *Ddi2$^{ex6-/-}$* mice**

**(A)** Lethality screening of *Ddi2$^{PD}$ and Ddi2$^{KO}$* mouse strains. Barplots show representative percentages of individual genotypes in two groups (living and dead embryos) for each developmental stage harvested. Data from lethality screening was analyzed using the program R, version 3.6.2 (2019-12-12).

**(B)** Tables showing total counts of harvested embryos of *Ddi2$^{PD}$ and Ddi2$^{KO}$* mouse strains in the lethality screen. Tables include resorbed fetuses (N/A) at each stage of harvest.

**(C)** Overlay of 1D NMR spectra of DDI2$^{WT}$ (blue) and DDI2$^{PD}$ protein (red) shows acquired secondary structures for both versions of the protein. Coding sequences were acquired from *Ddi2$^{ex6+/+}$* and *Ddi2$^{ex6-/-}$* embryonal mRNA.

**(D)** Recombinantly expressed DDI2$^{PD}$ protein appears to form aggregates based on melting curves measured by DSF - DDI2$^{WT}$ (red) and DDI2$^{PD}$ (blue).

**(E)** The level of the mouse DDI2$^{PD}$ protein signal is much lower than that of DDI2$^{WT}$ at identical time points after transfection of HEK293-TetOff-A2 cells. Cells transiently transfected with pTreTight vector encoding DDI2$^{PD}$, DDI2$^{WT}$ or an empty vector (CTRL) were harvested after 4, 8, 16, 24, 32, 40 and 48 hours. β-actin detection was used as a loading control.

**(F)** *Ddi2* expression increases at E9.5 compared to E10.5 and E11.5. Expression of *Ddi2* in *C57BL/6NCrl* embryos at stages E9.5, E10.5 and E11.5 (n=6) was analyzed relatively to expression of the *H2afz* housekeeping gene by qRT-PCR. Error bars denote min to max values. Data were analyzed in MS Excel (2019) and the graph was created using GraphPad Prism 6 software.

**(G)** Mapping of *Ddi2* expression using a *LacZ* reporter gene. β-galactosidase activity was measured in *Ddi2$^{+/-}$* embryos at stages E9.5, E10.5 and E12.5. While expression of *LacZ* (*Ddi2*) at E9.5 is present in rapidly developing tissues, such as limb buds and orofacial processes, expression at later stages is more ubiquitous in the embryonic body. Expression in E10.5 placenta most likely occurs in the chorionic plate.

**(H)** *LacZ* reporter gene expression mapping on sagittal sections of E14.5 stage *Ddi2$^{+/-}$* embryos. Details of the head, heart and liver are shown. NC is a negative control for β-Galactosidase activity staining of *Ddi2$^{+/+}$* embryo sections. Localization of *Ddi2* expression appears to be specific for ectodermal and mesodermal tissue, such as skin, brain (forebrain and trigeminal ganglion), cranium, and smooth muscle of heart tissue. Liver cells that likely correspond to Kupffer cells also exhibit *Ddi2* expression at stage E14.5. Abbreviations: C – cerebellum, F – forebrain CMT – colliculus midbrain tectum, G - ganglions, H – heart, P – pallium, M – medulla, T - thalamus, TG – trigeminal ganglion.

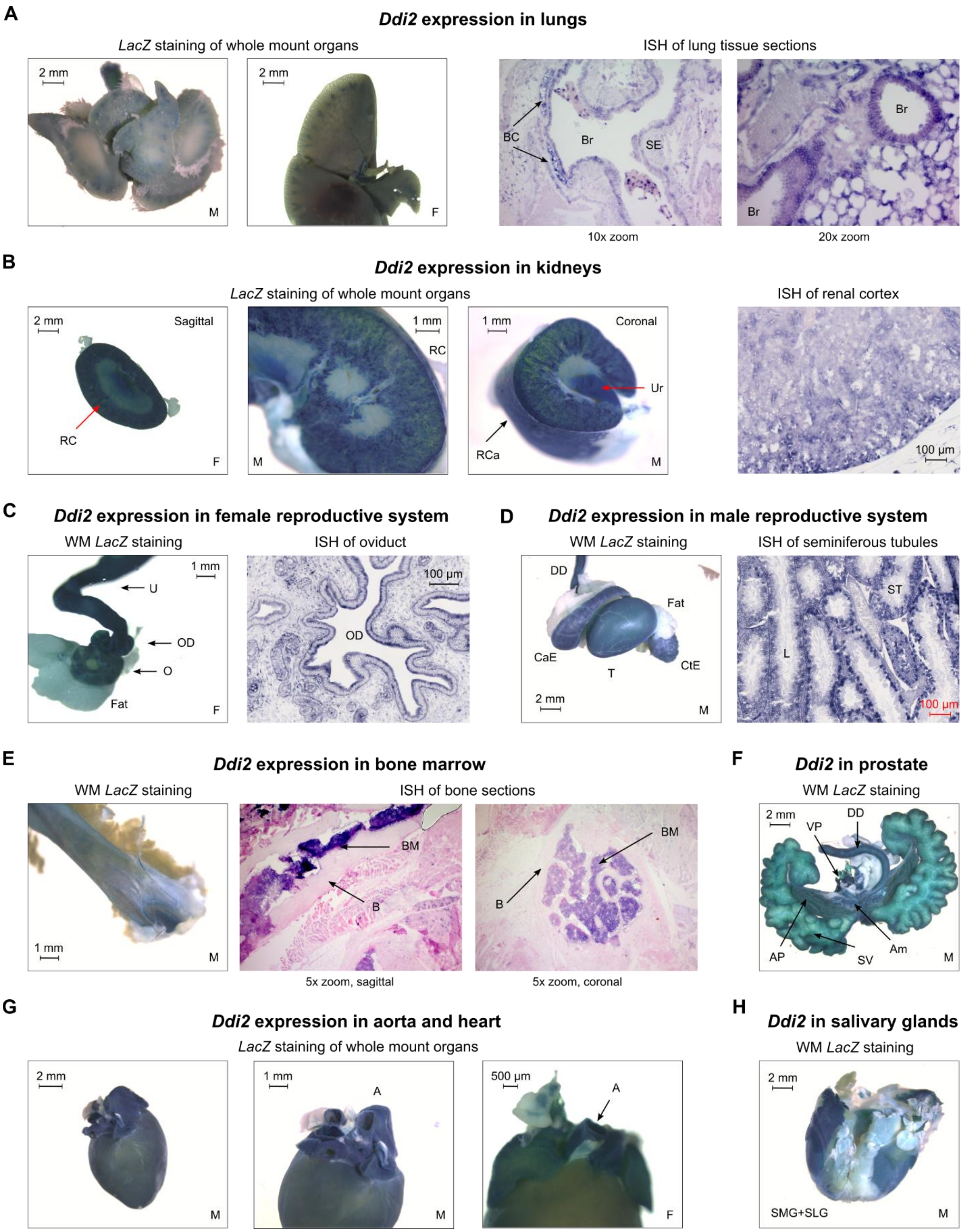

A
Ddi2 expression in lungs
LacZ staining of whole mount organs
2 mm
M
2 mm
F
ISH of lung tissue sections
BC
Br
SE
Br
Br
10x zoom
20x zoom
B
Ddi2 expression in kidneys
LacZ staining of whole mount organs
2 mm
Sagittal
RC
F
1 mm
RC
M
1 mm
Coronal
Ur
RCa
M
ISH of renal cortex
100 µm
C
Ddi2 expression in female reproductive system
WM LacZ staining
1 mm
U
OD
O
Fat
F
ISH of oviduct
100 µm
OD
D
Ddi2 expression in male reproductive system
WM LacZ staining
DD
Fat
CaE
T
CtE
2 mm
M
ISH of seminiferous tubules
ST
L
100 µm
E
Ddi2 expression in bone marrow
WM LacZ staining
1 mm
M
ISH of bone sections
BM
B
BM
B
5x zoom, sagittal
5x zoom, coronal
F
Ddi2 in prostate
WM LacZ staining
2 mm
DD
VP
AP
SV
Am
M
G
Ddi2 expression in aorta and heart
LacZ staining of whole mount organs
2 mm
M
1 mm
A
M
500 µm
A
F
H
Ddi2 in salivary glands
WM LacZ staining
2 mm
SMG+SLG
M

**Figure S2: *Ddi2* expression in mesodermal and endodermal tissue**

*Ddi2* expression was mapped using a β-galactosidase activity assay on whole-mount tissue of *Ddi2*$^{+/-}$ adult mice or using ISH on *C57BL/6NCrl* mouse tissue sections. The sex of the animal is marked with M or F in one of the bottom corners of each image.

**(A)** *Ddi2* is expressed in lungs in bronchiolar endothelial cells, capillaries between pneumocytes and in bronchial cartilage.

**(B)** *Ddi2* is expressed in epithelial cells of the renal cortex as can be observed in both sagittal and coronal sections of kidneys. Expression also occurs in ureter.

**(C)** *Ddi2* is expressed in the female reproductive system in the oviduct, where it is specifically expressed in epithelial layer.

**(D)** Testes, ductus deferens and epididymis stain positive for expression of *Ddi2* in the male reproductive system. In testes, expression occurs in the epithelial layer of seminiferous tubules.

**(E)** As shown by WM staining of a femur cut in half and on sagittal and coronal sections of femur, *Ddi2* is expressed in bone marrow.

**(F)** *Ddi2* is expressed in ampulla, prostate and ductus deferens.

**(G)** *Ddi2* expression can be observed in arterial endothelial cells.

**(H)** Salivary glands are shown as examples of gland organs of endodermal derivation in which *Ddi2* is expressed.

Legend: A – aorta, Am – ampulla, AP - anterior prostate, B – bone, BC – bronchial cartilage, BM – bone marrow, Br – bronchiole, CaE – cauda epididymis, CtE – caput epididymis, DD – ductus deferens, F – female, L – lumen, M – male, O – ovary, OD – oviduct, RC – renal cortex, RCa – renal capsule, SE – squamous epithelium, SMG+SLG – submandibular and sublingual glands, ST – seminiferous tubules, SV – seminal vesicles, T – testes, U – uterus, Ur – ureter, VP – ventral prostate.

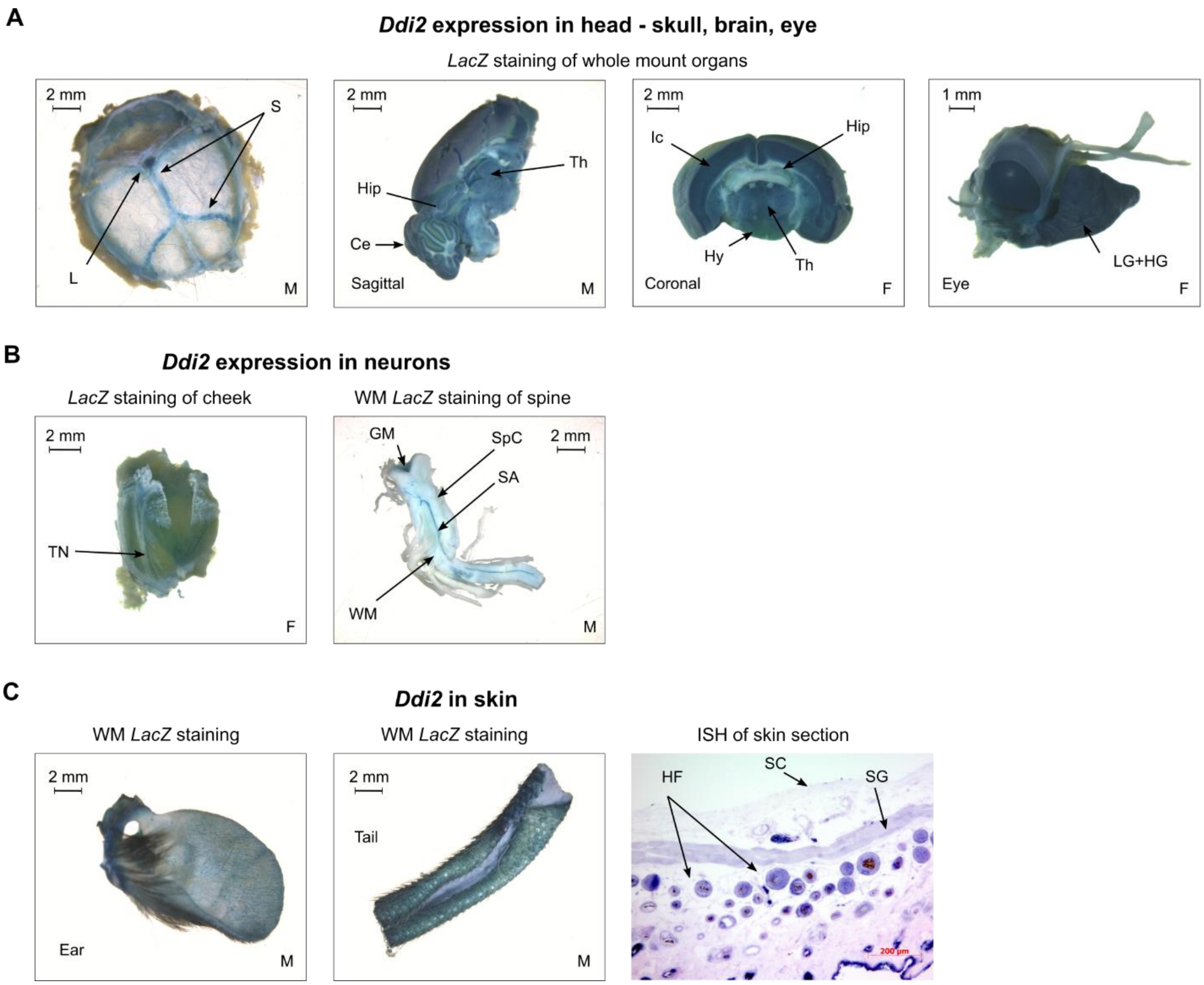


**Figure S3: *Ddi2* expression in ectodermal tissue**

*Ddi2* expression was mapped using a β-galactosidase activity assay on whole-mount tissue of *Ddi2*$^{+/-}$ adult mice or using ISH on *C57BL/6NCrl* mouse tissue sections. The sex of the animal is marked with M or F in one of the bottom corners of each image.

**(A)** Head: *Ddi2* is expressed in sutures of skull, in grey matter in brain (cerebellum, isocortex, thalamus) as seen on both sagittal and coronal sections, and in the eye and intraorbital glands (lacrimal and Harderian).

**(B)** *Ddi2* is expressed in peripheral neurons such as trigeminal ganglion and the grey matter of the spinal cord.

**(C)** In skin, *Ddi2* is expressed in the epidermis (stratum granulosum) and hair follicles.

Legend: Ce – cerebellum, GM – grey matter, HF – hair follicles, Hip – hippocampus, Hy – hypothalamus, Ic – isocortex, L – lambda, LG+HG – lacrimal and Harderian glands, S – sutures, SA – spinal artery, SC –

stratum corneum, SG – stratum granulosum, SpC – spinal cord, Th – thalamus, TN – trigeminal nerve, WM – white matter.

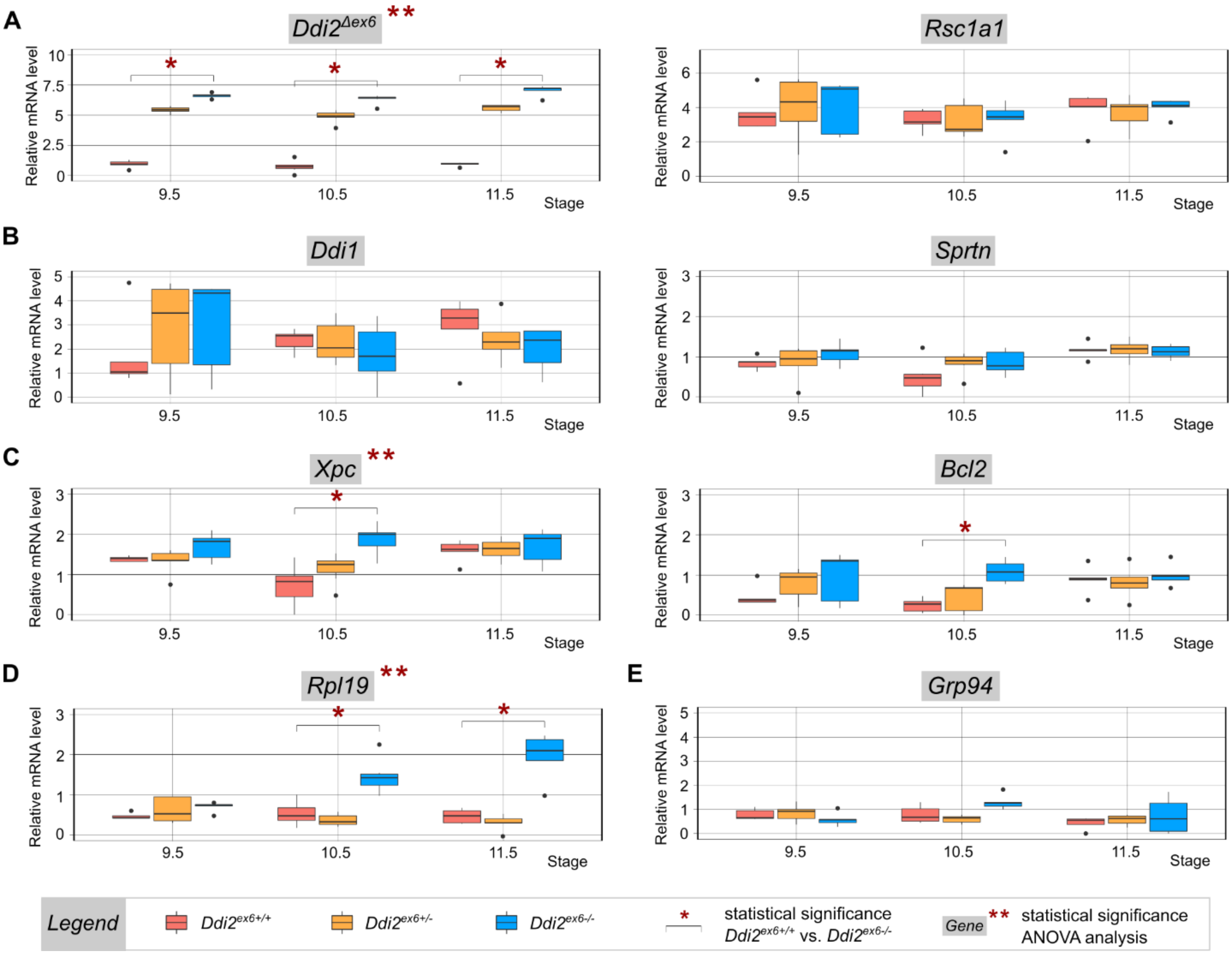


**Figure S4: qRT-PCR screen in *Ddi2*$^{PD}$ embryos**

**(A)** Control expression analysis of the *Ddi2*$^{\Delta ex6}$ and *Rsc1a1* genes. As the vertebrate-specific gene *Rsc1a1* was inserted upstream of the UBA domain of the *Ddi2* gene locus during evolution (Siva et al., 2016), analysis of its expression was included in our qRT-PCR screen as a control to determine whether TALEN-mediated excision of exon 6 affected *Rsc1a1* expression.

**(B)** Expression of neither *Ddi1* nor *Sprtn* is altered during mid-gestation embryonic development upon DDI2 depletion in mice. *Ddi1* is a single-intron homolog of *Ddi2*. *Sprtn* is the mouse ortholog of a yeast metalloprotease involved, along with yeast Ddi1p (Svoboda et al., 2019), in the DNA replication stress response.

**(C)** Both the DNA damage marker *Xpc* and the apoptosis marker *Bcl2* exhibit a statistically significant increase in expression at stage E10.5 upon *Ddi2* loss of function.

**(D)** The Ribosomal protein *Rpl19* shows an increase in expression in embryos lacking functional *Ddi2*.

**(E)** *Grp94* in the UPS exhibits no response to depletion of *Ddi2* function in mouse embryos.

Legend: *Ddi2*$^{ex6+/+}$ - red, *Ddi2*$^{ex6+/-}$ - yellow, *Ddi2*$^{ex6-/-}$ - blue; E9.5 (n=5), E10.5 (n=7), E11.5 (n=5). Relative expression of genes was normalized to the *Tbp* and *H2afz* housekeeping genes; outliers were omitted based on the Grubbs' test. Statistical significance was calculated either for each gene throughout all three developmental stages with application of ANOVA statistical analysis (**) or using a linear mixed-effects model (LMM) to compare gene expression of wild-type and homozygous embryos at each developmental stage (*). Both analyses were subjected to Bonferroni correction. Data and boxplots were processed using the program R, version 3.6.2 (2019-12-12). All primers used for qRT-PCR screen are listed in Table S4.

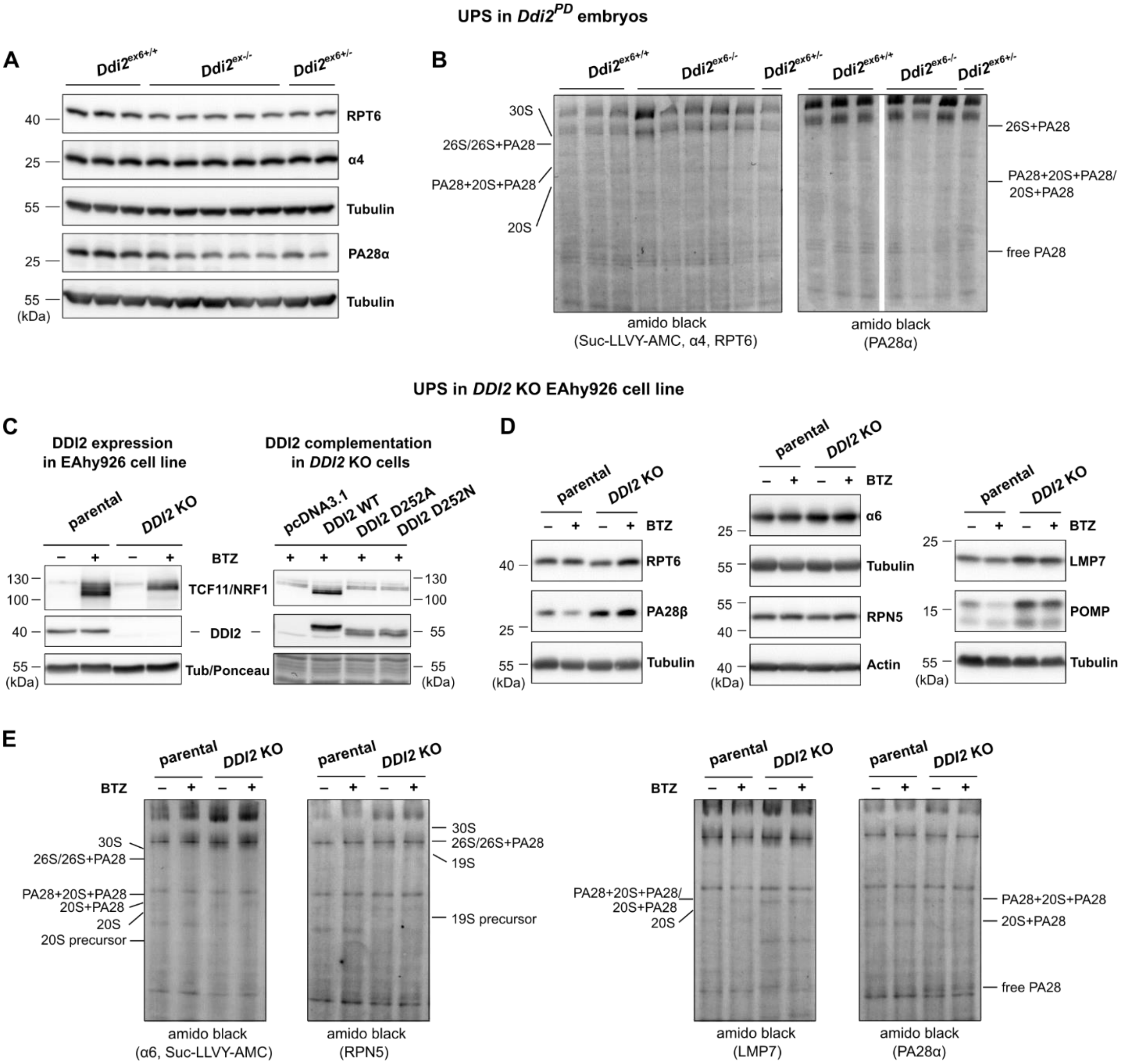


**Figure S5: Proteasomal subunit composition changes upon DDI2 depletion in stage E10.5 *Ddi2$^{PD}$* embryos and human endothelial knock-out cells**

**(A)** Expression of several proteasomal subunits (RPT6, α4 and PA28α) in *Ddi2$^{PD}$* embryos was analyzed on immunoblots using tissue lysates of *Ddi2$^{ex6+/+}$* (n=6), *Ddi2$^{ex6–/–}$* (n=10) and *Ddi2$^{ex6+/–}$* (n=3) embryos. Tubulin was used as a loading control. The total amount of protein loaded was 20ug/lane (RPT6 and α4) and 40ug protein/lane (PA28α).

**(B)** Amido black loading controls of native PAGE analysis (Figure 3) of embryo lysates (15 µg protein/lane).

**(C)** Left: Western blot analysis of TCF11/NRF1 and DDI2 protein levels in EAhy926 parental and *DDI2* KO cells treated with 50 nM BTZ for 8 hours compared to non-treated controls. Tubulin was used as a loading control.

Right: Western blot analysis of whole EAhy926 *DDI2* KO cell lysates transfected with either empty pcDNA3.1V5-His TOPO vector or with V5-tagged constructs of the wild type (WT) DDI2 protein, the D252A active site mutant, or the D252N active site mutant (n=2). Cells were treated with 500 nM BTZ for 3 hours. Immunoblots were stained for TCF11/NRF1 and DDI2. Ponceau staining served as a loading control.

**(D)** Western blot analysis of protein expression in whole cell extracts of EAhy926 parental and *DDI2* KO cells treated with 50 nM BTZ for 8 hours (n=4). Non-treated cells served as control. Membranes were probed for the proteasomal subunits RPT6, PA28β, α6, RPN5, LMP7 and POMP. Tubulin and actin served as loading controls. The total amount of protein loaded was 15 µg/lane for RPT6 and PA28β and 20 µg/lane for all other subunits.

**(E)** Amido black loading controls of native PAGE analysis of EAhy926 cell lysates (20 µg protein/lane). For more information see Figure 3.

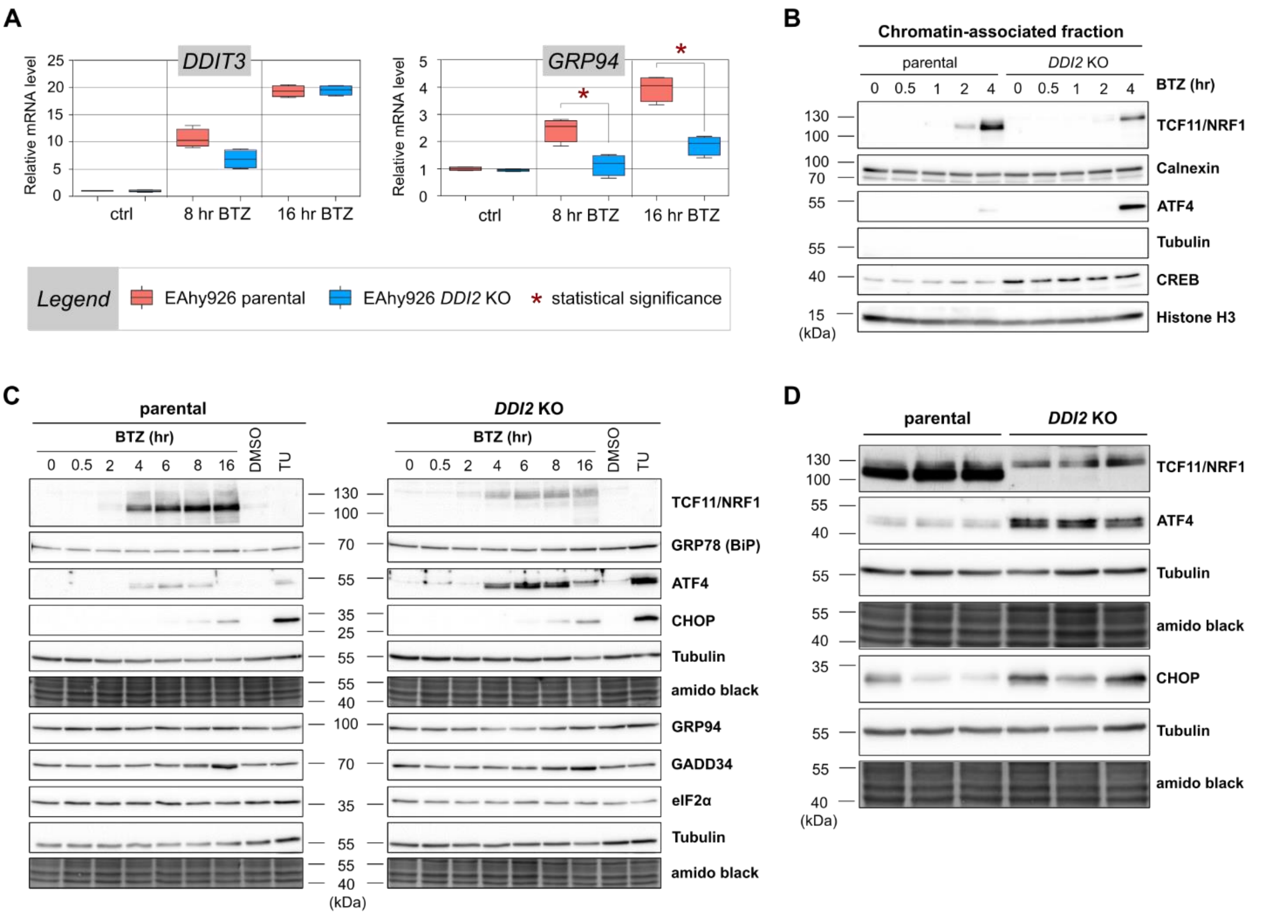


**Figure S6: Depletion of DDI2 in human endothelial EAhy926 cells results in upregulation of several UPR pathway markers and changes in response to proteasomal inhibition**

**(A) q**RT-PCR analysis of *GRP94* and *DDIT3* (CHOP) mRNA levels in EAhy926 parental (red) and *DDI2* KO (blue) cells treated with 50 nM BTZ for 8 or 16 hours. Non-treated cells were used as a negative control; mRNA amounts were normalized to *RPLP0* and Grubbs' outlier test was applied. The statistical significance of the difference between parental and *DDI2* KO cells was calculated individually for each BTZ treatment (n=4) and non-treatment control (n=4) time point using the unpaired two-sided t-test with applied Holm-Šidák correction (with *α=5%) using GraphPad Prism 6 software.

**(B)** Cellular fractionation of chromatin-associated proteins of EAhy926 parental and *DDI2* KO cells treated with 50 nM BTZ for up to 4 hours (n=4). Immunoblots were probed for TCF11/NRF1 and ATF4. Calnexin, tubulin, CREB and histone H3 served as marker proteins. For non-nuclear and nuclear fractions see Figure 5.

**(C)** Western blot analysis of TCF11/NRF1 and several downstream targets of the PERK branch of the UPR signaling pathway in whole cell extracts of EAhy926 parental and *DDI2* KO cells (n=3). Cells were treated with 50 nM BTZ for up to 16 hours and compared a to non-treated control. Treatment with 4 µg/mL of tunicamycin (TU), an inhibitor of N-glycosidic linkages in glycoprotein synthesis, for 6 hours served as a positive control for ER stress. Cells treated with 0.04% DMSO were included as a solvent control for TU treatment. Immunoblots were probed for TCF11/NRF1, GRP78 (BiP), ATF4 and CHOP (40 µg protein/lane) as well as GRP94, GADD34 and eIF2α (20 µg protein/lane). Both amino black and tubulin were used as loading controls.

**(D)** Western blot analysis of TCF11/NRF1, ATF4 and CHOP expression in whole cell extracts of EAhy926 parental and *DDI2* KO cells treated with 50 nM BTZ for 8 hours (TCF11/NRF1 and ATF4 detection) or 16 hours (CHOP detection). Note that 3 replicates are shown. This immunoblot was used for normalization of quantified protein expression levels. For further information see Figure 5.

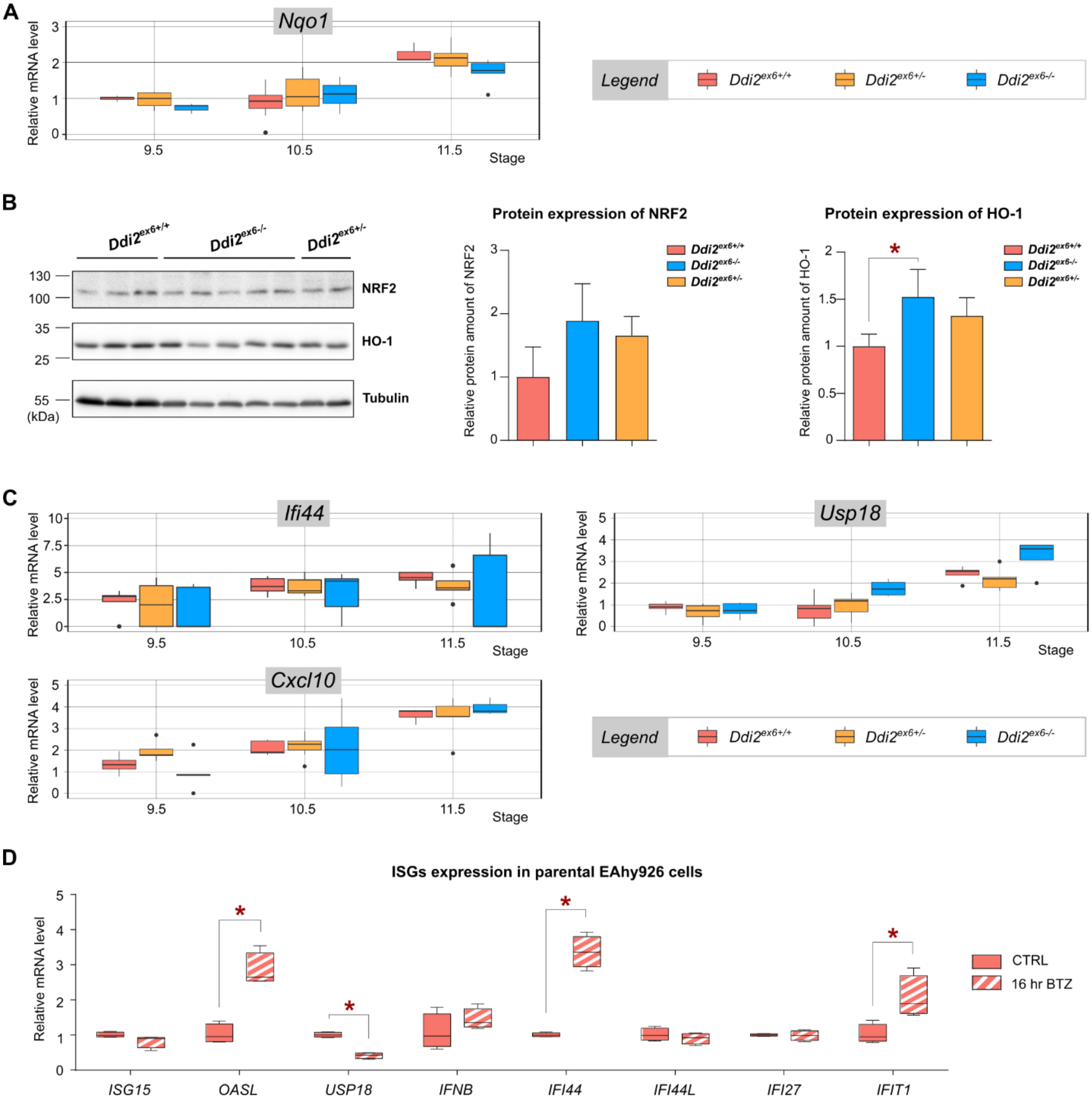


**Figure S7: Changes in expression of genes connected to oxidative stress response and interferon signaling upon DDI2 depletion in *Ddi2$^{PD}$* embryos and EAhy926 cells.**

**(A)** Expression of *Nqo1*, a gene involved in oxidative stress response, in *Ddi2$^{PD}$* strain embryos in qRT-PCR screening. Legend: *Ddi2$^{ex6+/+}$* - red, *Ddi2$^{ex6+/-}$* - yellow, *Ddi2$^{ex6-/-}$* - blue; E9.5 (n=5), E10.5 (n=7), E11.5 (n=5). Relative expression of *Nqo1* was normalized to *Tbp* and *H2afz* housekeeping genes; outliers were omitted based on the Grubbs' test. Statistical significance was calculated either for each gene throughout all three developmental stages with application of ANOVA statistical analysis or using a linear mixed-effects model (LMM) to compare gene expression in wild-type and homozygous embryos at each developmental stage.

Both analyses were subjected to Bonferroni correction. Data and boxplots were processed using the program R, version 3.6.2 (2019-12-12). All primers used for qRT-PCR screen are listed in Table S4.

**(B)** Western blot analysis of NRF2 and HO-1 expression in tissue lysates of *Ddi2*$^{ex6+/+}$ (n=3, red), *Ddi2*$^{ex6–/–}$ (n=5, blue) and *Ddi2*$^{ex6+/–}$ (n=2, yellow). Protein expression was quantified using ImageJ software and normalized using a tubulin control. Statistical significance was determined using GraphPad Prism 6 software (mean ± SD, *p value < 0.05, unpaired two-sided t-test).

**(C)** Expression of several interferon genes in *Ddi2*$^{em1}$*/Ph* strain embryos in qRT-PCR screening. *Ddi2*$^{ex6+/+}$ - red, *Ddi2*$^{ex6+/-}$ - yellow, *Ddi2*$^{ex6-/-}$ - blue; E9.5 (n=5), E10.5 (n=7), E11.5 (n=5). Computation of relative gene expression, statistical analysis and data processing were performed as described in (A).

**(D)** qRT-PCR analysis of ISG mRNA levels in EAhy926 parental cells treated with 50 nM BTZ for 16 hours (red-white pattern) and in non-treated control cells (red). Messenger RNA amounts were normalized to *RPLP0* and the Grubbs' outlier test was applied. The statistical significance of the difference between the BTZ-treated and non-treated control values was calculated for each gene individually using the unpaired two-sided t-test (p value < 0.05, n=4) with applied Holm-Šidák correction (*α=5%) in GraphPad Prism 6 software.

### TABLES

**Table S1:** TALEN sequences used for generation of the *C57BL/6NCrl-Ddi2*$^{em1}$*/Ph* strain.

| Intron | site | TALEN sequence |
|---|---|---|
| 5 | 5´ | HD NG NG HD NI HD NG NN NN NN NN HD NI NN HD NN NG |
| 5 | 3´ | HD HD NI HD HD NI NI HD NI NN NI NI NI NI NG |
| 6 | 5´ | NN NG NN NG HD HD NG NG NN NG NN NG NI HD NN NN NN |
| 6 | 3´ | HD HD HD HD NI NN NG NN HD NG NN HD HD HD NG HD NG NN |

**Table S2:** Primer sequences used for off-target screening in the *C57BL/6NCrl-Ddi2*$^{em1}$*/Ph* strain generated by TALEN-mediated gene alternation.

| Primer name | Sequence |
|---|---|
| Ddi2_OT_1_F | 5´-TCCCTTTCATGAGGCCATTC-3´ |
| Ddi2_OT_1_R | 5´-AGCGCAGAGAATGAAAAAGC-3´ |
| Ddi2_OT_2_F | 5´-TGCTGAATTAGTGCTTTCATGTGG-3´ |
| Ddi2_OT_2_R | 5´-TACCATGCACACGCATCTCA-3´ |
| Ddi2_OT_3_F | 5´-TTCTTTCCAACTAACCCACA-3´ |
| Ddi2_OT_3_R | 5´-CTGGGATGAGAAGTTTTGAG-3´ |
| Ddi2_OT_4_F | 5´-TGACCAATGTAGTGGATAG-3´ |
| Ddi2_OT_4_R | 5´-TGGTGGATGTCAAGGATTAT-3´ |
| Ddi2_OT_5_F | 5´-GAACCTGAGTCTTCTGCAA-3´ |
| Ddi2_OT_5_R | 5´-AAGCACTCTACTGCTTTCC-3´ |
| Ddi2_OT_6_F | 5´-GAGGAACCACCTAGGGCTGA-3´ |
| Ddi2_OT_6_R | 5´-CAGGTCAGAGATGGGTCTGC-3´ |
| Ddi2_OT_7_F | 5´-CTGGACACTGGCTCTTC-3´ |
| Ddi2_OT_7_R | 5´-CGCAGTAGAAACATTGCAA-3´ |
| Ddi2_OT_8_F | 5´-AGCATGGGTACCAATTCCAGA-3´ |
| Ddi2_OT_8_R | 5´-TGCACCATGTAGACATTGACG-3´ |
| Ddi2_OT_9_F | 5´-TCTCCCTTGCCCCTTTAG-3´ |
| Ddi2_OT_9_R | 5´-TAATGGGGGAGTAGGACAGT-3´ |
| Ddi2_OT_10_F | 5´-TATAAGCCTGGCCTTTCTTGT-3´ |
| Ddi2_OT_10_R | 5´-TGTGCTCTCACACCCAC-3´ |
| Ddi2_OT_11_F | 5´-GTTGCAGCTCACCTTGAACG-3´ |
| Ddi2_OT_11_R | 5´-TTTGCCAGTCTCAGGTTGCT-3´ |
| Ddi2_OT_12_F | 5´-TCTGCTGCATTGTTTTATTGC-3´ |
| Ddi2_OT_12_R | 5´-CACAGGAACTTCTGGTGACTT-3´ |

**Table S3:** Primer sequences used for genotyping of both *Ddi2*$^{KO}$ and *Ddi2*$^{PD}$ strains.

| Primer name | Sequence | Use |
|---|---|---|
| Ddi2F | 5´-GTCTGGTCCTTGTCCGTGTT-3´ | genotyping *Ddi2*$^{PD}$ |
| Ddi2R | 5´-AGTCTGTCATCCCGAGTTGG-3´ | genotyping *Ddi2*$^{PD}$ |
| Ddi2 nested F | 5´-GTGAGACCCTGACTCGGCAA-3´ | genotyping *Ddi2*$^{PD}$ |
| Ddi2 long R | 5´-CCTGGCAACCTGAAATCAAG-3´ | genotyping *Ddi2*$^{PD}$ |
| Ddi2 IN R | 5´-GACTGTAAAACATAAGCCAC-3´ | genotyping *Ddi2*$^{PD}$ |
| Ddi2tm1b WT F | 5´-GCATGGGCTTACAGTGGTTACTC-3´ | genotyping *Ddi2*$^{KO}$ |
| Ddi2tm1b RV | 5´-CTTACTAGTTGCACAGCTGATGACATC-3´ | genotyping *Ddi2*$^{KO}$ |
| LacZ_R | 5´-ACGGTTTCCATATGGGGATT-3´ | genotyping *Ddi2*$^{KO}$ |

**Table S4:** Sequences of primer pairs used in qRT-PCR analysis of gene expression (listed in alphabetical order).

| Gene | forward primer | reverse primer |
|---|---|---|
| *Atf4* | CGGCAAGGAGGATGCCTT | TGGTTTCCAGGTCATCCATT |
| *Bcl2* | ATCGCCCTGTGGATGACTGA | ACAGCCAGGAGAAATCAAACAGA |
| *Bip* | GTTTGTCCCCTTACACTTGG | GTCGTTCACCTTCATAGACC |
| *Chop* | GAACCTGAGGAGAGAGTGTT | TATAGGTGCCCCCAATTTCA |
| *Cxcl10* | GTGTTGAGATCATTGCCACG | AAGGAGCCCTTTTAGACCTT |
| *Ddi1* | ACTACCGGCTCACAGACTCA | GACAGGGTGTTTGATAGTACCTCC |
| *Ddi2* | CACACAGAAGATTATTGGAAGG | CGTTTCAGCATGTCCAGACC |
| *Ddi2 Δex6* | TTGTCGACTCAGCTCAGGTTCA | AGCATGTCCATAGGCTGTTCTT |
| *Gclm* | TCCCGATGAAAGAGAAGAAATGAA | GTGCAACTCCAAGGACGGA |
| *Grp94* | TATGGATGGTCTGGCAACAT | TTTAAACTGAGGCGAAGCAT |
| *Gss* | AAGATGTCTCTGAAAGGGGTTCT | AGGCGTGCTTCCCAGTT |
| *H2afz* | TAGGACAACCAGCCACGGA | GACGAGGGGTGATACGCTTT |
| *Herpud1* | CATCCTTTACTTCTACTCCTCGCT | TCTGTCTGAACGGAAACCACC |
| *Hmox1* | AGGGTGACAGAAGAGGCTAAGA | AGCTAGTGCTGATCTGGGGTT |
| *Ifi44* | ACAGATACCAGTTCGATTCCA | CACGTGTGTAAGTAAAGCCA |
| *iNos* | CGGACGAGACGGATAGGCA | CGTGGGGTTGTTGCTGAACT |
| *Ngly1* | TCAGGAAACCAAACAGGGCAG | ATGCTGAATGTTGGACTGAAGAAC |
| *Nqo1* | AAGAGCTTTAGGGTCGTCTTGG | CATGGCGTAGTTGAATGATGTCT |
| *Nrf1* | GGGTGCGGAAGCTCTGG | CGGGGGACTCACTCTCACT |
| *Nrf2* | CAGCACATCCAGACAGACACC | ACTCATGGTCATCTACAAATGGGAA |
| *Nrf3* | TGCCAGATGCAGGCGGA | CTGACACCCCTTCCTCGTTT |
| *Psma4* | GATCTGCACCCTCACCGTC | CCACTTGGTATAAGCGACCTTC |
| *Psma6* | AAGTACCCGACAAACTACTGGA | ATAGCGTGCCCTCTGTACCT |
| *Psmb6* | GTCTCCACAGGGACCACGA | TGAGCGGCAGCAGAAGATG |
| *Rad23A* | CGCATGGAACCTGACGAGAC | AGCCACGGGGAAAGCATC |
| *Rad23B* | AGAAAAGCCAGCCCAGACA | AGGGCACTTGTTGCATCTT |

| | | |
|---|---|---|
| *Rpl19* | CAGATAATGGGCGGAGCCTG | GGCAAGCCTCTTCTGTAGCC |
| *Rsc1a1* | GCCTGCTTTTATTCTATTGAGGATT | TGACATTCCCAGTGAGGTAGATT |
| *Sprtn* | CATGTGCTCCATCCGTCTCA | CGTGTCCTTCCCGGTCTTT |
| *Tbp* | TATCTACCGTGAATCTTGGCTG | TTGTCCGTGGCTCTCTTATTCT |
| *Ubqln1* | TGATGGACTTACGGTTCACCTTGT | GTGCTTCCAGGGGCATTTGT |
| *Ubqln2* | CACCTACCACCACGAATAGCA | TGAAGCTCGGTGAAGTTGGG |
| *Usp18* | ATTGAAGAGGATAACAGTGCC | CGTGATCTGGTCCTTAGTCA |
| *Vcp/p97* | TGAGACATCCTGCGCTCTTT | AGGCTCCAGTTTCATTTGCC |
| *Xpc* | GGGTATTGTCGTGGAGAAGCA | GGGCACGGTTAGAGAAGCC |

**Table S5:** Sequence of the *Ddi2* antisense probe used in ISH studies.

| Probe | Sequence |
|---|---|
| Ddi2 AS | 5'- TGCAGCACACACACACTACATGCATCATGGCTTCTGACGCTCTGCATCCTCGGCTGATTTTTGAATG GCTTCTGCTAATTCTTGGTCTGCAATTTCCTCTGGCCGTATGTCCTCTCGCCCGGTTCCATATGCCAAAC GAGCACACTCTGGTAACTCCCCCTCAGGAAGGAAGGTGGTCTGTGAGCCCGTGGTGCCAATCACCAG AACATTTTTCTTCAAGTCAATAGAACACTGGTGCCGTTTCAGCATGTCCAGACCCAGAAGCATGTCCAT AGGCTGTTCTTCCAGAATTGAGAAGGAGCACGCCAGAAAATCGCCTTCAATCTGAACCTGAGCTAGAT GCACCCTTCCAATAATCTTCTGTGTGCCTACTCCTTTGGCAATGCCAGCCCACCGACGATCTACCAGTCT CATTATGTTACACCTTTCTGCACAGGCCTGGCTCATAATAGTCATCTGGGCTCCTGAGTCGACAAAGGC TTTCACAGGATGCCCGTTCACTCTGCAGTTGATGTACAGCATGGCGACCTGGCCGAAGCTCTCCGGGG CCTCCTCCATAGCTATCGTCATGTTCTCCTCGATGTTCTGCTGCCTTATATCTTCTTCTATCTTTGCCTGA GCTTCGAGATCAAAGGGGTCAGCAGAAAACAGACGAATCCGTTCTTGTTCTCTCCGGGCTCGGTCCTG CTGCTGCTCCACCAAGACCCTAGAGAATTTCTCAAGATCTCCACTGAGCAGAGCTTCCGCCAAGGGTG GGTTGCGCTCCTTCAGCAAGGACAACTCATGCGGGTTGGCCAGCAGCATGTCTCGGAGCAGGGCTGG GTTGTCTAGGCCTTGAGGAGAAGATGCCATTTCCCCAGGAGATGAGTGCTGAGCCTGGGTTCTGGGC AGCTGGCGCTGTTGGGGGTTTGACGTGCCAGGCACAGCTATGCTACTGAAGTCAATCCGGGGTAAGT TTGAGAACTGCACTGCAGGGCGAGGGTCTGCATTCTCCTTCTGTCG - 3' |
